# A murine model of hnRNPH2-related neurodevelopmental disorder recapitulates clinical features of human disease and reveals a mechanism for genetic compensation of *HNRNPH2*

**DOI:** 10.1101/2022.03.17.484791

**Authors:** Ane Korff, Xiaojing Yang, Kevin O’Donovan, Abner Gonzalez, Brett J. W. Teubner, Haruko Nakamura, James Messing, Fen Yang, Alex Carisey, Yong-Dong Wang, Tushar Patni, Heather Tillman, Stanislav S. Zakharenko, Yuh Min Chook, J. Paul Taylor, Hong Joo Kim

## Abstract

Mutations in *HNRNPH2* cause an X-linked neurodevelopmental disorder with a phenotypic spectrum that includes developmental delay, intellectual disability, language impairment, motor function deficits, and seizures. More than 90% of patients with this disorder have a missense mutation within or adjacent to the nuclear localization signal (NLS) of hnRNPH2, although the specific pathogenic consequences of these mutations have not been examined. Here we found that hnRNPH2 NLS mutations result in reduced interaction with the nuclear transport receptor Kapβ2 in vitro and in cultured human cells. These mutations also cause modest accumulation of hnRNPH2 in the cytoplasm, suggesting that mislocalization of the protein might contribute to pathogenesis. We generated two knock-in mouse models with human-equivalent mutations in the endogenous mouse gene *Hnrnph2*, as well as *Hnrnph2* knockout (KO) mice, and subjected them to extensive phenotyping. Mutant knock-in mice displayed a spectrum of phenotypes that recapitulated aspects of the human disorder, including reduced survival in males, craniofacial abnormalities, impaired motor and cognitive functions, and increased susceptibility to audiogenic seizures. Mutant knock-in male mice developed more severe phenotypes than female mice, likely due to differences in X-chromosome gene dosage. In contrast, two independent lines of *Hnrnph2* KO mice showed no detectable phenotypes. Notably, KO mice had upregulated expression of *Hnrnph1*, a close paralog of *Hnrnph2*, whereas mutant *Hnrnph2* knock-in mice failed to upregulate *Hnrnph1.* Thus, genetic compensation by *Hnrnph1* might be sufficient to counteract the loss of hnRNPH2. These findings suggest that the pathogenesis of *HNRNPH2*-related disorder in humans may be driven by a toxic gain of function or a complex loss of *HNRNPH2* function with impaired compensation by *HNRNPH1.* The mutant knock-in mice described here are an important resource for preclinical studies to assess the potential benefit of either gene replacement or therapeutic knockdown of mutant hnRNPH2.

## Introduction

De novo pathogenic variants in *HNRNPH2* were identified in 2016 in six unrelated individuals as a novel cause of an X-linked neurodevelopmental disorder whose features include developmental delay, intellectual disability, autism spectrum disorder, tone abnormalities, and seizure (OMIM 300986) (1). Since the initial identification of these mutations, the genotypic and phenotypic spectrum of the disorder has been expanded to include more than 30 individuals with 11 distinct de novo variants (2), as well as several maternally inherited cases (3–5). Although all six individuals in the initial report were female, subsequent studies have identified males carrying missense mutations in *HNRNPH2* associated with a range of overlapping phenotypes (5–7).

More than 90% of individuals with *HNRNPH2*-related neurodevelopmental disorder have a nonsynonymous single nucleotide variant within or adjacent to the nuclear localization signal (NLS) of hnRNPH2, with the two most common missense variants, R206W and R206Q, located within the NLS. Additional variants outside the NLS of hnRNPH2 have been reported in children with similar symptoms. Two of these, at residues 114 and 188, are recurrent, suggesting a potential pathogenic effect (4), whereas additional variants found in single patients are of less clear significance. Notably, individuals with NLS mutations have more severe symptoms than those with variants located outside this region, the latter of which have been reported almost exclusively in males (2, 4, 8).

Rare pathogenic variants in *HNRNPH1*, a close paralog of *HNRNPH2*, have been reported in patients with a syndrome very similar to that observed in *HNRNPH2*-related disorder (9, 10). Half of these variants (4 of 8) are located in the NLS of hnRNPH1. As with mutations in hnRNPH2, patients harboring variants within the NLS of hnRNPH1 display a much more severe phenotype than patients whose variants are located outside the NLS (9, 10). These observations suggest the possibility of a common basis for abnormal neurodevelopment related to impairment of functions shared between hnRNPH1 and hnRNPH2.

hnRNPH2 is a member of the heterogeneous nuclear ribonucleoprotein (hnRNP) family of proteins, which govern various aspects of nucleic acid metabolism including transcription, RNA processing, alternative splicing, mRNA trafficking, mRNA stability, and translation (11). hnRNPH2 is a member of the hnRNP H/F subfamily, which comprises hnRNPH1, hnRNPH2, hnRNPH3, and hnRNPF. As components of a messenger ribonucleoprotein (mRNP) complex, hnRNP H/F proteins shuttle between the nucleus and the cytoplasm and function in both cellular compartments. Nucleocytoplasmic transport of hnRNP F/H proteins is regulated by their proline-tyrosine NLS (PY-NLS), which is located in the center of the protein flanked by two RNA-recognition motifs (RRMs) (12). In humans, PY-NLSs are recognized for nuclear import by karyopherin β2 (Kapβ2) (13). Deletion of the PY-NLS domain in hnRNPF or mutation of the conserved PY-NLS motif in hnRNPH1 impairs nuclear localization of these proteins (12, 13).

Pathogenic mechanisms arising from variants in *HNRNPH2* remain largely unexamined. One recent in vitro study showed deficiencies in nucleocytoplasmic shuttling of hnRNPH2 with NLS mutations (R206W, P209L), as well as alterations in splicing associated with hnRNPH2 R114W (4). However, detailed characterizations of pathogenic mutations have not been reported, nor have faithful models that recapitulate features of the human clinical syndrome.

Mechanistic insight into the functional consequences of syndrome-causing mutations and robust, disease-relevant models are essential for therapeutics development.

Here we investigated the consequences of common pathogenic mutations in *HNRNPH2,* focusing on variants within the PY-NLS. Mutant proteins showed reduced interaction with Kapβ2 in vitro and in human cells, and modest, but abnormal, accumulation in the cytoplasm when expressed in human cells. Knock-in mice bearing two distinct pathogenic NLS mutations in *Hnrnph2* demonstrated a phenotypic spectrum highly similar to clinical features observed in human patients, including reduced survival in males, craniofacial abnormalities, impaired motor and cognitive functions, and increased susceptibility to audiogenic seizures. In contrast, two independent *Hnrnph2* knockout (KO) mice showed no detectable phenotypes, arguing against a simple loss of hnRNPH2 function as the primary driver of disease. RNA-seq analyses revealed that pathogenic mutations induce more severe transcriptome changes than KO both in human iPSC-derived neurons and in mice. Importantly, KO of *HNRNPH2/Hnrnph2* was associated with significant upregulation of the paralogous genes *HNRNPH1/Hnrnph1*, whereas knock-in of pathogenic mutations did not result in such upregulation. Thus, our data suggests the possibility of a toxic gain of function or a complex loss of function of hnRNPH2 driven by a failure in compensation by *Hnrnph1*. This study advances a pathogenic mechanism for *HNRNPH2*- related disorder, suggests a mechanism for genetic compensation of *HNRNPH2*, and provides valuable models for potential use in preclinical studies.

## Results

### Pathogenic variants alter the nucleocytoplasmic ratio of hnRNPH2 and enhance its recruitment to stress granules

Most hnRNPH2 mutations associated with neurodevelopmental phenotypes are single nucleotide variants located in its PY-NLS, which comprises a central hydrophobic or basic motif followed by the motif R/H/K-X2−5-PY (**Figure 1, A and B**). To examine the impact of disease-causing mutations on the subcellular localization of hnRNPH2, we expressed epitope-tagged wild-type (WT) and variant (R206W, R206Q, and P209L) forms of hnRNPH2 in HeLa cells. Under basal conditions, hnRNPH2 WT was almost exclusively located in the nucleus, consistent with established roles for this protein in nuclear RNA processing steps such as splicing. In contrast, disease-causing variants were found in both the nucleus and cytoplasm (**Figure 1, C and D**). The cytoplasmic accumulation of mutant proteins became more evident when cells were exposed to oxidative stress (0.5 mM NaAsO2), which induced the assembly of cytoplasmic stress granules as marked by eIF3η staining (**Figure 1, C and D**). All disease-causing mutant forms of hnRNPH2, but not hnRNPH2 WT, were associated with stress granules (**Figure 1, C and D**). These differences in stress granule association occurred despite comparable levels of hnRNPH2 expression across WT and mutant forms (**Figure 1E**), suggesting that cytoplasmic accumulation and recruitment of mutant hnRNPH2 to stress granules are not attributable to overexpression of protein. This result is also consistent with previous reports that mutations in the PY-NLS of other hnRNPs interfere with their nuclear import and enhance their incorporation into stress granules (14–16).

**Figure 1.**
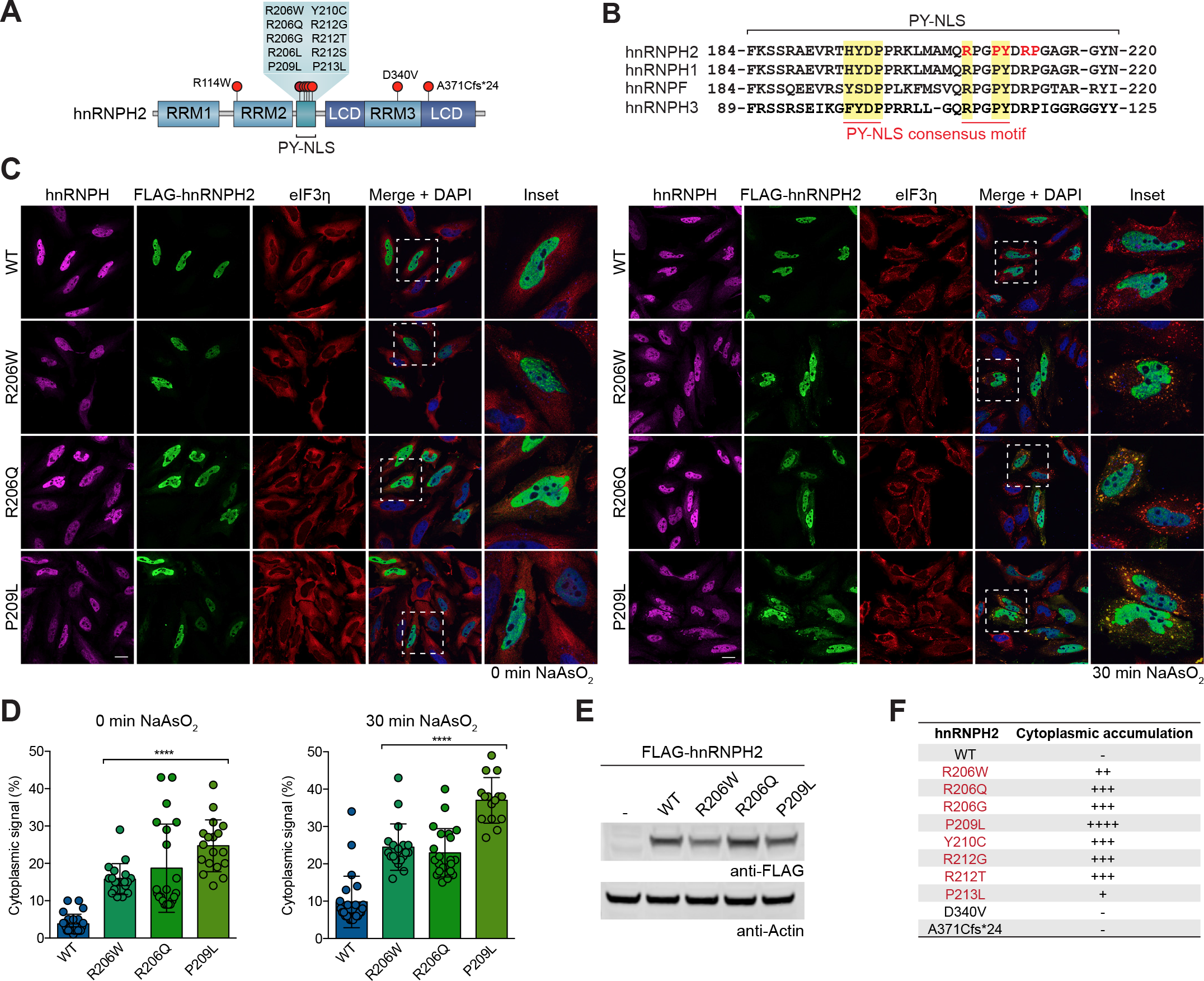
Pathogenic variants alter the nucleocytoplasmic ratio of hnRNPH2 and enhance its recruitment to RNP granules. (**A**) Schematic representation of hnRNPH2 structure, including mutations identified in patients with a neurodevelopmental disorder. hnRNPH2 contains RNA recognition motifs 1, 2, and 3 (RRM1, RRM2, RRM3), and a nuclear localization signal (PY-NLS). (**B**) Sequence alignment of four human paralogs of the hnRNP F/H family showing high conservation of disease-affected and surrounding residues. Consensus PY-NLS motifs are highlighted in yellow. Amino acids mutated in hnRNPH2 in patients are in red. (**C**) Intracellular localization of FLAG-tagged hnRNPH2 WT, R206W, R206Q, and P209L mutants under basal (left) and stressed (right) conditions in HeLa cells. hnRNPH antibody was used to show endogenous hnRNPH1 and hnRNPH2 localization patterns in untransfected cells. eIF3η was used as a cytoplasmic and stress granule marker. Scale bar, 10 μm. (**D**) Quantification of hnRNPH2 cytoplasmic signal intensity in HeLa cells as shown in (**C**). An interleaved scatter plot with individual data points is shown; error bars represent mean ± SD. For WT, R206W, R206Q, and P209L mutants, n = 24, 19, 21, and 18 cells for basal conditions, n = 24, 19, 22, and 15 cells for stressed conditions, respectively. *\*\*\*\*P* < 0.0001 by one-way ANOVA with Dunnett’s multiple comparisons test. (**E**) Immunoblot showing comparable levels of expression between hnRNPH2 WT, R206W, R206Q, and P209L mutants. Representative images are shown from n ≥ 3 experiments. (**F**) Summary of intracellular localization of FLAG-tagged hnRNPH2 WT and mutants in HeLa cells, with or without arsenite stress. Images are shown in Supplemental Figure 1. Proteins with PY-NLS mutations (red font) showed cytoplasmic accumulation.

We next characterized the subcellular localization of 7 additional variants, 5 of which alter the amino acid sequence within the PY-NLS (R206G, Y210C, R212G, R212T, P213L). The remaining two variants alter the amino acid sequence in RRM3 (D340V) or the C-terminal low complexity domain (LCD; A371Cfs*24), respectively. All hnRNPH2 variants within the PY-NLS showed accumulation of hnRNPH2 protein in the cytoplasm, although the levels of accumulation varied (**Figure 1F** and **Supplemental Figure 1**). Importantly, despite mutation-dependent redistribution to the cytoplasm, the majority of hnRNPH2 was still found in the nucleus. Indeed, even with the mutation (P209L) that caused the greatest amount of cytoplasmic accumulation, we estimated that ∼75% of hnRNPH2 was found in the nucleus. In contrast, the two non-PY-NLS variants did not show cytoplasmic accumulation of hnRNPH2 (**Figure 1F** and **Supplemental Figure 1**); thus, cytoplasmic redistribution of hnRNPH2 is not an invariant feature of the neurodevelopmental syndrome. We note that these two non-PY-NLS variants were found in male patients, who are hemizygous and therefore express only mutant hnRNPH2, in contrast to heterozygous female patients who express a mix of WT and mutant protein.

#### Pathogenic variants impair the interaction between hnRNPH2 and its nuclear transport receptor Kapβ2

Closely paralogous proteins hnRNPH1 and hnRNPF interact with the nuclear import receptor Kapβ2 via their PY-NLS (12, 13, 17). Given the high degree of identity among hnRNPH1, hnRNPF, and hnRNPH2 (**Figure 1B**), we predicted that hnRNPH2 would bind to Kapβ2 via its PY-NLS for nuclear import and that disease-causing mutations in the PY-NLS would alter this interaction. To test this hypothesis, we performed GST pulldown assays of GST-tagged WT and mutant (R206W, R206Q, and P209L) versions of the hnRNPH2 PY-NLS (aa 179-215) (**Supplemental Figure 2, A and B**). As a positive control, we used an M9M peptide designed to bind to the PY-NLS binding site of Kapβ2 with an affinity that is ∼200-fold stronger than a natural PY-NLS (18). We observed pulldown of Kapβ2 with GST-M9M and GST- hnRNPH2 WT peptide, but not with mutant peptides (**Supplemental Figure 2B**). When we increased the length of the N- and C-terminal flanking sequences included in the PY-NLS peptide (aa 169-225), the interaction between Kapβ2 and GST-hnRNPH2 PY-NLS peptide was greatly enhanced (**Supplemental Figure 2C**). Indeed, this form of GST-hnRNPH2 PY-NLS WT (aa 169-225) bound Kapβ2 as efficiently as GST-M9M (**Supplemental Figure 2C**). However, the disease-associated mutant peptides showed reduced interaction with Kapβ2 even when expressed with this larger flanking sequence (**Figure 2, A and B** and **Supplemental Figure 2C**). The degree to which the PY-NLS mutations impaired Kapβ2 binding correlated with the degree of cytoplasmic redistribution observed in cells, with P209L having the greatest effect (**Figure 1, C–F** and **Figure 2, A and B**).

**Figure 2.**
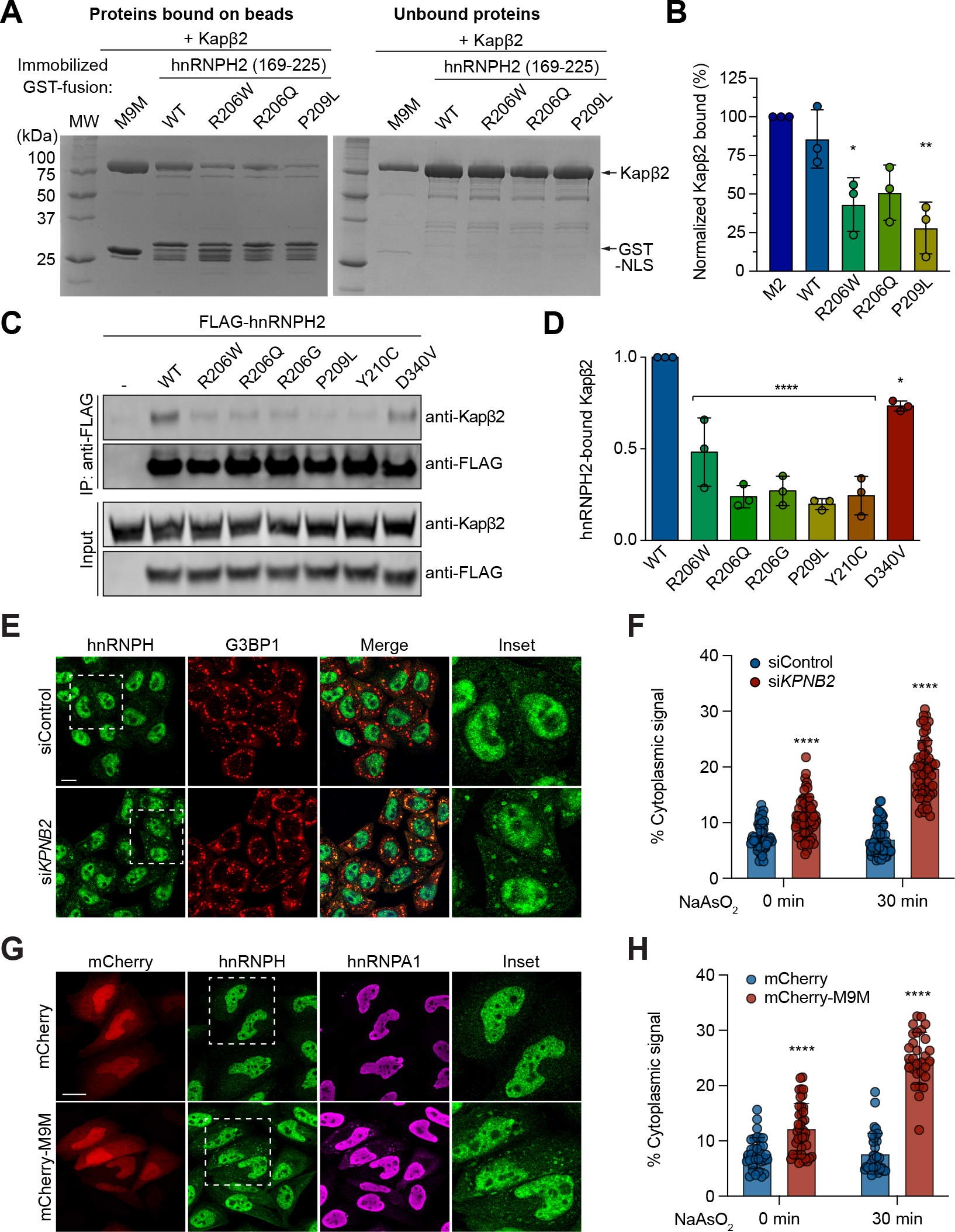
Pathogenic variants impair the interaction between hnRNPH2 and its nuclear transport receptor Kapβ2. (**A**) GST pulldown of purified GST-hnRNPH2 peptides with Kapβ2. Proteins bound (left) and unbound (right) to beads are shown. Proteins were visualized by Coomassie Blue staining. (**B**) Graph showing relative band intensities of bound Kapβ2 in triplicate experiments. Error bars represent mean ± SD, n = 3 biological repeats. *\*P* < 0.0263 and *\*\*P* = 0.0042 by one-way ANOVA with Dunnett’s multiple comparisons test. (**c**) HEK293T cells were transfected with indicated FLAG-hnRNPH2 constructs, immunoprecipitated for FLAG, and immunoblotted. Kapβ2 binding was reduced in hnRNPH2 variants with PY-NLS mutations. (**D**) Quantification of hnRNPH2 and Kapβ2 interaction using densitometry from immunoblots as shown in (**C**). Error bars represent mean ± SD, n = 3 biological repeats. *\*\*\*\*P* < 0.0001 and *\*P* = 0.0138 by one-way ANOVA with Dunnett’s multiple comparisons test. (**E**) Fluorescent staining of HeLa cells transfected with non-targeting siRNA (siControl) or siRNA targeting *KPNB2*/*TNPO1*. Forty-eight hours post-transfection, cells were treated with 0.5 mM NaAsO2 for 30 min, fixed, and stained with indicated antibodies. G3BP1 was used as a stress granule marker. Scale bar, 10 μm. (**F**) Quantification of hnRNPH2 cytoplasmic signal intensity in HeLa cells as shown in (**E**). An interleaved scatter plot with individual data points is shown; error bars represent mean ± SD. For siControl and si*KPNB2*, n = 56 and 62 cells for basal conditions, n = 63 and 57 cells for stressed conditions, respectively. *\*\*\*\*P* < 0.0001 by two-way ANOVA with Sidak’s multiple comparisons test. (**G**) Fluorescent staining of HeLa cells transfected with mCherry or mCherry-M9M. Twenty-four hours post-transfection, cells were fixed and stained with indicated antibodies. hnRNPA1 was used as a positive control for mCherry-M9M. Scale bar, 10 μm. (**H**) Quantification of hnRNPH2 cytoplasmic signal intensity in HeLa cells as shown in (**G**). An interleaved scatter plot with individual data points is shown; error bars represent mean ± SD. For mCherry and mCherry-M9M, n = 34 and 40 cells for basal conditions, n = 33 and 29 cells for stressed conditions, respectively. *\*\*\*\*P* < 0.0001 by two-way ANOVA with Sidak’s multiple comparisons test.

We next expressed full-length hnRNPH2 constructs in cells to test their interaction with Kapβ2 via immunoprecipitation. Consistent with our GST pulldowns, full-length hnRNPH2 WT co-immunoprecipitated efficiently with Kapβ2, whereas all disease-associated mutant proteins showed reduced interaction with Kapβ2 (**Figure 2, C and D**). Notably, PY-NLS mutants (R206W/Q/G, P209L, and Y210C) showed a far greater reduction in Kapβ2 interaction (∼50- 75%) compared with the non-PY-NLS mutant D340V (**Figure 2, C and D**).

To test the functional consequences of the hnRNPH2-Kapβ2 interaction, we next inhibited Kapβ2 by RNAi-mediated knockdown (**Supplemental Figure 2, D and E**). Expression of siRNA targeting *KPNB2* (also known as *TNPO1*) resulted in a ∼90% decrease in Kapβ2 protein levels (**Supplemental Figure 2, D and E**) accompanied by increased cytoplasmic localization of endogenous hnRNPH protein and increased association of hnRNPH with stress granules as assessed by staining with the stress granule marker G3BP1 (**Figure 2, E and F**). We observed a similar result with overexpression of mCherry-M9M peptide, which caused both endogenous hnRNPH and hnRNPA1 to accumulate in the cytoplasm and form cytoplasmic puncta (**Figure 2, G and H**). These results support the hypothesis that disease-associated PY-NLS mutations impair the ability of hnRNPH2 to bind Kapβ2, thereby diminishing nuclear import of hnRNPH2 and leading to cytoplasmic accumulation.

### hnRNPH2 P209L and R206W mice, but not KO mice, have reduced survival and body weight

Like human *HNRNPH2*, the mouse *Hnrnph2* gene is located on the X chromosome. Human hnRNPH2 and mouse hnRNPH2 have 99% identity; both are 449 amino acids in length, with conservative differences in the identity of only 4 amino acids (<1%), and the PY-NLS motif is perfectly conserved between the two species (**Supplemental Figure 3**). To investigate the effects of mutations on normal hnRNPH2 function and to model the pathogenicity of mutant hnRNPH2, we generated mouse models by homologous knock-in of the human hnRNPH2 R206W or P209L mutations into mouse *Hnrnph2*. To this end, we substituted two conserved C nucleotides at positions 833 and 835 with T and G, respectively (c.833 C > T and c.835 C > G) for the R206W mutation, and the C nucleotide at position 842 with T (c.842 C > T) for the P209L mutation (**Figure 3, A and B** and **Supplemental Figure 4A**). While generating these knock-in mouse lines, we also serendipitously obtained a KO line in which a frameshift caused by an indel generated a premature stop codon, leading to nonsense-mediated mRNA decay (**Figure 3B** and **Supplemental Figure 4, B–D**). We also obtained a second KO line from the Knockout Mouse Project (KOMP) Repository (C57BL/6NJ-*^Hnrnph2em1(IMPC)J^*/MmJax, The Jackson Laboratory). These two KO lines differ in two respects: first, in contrast to our KO line, which is driven by an indel with consequent nonsense-mediated decay, the KOMP KO line was generated by a 1451-bp deletion in exon 4, which is predicted to result in a truncated, non-functional transcript. Second, the background strain of our *Hnrnph2* mutant and KO lines is C57BL/6J, whereas the KOMP KO is on a C57BL/6NJ background. Thus, we selected two distinct disease-associated *Hnrnph2* knock-in mouse lines (R206W and P209L) and two distinct *Hnrnph2* KO lines for in-depth phenotypic analysis. Importantly, extensive phenotypic analysis of our indel-based KO line and the KOMP KO line yielded equivalent results. For ease of presentation, we include data from our own indel-based KO line (hereafter referred to as KO) in all subsequent figures alongside data from our knock-in lines, whereas all results from the KOMP KO line are compiled in Supplemental Figure 5.

**Figure 3.**
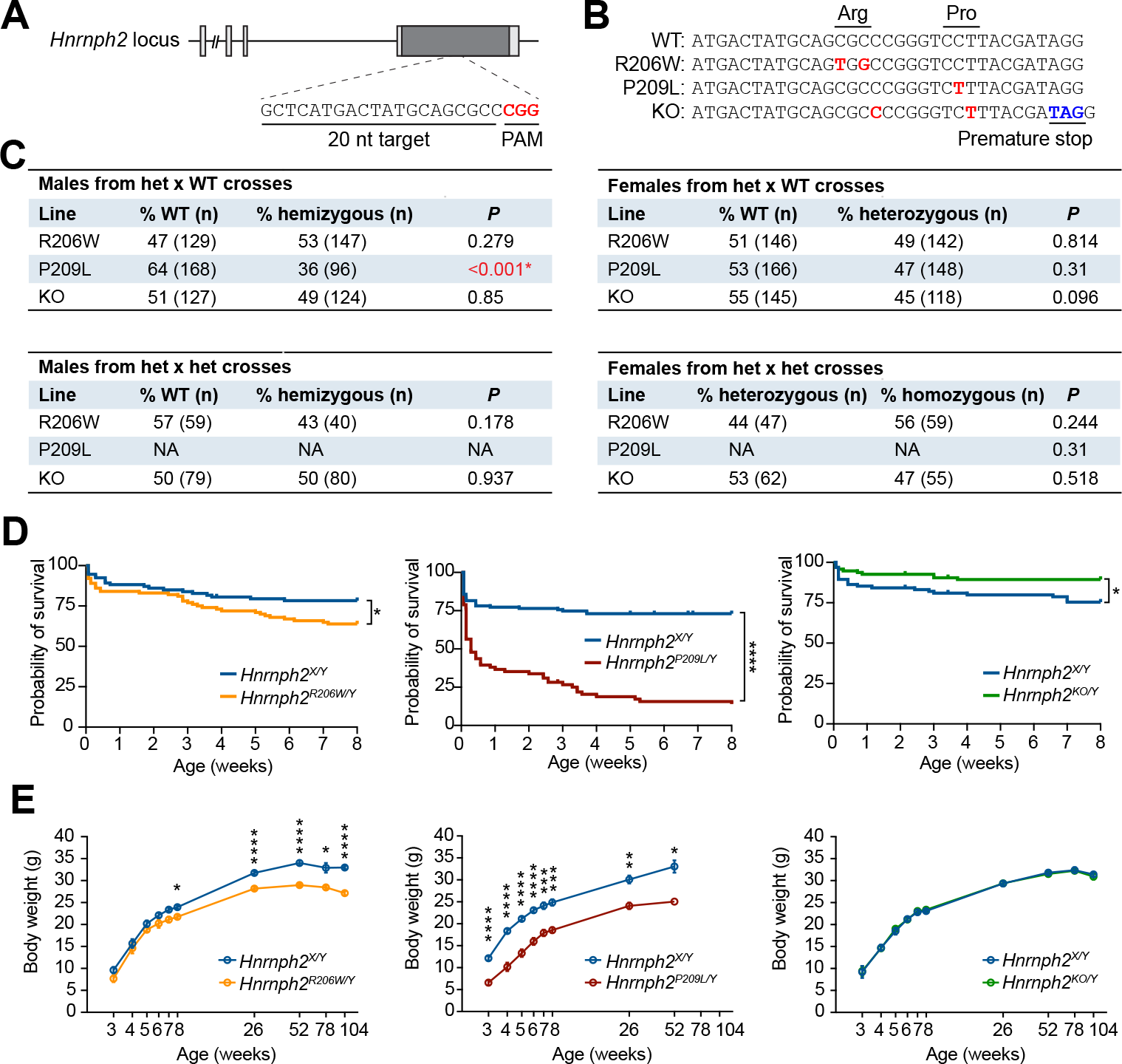
Generation, survival, and body weight of *Hnrnph2* mutant and KO mice. (**A**) Schematic of the mouse *Hnrnph2* locus. sgRNA target sequence is shown. Red text indicates protospacer adjacent motif (PAM). (**B**) Nucleotide sequences showing types of editing induced by the sgRNA and single-stranded oligo donor in mouse. Red text indicates edited nucleotide sequences; blue text indicates a premature stop codon introduced by an indel. (**C**) Ratios of genotyped mice organized by sex and breeding strategy. Significant *P* values shown in red (Chi- square test), NA; not applicable. (**D**) Kaplan-Meier survival curves of male mice up to 8 weeks of age, \**P* = 0.0371, hazard ratio (HR) = 1.764 for *Hnrnph2^R206W/Y^* (n = 100) vs. *Hnrnph2^X/Y^*(n = 93); \*\*\*\**P* < 0.0001, HR = 4.363 for *Hnrnph2^P209L/Y^* (n = 71) vs. *Hnrnph2^X/Y^* (n = 119); \**P* = 0.0144, HR = 0.4111 for *Hnrnph2^KO/Y^* (n = 95) vs. *Hnrnph2^X/Y^* (n = 95) by Mantel-Cox test. (**E**) Mean body weight of male mice over time. Error bars represent mean ± SEM. *Hnrnph2^R206W/Y^* (n = 13) vs. *Hnrnph2^X/Y^* (n = 13) \**P* < 0.05 at 8 and 78 weeks, \*\*\*\**P* < 0.0001 at 26, 52, and 104 weeks; *Hnrnph2^P209L/Y^* (n = 12) vs. *Hnrnph2^X/Y^*(n = 7) \*\*\*\**P* < 0.0001 at 3-6 weeks, \*\*\**P* < 0.001 at 7-8 weeks, \*\**P* = 0.0029 at 26 weeks, \**P* = 0.0414 at 52 weeks; ns for *Hnrnph2^KO/Y^* (n = 18) vs. *Hnrnph2^X/Y^* (n = 10), by mixed-effects model (REML) with Sidak’s multiple comparisons test. Two *Hnrnph2* KO lines (KOMP KO and indel KO2) were used for all analyses and both lines showed the same results. For simplicity, data from our indel-based KO line are included in each graph. Data from the KOMP KO line are summarized in Supplemental Figure 5.

All lines produced viable offspring. All heterozygous females (R206W, P209L, and KO) were born with expected frequencies. In contrast, hemizygous P209L mutant males, but not R206W and KO males, were detected at a lower frequency than predicted by Mendelian laws (64% WT vs. 36% *Hnrnph2^P209L/Y^*), suggesting partial embryonic lethality of males bearing the P209L mutation (**Figure 3C**). We also crossed heterozygous mutant females to hemizygous mutant males to produce homozygous mutant mice. Again, homozygous females from R206W and KO lines were born close to expected frequencies (**Figure 3C**). We note that this experiment could not be performed in the P209L line, as most hemizygous P209L males did not survive long enough to breed. Indeed, less than 15% of *Hnrnph2^P209L/Y^* males lived to 8 weeks of age (**Figure 3D**). *Hnrnph2^R206W/Y^*males also had significantly reduced survival up to 8 weeks of age (**Figure 3D**). In contrast, heterozygous females from all lines showed no significant changes in survival up to 8 weeks of age (**Supplemental Figure 6A**), and hemizygous KO males survived slightly better than their littermate controls (**Figure 3D**). Homozygous females from R206W and KO lines also did not have significantly reduced survival up to 8 weeks, although there was a trend toward reduced survival in *Hnrnph2^R206W/R206W^*females compared to *Hnrnph2^R206W/X^* female littermates (**Supplemental Figure 6B**). These results suggest a dosage-dependent effect of *Hnrnph2* mutation on survival and indicate that the P209L mutation has a more severe effect on survival than R206W, consistent with their respective effects in vitro and in cell lines (**Figures 1 and 2**).

To investigate the effects of hnRNPH2 mutations on long-term survival, we monitored a subset of mice for up to 2 years. In this smaller cohort, we found no significant difference in long-term survival between hemizygous males or heterozygous females and WT littermate controls (**Supplemental Figure 6C**). *Hnrnph2^P209L/Y^* and *Hnrnph2^R206W/Y^*males weighed significantly less than their WT littermate controls (**Figure 3E**). *Hnrnph2^R206W/X^* females also had significantly reduced body weight compared to littermate controls (**Supplemental Figure 6D**). *Hnrnph2^P209L/X^*females tended to weigh less than littermates, but the difference was not significant (**Supplemental Figure 6D**). Once more, neither male nor female KO mice were significantly different from controls in body weight or long-term survival, suggesting that reduced survival or reduced body weight in knock-in mice likely does not arise from simple loss of hnRNPH2 function.

### hnRNPH2 P209L and R206W mice, but not KO mice, have craniofacial abnormalities and increased incidence of hydrocephalus

All human patients with hnRNPH2-related phenotypes have dysmorphic facial features including almond-shaped eyes, short palpebral fissures, a short philtrum, full lower lip, long columella, hypoplastic alae nasi, and micrognathia (1). Although most of these patients have unremarkable MRIs, some do present with vertical configuration of the splenium of the corpus callosum, delayed myelination, and decreased cerebellar volume (2). During initial breeding of founders to WT mice, we noticed that in addition to being smaller overall, *Hnrnph2^P209L/Y^* males, and to a lesser extent *Hnrnph2^R206W/Y^* males, appeared to have short snouts and wide-set eyes (**Figure 4A**). To further investigate this phenotype, we performed in vivo µCT and MRI on a cohort of mutant knock-in and KO mice at 6 and 24 weeks of age.

**Figure 4.**
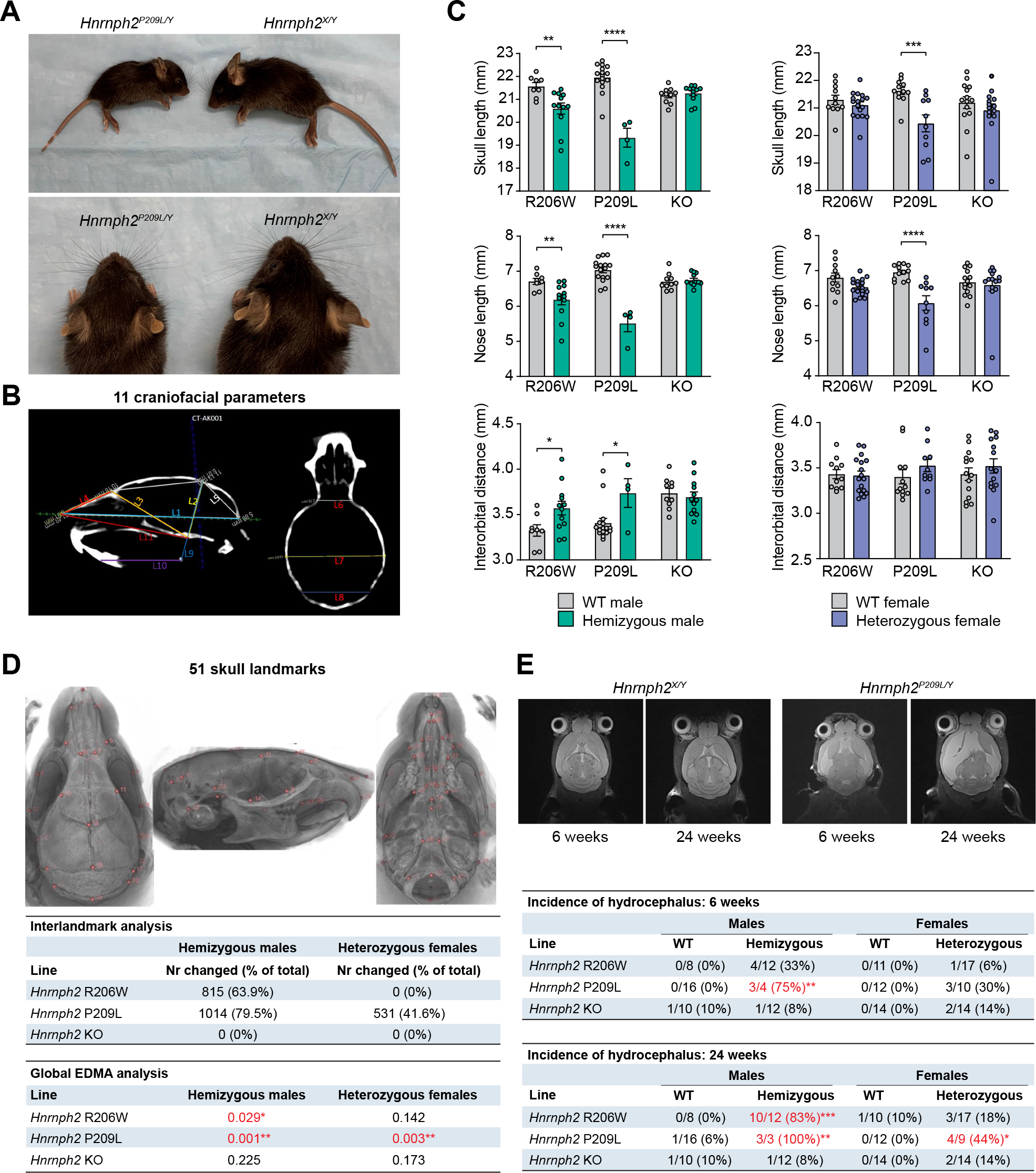
hnRNPH2 mutant mice have craniofacial dysmorphology and an increased incidence of hydrocephalus. (**A**) Representative images of a 3-week-old male *Hnrnph2^P209L/Y^* mouse and WT littermate in lateral and dorsal view. (**B**) Key craniofacial parameters measured manually on individual µCT scans. (**C**) Linear measurements in male and female mice. Error bars represent mean ± SEM. Skull length: \*\**P* = 0.0019 *Hnrnph2^R206W/Y^* vs. *Hnrnph2^X/Y^*; \*\*\*\**P* < 0.0001 *Hnrnph2^P209L/Y^* vs. *Hnrnph2^X/Y^*; \*\*\**P* = 0.0006 *Hnrnph2^P209L/X^* vs. *Hnrnph2^X/X^*. Nose length: \*\**P* = 0.0037 *Hnrnph2^R206W/Y^* vs. *Hnrnph2^X/Y^*; \*\*\*\**P* < 0.0001 *Hnrnph2^P209L/Y^* vs. *Hnrnph2^X/Y^*; \*\*\*\**P* < 0.0001 *Hnrnph2^P209L/X^* vs. *Hnrnph2^X/X^*. Interorbital distance: \**P* = 0.0435 *Hnrnph2^R206W/Y^* vs. *Hnrnph2^X/Y^*; \**P* = 0.0233 *Hnrnph2^P209L/Y^* vs. *Hnrnph2^X/Y^*, by two-way ANOVA with Sidak’s multiple comparisons test. Group sizes for µCT and MRI were as follows: *Hnrnph2^R206W/Y^* (n = 12) vs. *Hnrnph2^X/Y^*(n = 8); *Hnrnph2^P209L/Y^* (n = 4) vs. *Hnrnph2^X/Y^* (n = 16); *Hnrnph2^KO/Y^* (n = 12) vs. *Hnrnph2^X/Y^* (n = 10); *Hnrnph2^R206W/X^* (n = 17) vs. *Hnrnph2^X/X^* (n = 11); *Hnrnph2^P209L/X^* (n = 10) vs. *Hnrnph2^X/X^* (n = 12); *Hnrnph2^KO/X^* (n = 14) vs. *Hnrnph2^X/X^* (n = 14). (**D**) Location of 51 landmarks on mouse skull atlas, number of significantly changed linear interlandmark distances, and results of global EDMA analysis. Significant *P* values are shown in red. (**E**) Representative MRI images showing hydrocephalus in a *Hnrnph2^P209L/Y^*hemizygous male compared to a WT littermate and incidence of hydrocephalus at 6 (\*\**P* = 0.0035) and 24 weeks (\*\*\**P* = 0.0007, \*\**P* = 0.0041, \**P* = 0.0211) of age by Fisher’s exact test.

Manual linear measurements of 11 key craniofacial parameters (19) revealed significant reduction in skull and nose length in *Hnrnph2^P209L/Y^* males, *Hnrnph2^R206W/Y^*males, and *Hnrnph2^P209L/X^* females at 6 weeks of age, as well as a significant increase in interorbital distance in *Hnrnph2^P209L/Y^*and *Hnrnph2^R206W/Y^* males (**Figure 4, B and C**). Furthermore, upper jaw length was significantly reduced in *Hnrnph2^P209L/Y^* males, *Hnrnph2^R206W/Y^*males, and *Hnrnph2^P209L/X^* females, in addition to a reduction in lower jaw length in *Hnrnph2^P209L/Y^* males (**Supplemental Figure 7A**). Importantly, no changes were seen in *Hnrnph2* KO mice (**Figure 4C and Supplemental Figure 7A**). To investigate this craniofacial phenotype more extensively, we used a population-level atlas for the automatic identification of 51 skull landmarks (20), followed by pairwise comparison between all landmarks (**Figure 4D**). After correction for multiple comparisons, *Hnrnph2^P209L/Y^*males, *Hnrnph2^R206W/Y^* males, and *Hnrnph2^P209L/X^*females had many significant changes in inter-landmark distances, mostly decreased compared to littermate controls, whereas KO mice showed no differences (**Figure 4D**). As these measures were not normalized, we were concerned that the observed changes were due to a reduction in the overall size of the mutant mice and not a change in craniofacial shape. To address this, we performed Euclidean distance matrix analysis (EDMA) on 3D landmark coordinate data scaled to centroid size (21, 22). EDMA based on all 51 landmarks revealed a significant difference in global skull shape of *Hnrnph2^P209L/Y^* males, *Hnrnph2^R206W/Y^*males, and *Hnrnph2^P209L/X^* females, but not *Hnrnph2* KO mice (**Figure 4D**). EDMA based on subsets of biologically relevant landmarks (23) showed significant changes in several anatomical regions of *Hnrnph2^P209L/Y^*males, *Hnrnph2^R206W/Y^* males, *Hnrnph2^P209L/X^*females and, to a lesser extent, *Hnrnph2^R206W/X^* females (**Supplemental Figure 7B**). In this analysis, the only changes detected in *Hnrnph2* KO mice were slight but significant differences in the neural crest-mesoderm boundary (**Supplemental Figure 7B**). Results at 24 weeks of age did not differ significantly from those at 6 weeks (data not shown).

hnRNPH2 P209L and R206W mice often had domed heads typically associated with hydrocephalus that develops before ossification of the cranial sutures. Although the C57BL/6J background has a relatively high incidence of hydrocephalus (0.029% at The Jackson Laboratory), the number of mice with pathologically confirmed hydrocephalus suggested an increased incidence in hnRNPH2 P209L and R206W mutant mice, but not in KO mice, compared with WT controls (**Supplemental Figure 7C**). The cause of hydrocephalus in these mice is unknown, as we found neither evidence of aqueduct blockage (**Supplemental Figure 7D**), nor abnormal morphology of cilia on ependymal cells lining the dilated ventricles, nor motile ciliary dysfunction in the respiratory system (data not shown) (24). For a more quantitative measurement of the incidence of hydrocephalus in these lines, mice in the MRI cohort were scored as moderate, high, or severely hydrocephalic. Significantly more 6-week-old *Hnrnph2^P209L/Y^* males, as well as 24-week-old *Hnrnph2^P209L/Y^* males, *Hnrnph2^R206W/Y^*males, and *Hnrnph2^P209L/X^* females had at least moderate hydrocephalus compared to WT littermates (**Figure 4E and Supplemental Figure 7E**). Neither male nor female *Hnrnph2* KO mice showed increased incidence of hydrocephalus compared to WT littermates (**Figure 4E and Supplemental Figure 7C**). Notably, hydrocephalic mice in this cohort did not have obvious doming of the skull, suggesting onset of hydrocephalus after closure of cranial sutures in this group. In agreement with histology, no evidence of aqueduct blockage was detected on MRI (data not shown). Given the MRI abnormalities observed in some patients with hnRNPH2 mutations, we performed automated brain parcellation and volumetrics to investigate group differences in total and regional brain volumes. Automated alignment of MRI images to the DSURQE atlas (25) revealed a small but significant decrease in total brain tissue volume (grey matter plus white matter) in 6-week-old *Hnrnph2^P209L/X^* females, most likely attributable to their small body size. This difference was lost when the CSF volume was added to calculate total intracranial volume (brain tissue volume plus CSF). We did not observe any changes in *Hnrnph2^R206W/Y^*males, *Hnrnph2^R206W/X^* females, *Hnrnph2* KO males, or *Hnrnph2* KO females (**Supplemental Figure 7F**). We note that this analysis could not be performed in *Hnrnph2^P209L/Y^* males due to the low number of mutant males surviving to 6 weeks and the failure of these mice to pass atlas alignment quality control in all but one case. No significant differences were detected in any of the 356 cortical, white matter, subcortical, or CSF defined regions when normalized to total brain tissue volume, suggesting that brain growth was relatively preserved in mutant mice (**Supplemental Table 1**).

### Neurons cultured from hnRNPH2 P209L mice show defects in dendritic arborization of cortical neurons

To characterize the possible developmental abnormalities in the central nervous system of the mutant mice, we performed a systematic histological analysis of hemizygous male mutant or KO mice and their WT littermates. Hematoxylin and eosin (H&E) staining revealed no gross abnormalities in the brains of hnRNPH2 P209L, R206W, or KO male mice (**Supplemental Figure 8A**), apart from the presence of varying degrees of dilatation of the ventricles. Furthermore, immunohistochemistry with markers for astrocytes (GFAP) and microglia (IBA1) did not reveal evidence of inflammation in hnRNPH2 P209L, R206W or KO male brains compared to their WT littermate controls (**Supplemental Figure 8, B and C**). In addition, Luxol fast blue staining and immunohistochemistry using OLIG2, a specific and universal marker of oligodendrocytes in the brain, did not reveal any changes in central nervous system myelination (**Supplemental Figure 8, B and D**). Finally, immunohistochemistry using the neuronal marker NeuN, and immunofluorescence with cortical layer-specific markers SATB2 (layer II-IV), CTIP2 (layer V), and FOXP2 (layer VI) revealed neither significant cell loss nor altered lamination in the visual, somatosensory, or somatomotor cortex (**Supplemental Figure 9**).

To further examine the impact of pathogenic mutations on neuronal development at the cellular level, we assessed dendritic arborization and spine densities in neurons cultured from *Hnrnph2^P209L/Y^* and littermate control male mice. Compared with neurons cultured from control mice, neurons cultured from *Hnrnph2^P209L/Y^* mice had a dramatic reduction in the dendritic arbor (**Figure 5A**). We detected significantly reduced dendrite branch points, dendrite branch levels, and total dendritic lengths in neurons cultured from *Hnrnph2^P209L/Y^* mice (**Figure 5B**) that led to a large reduction in dendritic arbor complexity as assessed by Sholl analysis (**Figure 5C**). The total number of spines was also reduced in the *Hnrnph2^P209L/Y^* mice, although this difference did not reach statistical significance (**Figure 5D**). Notably, the spine density (number of spines per 10 μm) was comparable between WT and P209L mice (**Figure 5E**), suggesting that the observed reduction in total spine number can be attributed to reduced dendritic length. Taken together, these results suggest that overall brain growth is relatively preserved in hnRNPH2 mutant mice. However, dendritic arbor complexity is reduced in these mutant mice, suggesting that neural connection and activity may be affected.

**Figure 5.**
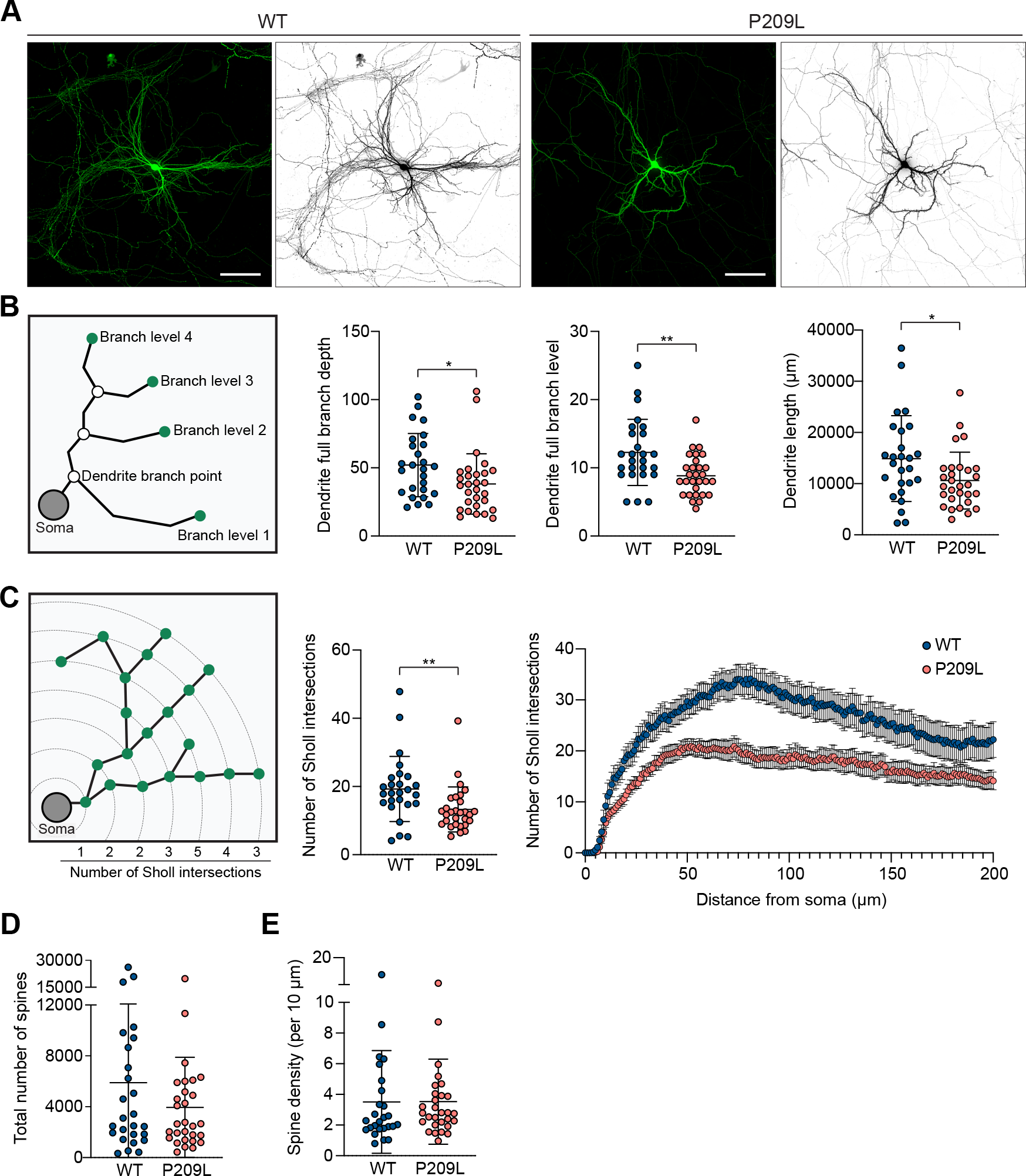
Pathogenic mutation of hnRNPH2 impairs dendritic arborization of cortical neurons. (**A**) Representative micrographs of cortical neurons isolated from WT and *Hnrnph2^P209L/Y^* mice. Cortical cells at P0 from *Hnrnph2^P209L/Y^*and littermate control mice were transfected with GFP at 7 DIV, and dendritic morphology was examined at 14 DIV. Scale bar, 100 μm. (**B**) Diagram of morphometric measurements and quantified data. Error bars represent mean ± SD. Dendrite full branch depth: \**P* = 0.0281; dendrite full branch level: \*\**P* = 0.0022; dendrite length: \**P* = 0.0281 *Hnrnph2^P209L/Y^* vs. *Hnrnph2^X/Y^* by unpaired t test. (**C**) Sholl analysis of neurons isolated from WT and *Hnrnph2^P209L/Y^* mice. The average number of Sholl intersections across the entire neuron (error bars represent mean ± SD) and the average number of Sholl intersections with 1 μm intervals (error bars represent mean ± SEM) are plotted. \*\**P* = 0.0084 *Hnrnph2^P209L/Y^* vs. *Hnrnph2^X/Y^* by unpaired t test. (**D** and **E**) Quantification of the total number of dendritic spines across the entire cortical neuron (**D**) and the average density of dendritic spines per 10 μm distance along the dendrites (**E**). Error bars represent mean ± SD. Data represent n = 26 WT and n = 29 P209L neurons.

### hnRNPH2 P209L and R206W mice, but not KO mice, have impaired motor function

We next sought to investigate the functional consequence of disease-associated mutations on behavior in our mouse model. In humans, pathogenic variants in *HNRNPH2* are associated with developmental delay, most often characterized by significant motor abnormalities accompanied by severe expressive and receptive language impairment (26); thus, we first focused on motor function. Mice selected for behavioral phenotyping were first subjected to an observational test battery at 8 weeks of age to obtain an initial and broad screen of phenotypes. Using a modified version of the SHIRPA level 1 protocol, a standardized protocol for comprehensive behavioral and functional assessment (27, 28), we found that global abnormality scores were significantly higher in *Hnrnph2^P209L/Y^* males compared to their WT littermates, whereas all other mutant and KO males and all females did not differ significantly from controls (**Figure 6A**). When abnormality scores were generated for specific functions tested in SHIRPA, we found that motor function was significantly impaired in *Hnrnph2^R206W/Y^* males, *Hnrnph2^P209L/Y^*males, and *Hnrnph2^P209L/X^* females compared to WT littermate controls (**Supplemental Figure 10A**). In addition, SHIRPA scores for autonomic function were significantly increased in hnRNPH2 P209L females and tended to be increased in males compared to WT littermates. In contrast, male KO mice had decreased autonomic function scores compared to WT controls, reflecting their better performance in assays of autonomic function (**Supplemental Figure 10B**). Scores for sensory and neuropsychiatric functions were not significantly different between *Hnrnph2* mutant or KO mice and their littermate controls (data not shown).

**Figure 6.**
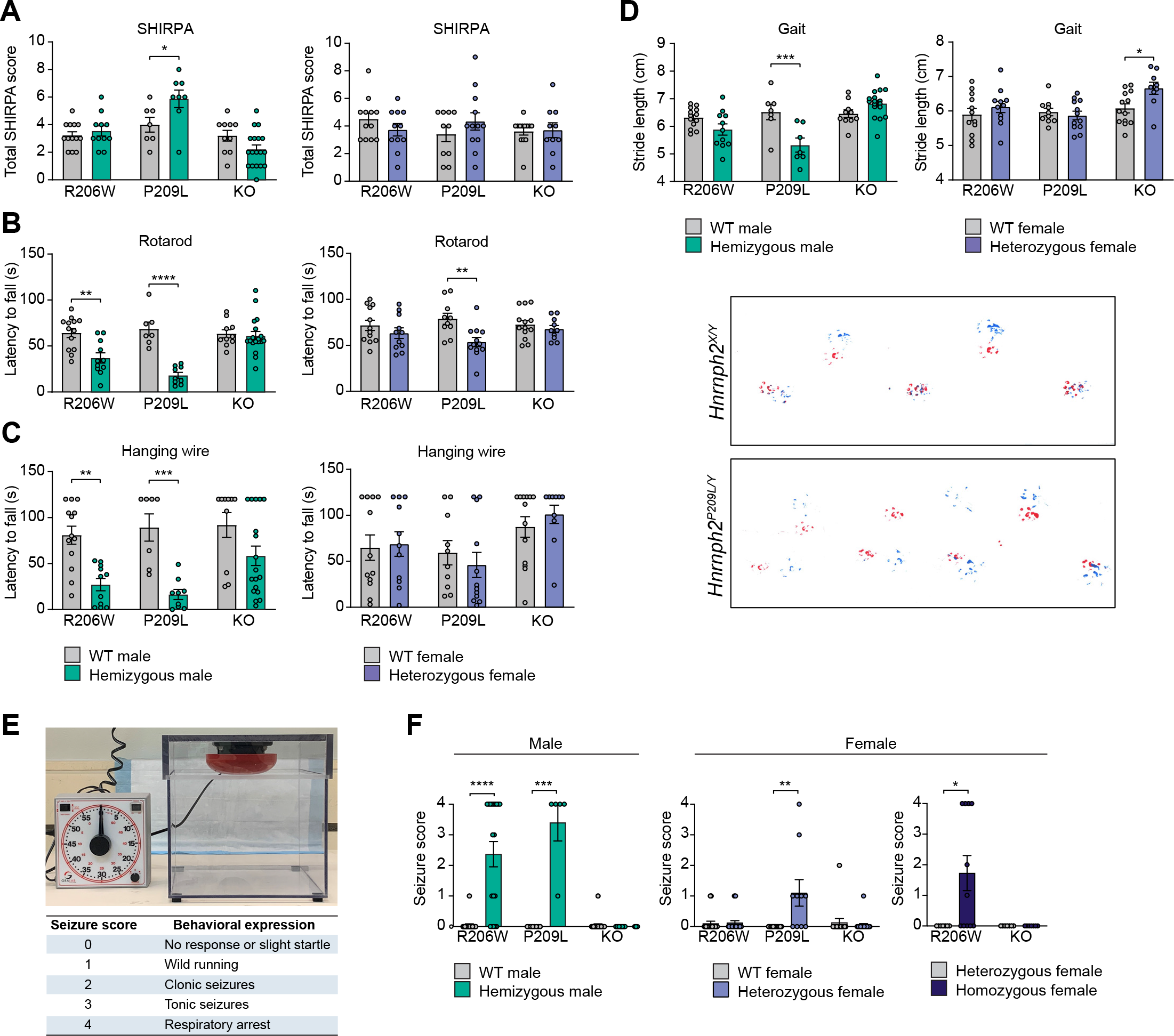
hnRNPH2 mutant mice have impaired motor function and increased susceptibility to audiogenic seizures. (**A**) Total SHIRPA abnormality scores; \**P* = 0.0397 *Hnrnph2^P209L/Y^* vs. *Hnrnph2^X/Y^* by two-way non-parametric ANOVA with Mann-Whitney U test for groupwise comparisons. (**B**) Latency to fall from rotarod; \*\**P* = 0.001 *Hnrnph2^R206W/Y^* vs. *Hnrnph2^X/Y^*; \*\*\*\**P* < 0.0001 *Hnrnph2^P209L/Y^* vs. *Hnrnph2^X/Y^*; \*\**P* = 0.0036 *Hnrnph2^P209L/X^* vs. *Hnrnph2^X/X^* by two-way ANOVA with Sidak’s multiple comparisons test. (**C**) Latency to fall from a wire cage top; \*\**P* = 0.0017 *Hnrnph2^R206W/Y^* vs. *Hnrnph2^X/Y^*; \*\*\**P* = 0.0005 *Hnrnph2^P209L/Y^* vs. *Hnrnph2^X/Y^* by two-way ANOVA with Sidak’s multiple comparisons test. (**D**) Quantification of stride length and representative images showing gait of an hnRNPH2 P209L male and WT littermate; Red and blue paints indicate the front and hind paws of each animal, respectively. \*\*\**P* = 0.0005 *Hnrnph2^P209L/Y^* vs. *Hnrnph2^X/Y^*; \**P* = 0.0265 *Hnrnph2^KO/X^* vs. *Hnrnph2^X/X^* by two-way ANOVA with Sidak’s multiple comparisons test. Group sizes for SHIRPA and motor tests were as follows: *Hnrnph2^R206W/Y^* (n = 11) vs. *Hnrnph2^X/Y^*(n = 13); *Hnrnph2^P209L/Y^* (n = 8) vs. *Hnrnph2^X/Y^* (n = 7); *Hnrnph2^KO/Y^* (n = 18) vs. *Hnrnph2^X/Y^* (n = 10); *Hnrnph2^R206W/X^* (n = 11) vs. *Hnrnph2^X/X^* (n = 12); *Hnrnph2^P209L/X^* (n = 12) vs. *Hnrnph2^X/X^* (n = 10); *Hnrnph2^KO/X^* (n = 10) vs. *Hnrnph2^X/X^* (n = 13) (**E**) Audiogenic seizure chamber and scoring of seizure behavior. (**F**) Audiogenic seizure severity score; \*\*\*\**P* < 0.0001 *Hnrnph2^R206W/Y^* vs. *Hnrnph2^X/Y^*; \*\*\**P* = 0.0009 *Hnrnph2^P209L/Y^* vs. *Hnrnph2^X/Y^*; \*\**P* = 0.0027 *Hnrnph2^P209L/X^* vs. *Hnrnph2^X/X^*; \**P* = 0.026 *Hnrnph2^R206W/X^* vs. *Hnrnph2^R206W/R206W^* by two-way non-parametric ANOVA with Mann-Whitney U test for groupwise comparison. Group sizes for audiogenic seizure susceptibility were as follows: *Hnrnph2^R206W/Y^* (n = 19) vs. *Hnrnph2^X/Y^* (n = 22); *Hnrnph2^P209L/Y^* (n = 5) vs. *Hnrnph2^X/Y^* (n = 8); *Hnrnph2^KO/Y^* (n = 20) vs. *Hnrnph2^X/Y^*(n = 22); *Hnrnph2^R206W/X^* (n = 24) vs. *Hnrnph2^X/X^* (n = 19); *Hnrnph2^P209L/X^* (n = 10) vs. *Hnrnph2^X/X^*(n = 12); *Hnrnph2^KO/X^* (n = 20) vs. *Hnrnph2^X/X^* (n = 15); *Hnrnph2^R206W/X^* (n = 7) vs. *Hnrnph2^R206W/R206W^*(n = 11); *Hnrnph2^KO/X^* (n = 7) vs. *Hnrnph2^KO/KO^* (n = 7). Error bars represent mean ± SEM in all graphs.

We next tested mice in specific and sensitive tests of motor function, including balance, coordination, and muscle strength. Rotarod performance was significantly impaired in *Hnrnph2^P209L/Y^* males, *Hnrnph2^R206W/Y^*males, and *Hnrnph2^P209L/X^* females compared with WT littermates, whereas all other mutant or KO males and females did not differ significantly from controls (**Figure 6B**). A similar impairment of balance and coordination was observed for *Hnrnph2^P209L/Y^*and *Hnrnph2^R206W/Y^* males in the beam walking test, with significantly increased time to cross (**Supplemental Figure 10C**), increased number of hind paw slips, and decreased neurological score compared to WT littermate controls (data not shown). Latency to fall from a wire cage top was significantly decreased in *Hnrnph2^P209L/Y^* and *Hnrnph2^R206W/Y^*males (**Figure 6C**) and grip strength was significantly decreased in *Hnrnph2^P209L/Y^* males (**Supplemental Figure 10D**). Finally, gait analysis revealed that *Hnrnph2^P209L/Y^* males had significantly decreased stride length compared to WT littermates (**Figure 6D**), whereas overlap, front base width, and hind base width were unchanged (data not shown).

### hnRNPH2 P209L and R206W mice, but not KO mice, have increased susceptibility to audiogenic seizures

Although the SHIRPA sensory function screen did not reveal abnormalities in *Hnrnph2* mutant or KO mice, hnRNPH2 patients have reported sensory issues including hypo- and hypersensitivity to pain, temperature, touch, and in some cases sound (2). Therefore, we next tested mice in specific and sensitive tests of sensory function, including visual acuity, olfactory function, and pain perception. We did not find any significant impairment for any of the *Hnrnph2* mutant or KO mice in the optomotor response, hot plate, or scent habituation tests (**Supplemental Figure 11**). However, we did find a significant increase in audiogenic seizure susceptibility, which has been used as a measure of both sensory hypersensitivity and epilepsy in mouse models of monogenic autism (29, 30). At postnatal day 21, *Hnrnph2^P209L/Y^* males, *Hnrnph2^R206W/Y^*males, and *Hnrnph2^P209L/X^* females had significantly increased incidence and severity of audiogenic seizures (**Figure 6, E and F**). Similarly, *Hnrnph2^R206W/R206W^*females were more susceptible to audiogenic seizures compared to *Hnrnph2^R206W/X^*female littermates (**Figure 6F**). In contrast, *Hnrnph2* KO mice showed no significant audiogenic seizure behavior (**Figure 6F**).

### hnRNPH2 P209L mice show increased delta power and epileptiform activity

Nearly half of patients with mutations in *HNRNPH2* have been reported to have a clinical seizure, and ∼10% have been reported to have abnormal brain activity by electroencephalography (EEG) without clinical seizures (2). Thus, we performed the EEG analysis in the hnRNPH2 mutant mice to measure their baseline brain and epileptic activity. To this end, we implanted electrodes near the lambdoid suture (superior/inferior colliculus regions) and between coronal and lambdoid sutures (somatosensory/visual cortex regions) to record local field potentials for 24-hour period. (**Supplemental Figure 12A**). We observed increased cortical power in *Hnrnph2^P209L/Y^* mice compared to WT littermate controls during the dark phase at the lead implanted at the right lambdoid suture (**Supplemental Figure 12B**), mainly due to increased power in the delta spectrum (**Supplemental Figure 12C**). However, this difference was absent during the light phase (**Supplemental Figure 12, D and E**) and both light and dark phases when the lead was implanted between the right coronal and lambdoid sutures (**Supplemental Figure 12, F and G**). Delta power is often associated with seizure susceptibility in mice (31). Quantification of epileptiform activity revealed that *Hnrnph2^P209L/Y^*mice had an increased percentage of time spent exhibiting spiking events during the dark phase (**Supplemental Figure 12, H and I**).

Video analysis of individual mice showed no seizure-like behavior in all four WT mice in the experiment; however, two of three *Hnrnph2^P209L/Y^*mice did show seizure-like behavior. One *Hnrnph2^P209L/Y^* mouse exhibited absence-like seizure activity for much of the 1 hour, characterized by high electrographic activity and immobility. The second *Hnrnph2^P209L/Y^* mouse had a seizure event with abnormal EEG waveforms (**Supplemental Figure 12J**) that included forelimb clonus followed by a curling of the entire body with forelimb clonus. Together, these data suggest that mutation to *HNRNPH2* increases EEG delta power and enhances seizure susceptibility.

### hnRNPH2 R206W males have increased anxiety, impaired spatial learning and memory, deficits in social interaction, and reduced marble burying

In addition to significant motor problems, pathogenic variants in *HNRNPH2* in humans are associated with intellectual disability and psychiatric diagnoses and concerns, including anxiety, autism spectrum disorder, social communication disorder, and obsessive-compulsive disorder or stereotyped behaviors (2).

Therefore, we evaluated 8-week-old *Hnrnph2^R206W/Y^* males and their WT littermates for anxiety, learning and memory, social interaction, and repetitive behavior. We note that we could not perform the same assays with *Hnrnph2^P209L/Y^* mice due to their high mortality (**Figure 3D**).

We began with the open field test, which is widely used to evaluate anxiety-like behavior by measuring the amount of time spent in the open center area. In this test, mutant males spent significantly less time in the center zone compared to controls, despite similar locomotor activity as measured by total distance traveled across the 20-minute test (**Figure 7A**). When data was analyzed in 5-minute time bins, the distance travelled within the first 5 minutes of the test was significantly lower for *Hnrnph2^R206W/Y^* males compared to controls, but was similar to controls for the remainder of the test (**Supplemental Figure 13A**). In contrast, the percentage of time spent in the center was significantly reduced in mutants across all 4 time bins (**Supplemental Figure 13A**). Together, these results suggest increased anxiety with impaired habituation to a novel environment in the *Hnrnph2^R206W/Y^*males.

**Figure 7.**
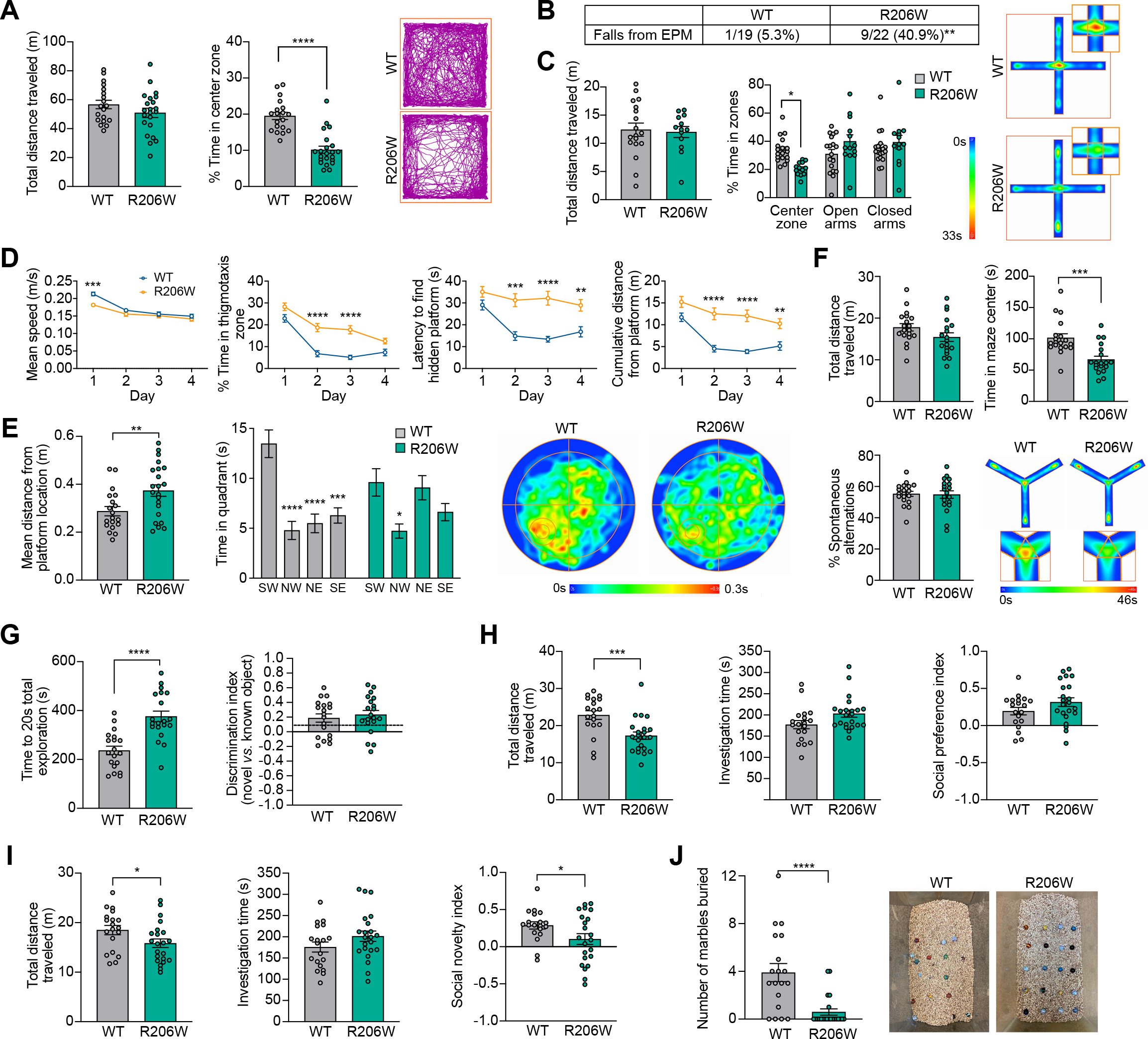
hnRNPH2 R206W males have increased anxiety, impaired spatial learning and memory, social interaction deficits and reduced marble burying. (A) Total distance traveled (left) and percentage time spent in the center zone (\*\*\*\**P* < 0.0001 *Hnrnph2^R206W/Y^* (n = 22) vs. *Hnrnph2^X/Y^*(n = 19) by unpaired t test) of the open field across the duration of the test. Open field track plot (right) of a representative *Hnrnph2^R206W/Y^* male and WT littermate showing the position of the animals’ center points for the duration of the test. (**B**) Incidence of falls from the elevated plus maze (\*\**P* = 0.0109 *Hnrnph2^R206W/Y^* vs. *Hnrnph2^X/Y^* by Fisher’s exact test). Review of videos recorded of the tests showed that WT mice and most of the mutant mice fell while turning around at the distal end of an open arm. The remaining mutant mice fell when climbing from the center of the maze onto the small ledge on the outside of a closed arm, and one mutant jumped from the end of an open arm. All mice that fell from the maze were excluded from further analysis. (**C**) Total distance traveled (left) and percentage time spent in the center zone, open arms, and closed arms (middle; \**P* = 0.0149 *Hnrnph2^R206W/Y^* (n = 13) vs. *Hnrnph2^X/Y^*(n = 18) by two-way ANOVA with Sidak’s multiple comparisons test) of the elevated plus maze. At right is an averaged heat map of the animals’ center point for the groups for the entire duration of the 5-minute test. Insets shows enlarged center zone of the maze. (**D**) Mean speed (day 1 \*\*\**P* = 0.0002 *Hnrnph2^R206W/Y^* (n = 22) vs. *Hnrnph2^X/Y^* (n = 19)), percentage time spent in the thigmotaxis zone (day 2 \*\*\*\**P* < 0.0001, day 3 \*\*\*\**P* < 0.0001 *Hnrnph2^R206W/Y^* (n = 22) vs. *Hnrnph2^X/Y^* (n = 19)), latency to find the hidden platform (day 2 \*\*\**P* = 0.0002, day 3 \*\*\*\**P* < 0.0001, day 4 \*\**P* = 0.0058 *Hnrnph2^R206W/Y^* (n = 22) vs. *Hnrnph2^X/Y^*(n = 19)), and cumulative distance from the hidden platform (day 2 \*\*\*\**P* < 0.0001, day 3 \*\*\*\**P* < 0.0001, day 4 \*\**P* = 0.0063 *Hnrnph2^R206W/Y^* (n = 22) vs. *Hnrnph2^X/Y^* (n = 19)) during the training phase of the Morris water maze by two-way ANOVA with Sidak’s multiple comparisons test. (**E**) Mean distance from platform location (\*\**P* = 0.0092 *Hnrnph2^R206W/Y^* (n = 22) vs. *Hnrnph2^X/Y^* (n = 19) by unpaired t test) and time spent in each quadrant (\*\*\*\**P* < 0.0001 SW vs. NW, \*\*\*\**P* < 0.0001 SW vs. NE, \*\*\**P* = 0.0002 SW vs. SE for *Hnrnph2^X/Y^* (n = 19); \**P* = 0.0198 SW vs. NW for *Hnrnph2^R206W/Y^* (n = 22) by two-way ANOVA with Tukey’s multiple comparisons test) during the probe trial of the Morris water maze. At right is an averaged heat map of the animals’ center point for the groups for the entire duration of the probe trial. (**F**) Total distance traveled, percent spontaneous alternations, and time spent in center zone of the Y maze (\*\*\**P* = 0.0002 *Hnrnph2^R206W/Y^* (n = 19) vs. *Hnrnph2^X/Y^*(n = 19) by unpaired t test). At lower right is an averaged heat map of the animals’ center point for the groups for the entire duration of the 8-minute test. Insets shows enlarged center zone of the maze. (**G**) Time to reach 20-second total object exploration time (\*\*\*\**P* < 0.0001 *Hnrnph2^R206W/Y^* (n = 20) vs. *Hnrnph2^X/Y^* (n = 19) by unpaired t test) and novel object discrimination index in the testing stage of the novel object recognition test. Dotted line indicates the previously published novel object discrimination index for C57BL/6J mice with a 24 hour test interval. (**H**) Total distance traveled (\*\*\**P* = 0.0009 *Hnrnph2^R206W/Y^* (n = 22) vs. *Hnrnph2^X/Y^* (n = 19) by unpaired t test), investigation time, and social preference index in the three-chamber social preference test. (**I**) Total distance traveled (\**P* = 0.0442 *Hnrnph2^R206W/Y^* (n = 22) vs. *Hnrnph2^X/Y^* (n = 19) by unpaired t test), investigation time, and social novelty index (\**P* = 0.0489 *Hnrnph2^R206W/Y^* (n = 22) vs. *Hnrnph2^X/Y^* (n = 19) by unpaired t test) in the three- chamber social novelty test. (**J**) Number of marbles buried in the marble burying test. \*\*\*\**P* < 0.0001 *Hnrnph2^R206W/Y^* (n = 22) vs. *Hnrnph2^X/Y^* (n = 19) by unpaired t test. Representative cages were photographed from above after the test. Error bars represent mean ± SEM in all graphs.

To further explore the anxiety phenotype in these mice, we used the elevated plus maze.

In this test, we noticed that significantly more R206W mutants fell from the maze as compared to controls (**Figure 7B**). The frequent falls of the mutant mice off the maze were unexpected and may be related to increased anxiety levels and/or impaired motor function. For mice that completed the 5-minute test, we found no significant difference in the total distance traveled, suggesting similar locomotor activity (**Figure 7C**). However, the percentage of time spent in the center zone was significantly reduced in *Hnrnph2^R206W/Y^*males, whereas the percentage of time spent in the open or closed arms was comparable between control and mutant mice (**Figure 7C**). As time spent in the center of the elevated plus maze has been linked with decision making related to approach/avoid conflict (32), this decrease suggests impaired decision making and risk assessment in the *Hnrnph2^R206W/Y^* males. When analyzing data in 1-minute time bins, we found that WT males, but not *Hnrnph2^R206W/Y^* males, spent significantly less time in the open arms and more time in the closed arms over time (**Supplemental Figure 13B**). Increasing open-arm platform avoidance and closed-arm preference is expected as the test progresses, a change in behavior that is considered to reflect the avoidance of potentially dangerous sections of the maze (33). Thus, the failure of mutant mice to avoid the more dangerous open arm platforms over time could be due to impaired spatial learning or impaired decision making and risk assessment (33, 34).

To assess spatial learning and memory, we used the Morris water maze, which assesses cued learning, spatial learning, and memory as determined by preference for the platform area when the platform is absent. During the cued phase of the test, there was no significant difference in mean speed or percentage thigmotaxis (percentage time spent within 10 cm of the pool wall) between *Hnrnph2^R206W/Y^* males and WT littermates (**Supplemental Figure 13C**). Although mutants took longer to find and climb onto the visible platform in early trials, both groups performed similarly in later trials (**Supplemental Figure 13C**). Together, this data suggests that *Hnrnph2^R206W/Y^* males possess the basic abilities, strategies, and motivation necessary to complete the spatial version of the task (35). During the spatial learning phase of the test, *Hnrnph2^R206W/Y^* males had significantly reduced swim speed on day 1, but not days 2 to 4 (**Figure 7D**). Percent thigmotaxis was increased in mutants compared to controls on days 2 and 3 of training, in contrast to their behavior during the cued trials (**Figure 7D**). Given that thigmotaxis has been linked to anxiety, stress, and fear (36, 37), it is possible that repeated swimming trials over multiple training days increased anxiety in the *Hnrnph2^R206W/Y^* males.

Thigmotaxis can also interfere with an animal’s ability to learn the location of the hidden platform, and it was therefore not surprising that the latency to find the platform, as well as the cumulative distance from the platform, were both significantly reduced in the mutant group on days 2 to 4 of training (38) (**Figure 7D** and **Supplemental Figure 13D**). In the probe trial to assess memory, *Hnrnph2^R206W/Y^* males did not demonstrate significant thigmotaxis, but did have significantly slower mean swim speed compared to WT littermates (**Supplemental Figure 13E**). Furthermore, in contrast to controls, mutant mice did not show a clear preference for the target quadrant, and mean distance from the target was significantly higher (**Figure 7E**). Given the increased thigmotaxis and apparent reduced spatial learning in the learning phases of the test, it is not possible to say whether the impaired performance in the probe trial was due to deficits in spatial learning, spatial memory, or both.

Next, we tested the mice in the Y maze spontaneous alternation test, a measure of working spatial memory. In this maze, mice typically prefer to investigate a new arm of the maze rather than returning to one that was previously visited. Total distance traveled during the test was not significantly different between groups, suggesting comparable locomotor activity (**Figure 7F**). However, the number of arm entries was significantly reduced in *Hnrnph2^R206W/Y^* males compared to WT littermates (**Supplemental Figure 13F**). As was observed in the elevated plus maze, the *Hnrnph2^R206W/Y^* males spent significantly less time in the center zone (**Figure 7F**), the region of the Y maze in which working memory is thought to be engaged (39). The percentage of spontaneous alternations was not significantly different between mutants and controls (**Figure 7F**). As there was no significant correlation between total number of arm entries and percent spontaneous alternations (**Supplemental Figure 13F**), these results suggest that *Hnrnph2^R206W/Y^* males have intact spatial working memory in the Y maze.

We next performed the novel object recognition test, which assesses visual recognition memory. In this test, mice are presented with a choice between novel and familiar objects, with normal behavior typically reflecting rodents’ innate preference for novelty. In the familiarization phase of the test, *Hnrnph2^R206W/Y^*males took longer to reach 20 seconds of total exploration of both objects, suggesting decreased exploration. For 20-second total exploration, neither mutants nor WT littermates showed significant preference for identical objects placed at the top left versus bottom right of the open field (**Supplemental Figure 13G**). In the testing phase of the test performed 24 hours later, *Hnrnph2^R206W/Y^* males again had significantly increased total time to reach 20 seconds of object exploration. For 20-second total exploration, *Hnrnph2^R206W/Y^* males showed no significant deficit in novel object recognition as measured by the discrimination index (**Figure 7G**).

We next used the three-chamber social test to assess social interaction and preference.

In the social preference test, *Hnrnph2^R206W/Y^* males showed a significant reduction in total distance traveled, indicating reduced locomotor activity (**Figure 7H**). However, total investigation time (social plus non-social stimulus investigation time) and social preference index was not significantly different between groups (**Figure 7H**). Similarly, in the social novelty test, mutant males had a significant decrease in total distance traveled, but similar total investigation time (novel plus known social investigation time) (**Figure 7I**). However, the social novelty index was significantly reduced in *Hnrnph2^R206W/Y^* males compared to littermate controls (**Figure 7I**). Together, these data suggest that while *Hnrnph2^R206W/Y^* males have intact social preference, they exhibit deficits in social recognition memory.

Lastly, we performed the marble burying test to assess any potential repetitive, obsessive compulsive-like behavior in *Hnrnph2^R206W/Y^* males. In this test, the number of marbles buried correlates with the intensity of the mice’s repetitive or compulsive digging behavior. To our surprise, we found that mutant males buried significantly fewer marbles compared to WT littermates (**Figure 7J**), suggesting a deficit in species-typical digging behavior (40).

### Pathogenic variants alter the nucleocytoplasmic ratio of hnRNPH2 in mice

To examine the impact of disease-causing mutations on the subcellular localization of hnRNPH2 in mice, we performed immunoblot analysis on nuclear and cytoplasmic fractions of cortical tissue. Nuclear hnRNPH2 levels were significantly reduced in *Hnrnph2^P209L/Y^* males and tended to be decreased in *Hnrnph2^R206W/Y^*males (**Supplemental Figure 14, A and B**). As we were unable to detect any hnRNPH2 protein in the cytoplasmic fraction of mutants or WT littermate controls by immunoblot (data not shown), we next examined brain sections including hippocampus and cerebellum using immunofluorescence, a technique that is more sensitive than immunoblot. Using an antibody specific for hnRNPH2, we observed cytoplasmic staining in neurons of both R206W and P209L mutants, but not in WT littermate controls (**Supplemental Figure 14, C–E**).

Together, these results suggest that disease-causing mutations modestly alter the subcellular localization of hnRNPH2 in neurons of mice, similar to what we observed for human hnRNPH2 in HeLa cells. Importantly, despite mutations to the PY-NLS, the vast majority of mutant hnRNPH2 was correctly localized in the nucleus of mouse neurons in vivo (**Supplemental Figure 14, C–E**), consistent with our observations in HeLa cells (**Figure 1, C–E**).

### Expression of *Hnrnph1* is increased in *Hnrnph2* KO mice but not in hnRNPH2 P209L or R206W mice

*Hnrnph1* is the autosomal conserved paralog of *Hnrnph2* and the two genes are believed to play similar and potentially redundant roles in RNA splicing (41). Mutations in *HNRNPH1* are also associated with a neurodevelopmental syndrome identified in boys that is very similar to hnRNPH2-linked phenotypes (9, 10). Given the high degree of homology between the two genes and the possibility of redundancy in function, we examined the expression of *Hnrnph1* in our *Hnrnph2* mutant and KO mice using digital droplet RT-PCR (ddRT-PCR). In the adult cortex, *Hnrnph1* mRNA levels were significantly increased in male *Hnrnph2* KOs, but not in P209L or R206W mutant males (**Figure 8A**). The increase of *Hnrnph1* mRNA in male *Hnrnph2* KO mice was accompanied by an increase in hnRNPH1 protein levels (**Supplemental Figure 15A**). In contrast, expression levels of two other members of the hnRNP F/H family, *Hnrnpf* and *Hnrnph3*, remained unaltered in both *Hnrnph2* mutant and KO mice (**Supplemental Figure 15B**). *Hnrnph2* mRNA levels were significantly decreased in male *Hnrnph2* KO mice, as expected for a transcript subject to nonsense-mediated decay, but unchanged in hemizygous P209L or R206W mutant males. Similar trends were observed for both transcripts in female mice, but differences were not statistically significant (**Figure 8A**).

**Figure 8.**
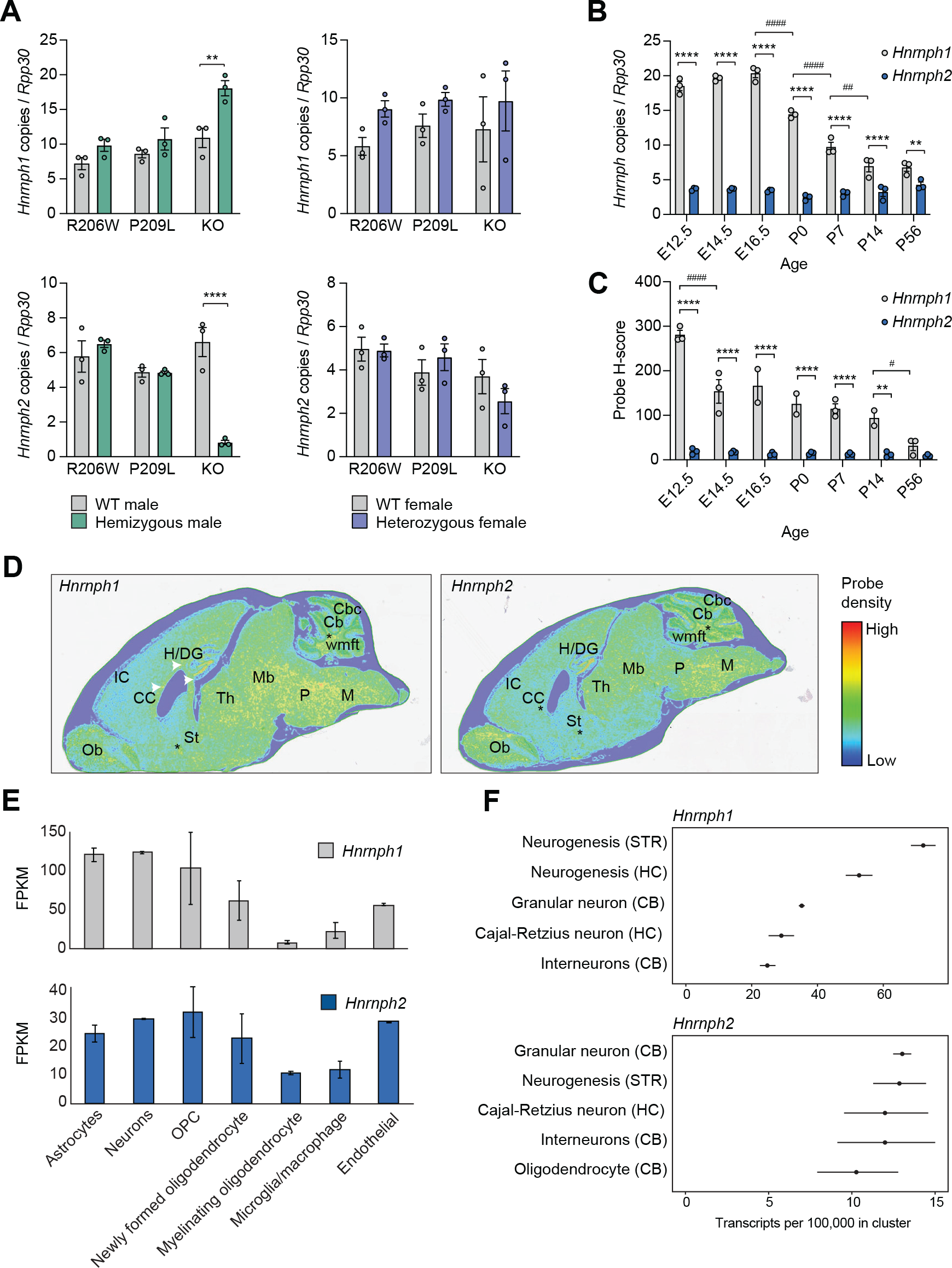
Spatiotemporal expression of *Hnrnph1* and *Hnrnph2* in mouse brain. (**A**) Number of *Hnrnph1* and *Hnrnph2* copies normalized to *Rpp30* in the cortex of *Hnrnph2^R206W^*, *Hnrnph2^P209L^*, and *Hnrnph2^KO^*mice by ddRT-PCR. Error bars represent mean ± SEM. *Hnrnph1* copies: \*\**P* = 0.0021 *Hnrnph2^KO/Y^* vs. *Hnrnph2^X/Y^*; *Hnrnph2* copies: \*\*\*\**P* < 0.0001 *Hnrnph2^KO/Y^*vs. *Hnrnph2^X/Y^* by two-way ANOVA with Sidak’s multiple comparisons test. n = 3 for all groups. (**B**) Number of *Hnrnph1* and *Hnrnph2* copies normalized to *Rpp30* in the cortex of WT C67BL/6J mice across 7 prenatal and postnatal developmental timepoints by ddRT-PCR; error bars represent mean ± SEM. \*\*\*\**P* < 0.0001 and \*\**P* = 0.006 *Hnrnph1* vs. *Hnrnph2* by two-way ANOVA with Sidak’s multiple comparisons test, ^####^*P* < 0.0001 *Hnrnph1* E16.5 vs. P0, P0 vs. P7, ^##^*P* = 0.0049 *Hnrnph1* P7 vs. P14 by two-way ANOVA with Tukey’s multiple comparisons test. n = 3 for all groups. (**C**) H-scores for *Hnrnph1* and *Hnrnph2* probes in the brain of WT C67BL/6J mice across 7 prenatal and postnatal developmental timepoints by Halo analysis of BaseScope ISH; error bars represent mean ± SEM. \*\*\*\**P* < 0.0001 *Hnrnph1* vs. *Hnrnph2* at E12.5 – P7 and \*\**P* = 0.0018 *Hnrnph1* vs. *Hnrnph2* at P14 by two-way ANOVA with Sidak’s multiple comparisons test, ^####^*P* < 0.0001 *Hnrnph1* E12.5 vs. E14.5, ^#^*P* = 0.046 *Hnrnph1* P14 vs. P56 by two-way ANOVA with Tukey’s multiple comparisons test. n = 3 for all groups. (**D**) Regional expression of *Hnrnph1* and *Hnrnph2* across the adult (P56) mouse brain by BaseScope ISH. White arrowheads show the location of the corpus callosum, asterisks indicate the location of fiber tracts. Ob, olfactory bulb; CC, corpus callosum; IC, cerebral cortex/isocortex; H/DG, hippocampus/dentate gyrus; St, bed of nuclei of the stria terminalis; P, pons; M, medulla; Th, thalamus; Mb, rostral collicular midbrain; Cb, cerebellum; Cbc, cerebellar cortex; wmft, white matter fiber tracts. (**E**) Expression of *Hnrnph1* and *Hnrnph2* in the major classes of brain cells at P7 by RNA. Data extracted from Brain RNA-seq (46, 47). FPKM, fragments per kilobase of transcript per million mapped reads. Error bars represent mean ± SD. (**F**) Top 5 expression cell clusters for *Hnrnph1* and *Hnrnph2* in the adult mouse brain by single-cell RNA sequencing (48).

To test whether the increase of *Hnrnph1* mRNA in our KO mice was a consequence of loss of *Hnrnph2*, we depleted *HNRNPH2* using siRNA and measured *HNRNPH1* RNA levels in HEK293T cells. At ∼30% KD of *HNRNPH2* (12 nM RNAi), we did not detect any increase in *HNRNPH1* transcript levels. However, when we reached ∼60% KD of *HNRNPH2* (18 nM RNAi), we detected an increase of ∼15% in *HNRNPH1* transcript levels. At ∼88% and ∼98% KD of *HNRNPH2* (24 and 30 nM RNAi), we found ∼25% and ∼55% increases of *HNRNPH1* transcript levels, respectively (**Supplemental Figure 15C**), suggesting that loss of hnRNPH2 leads to transcriptional upregulation of *HNRNPH1*.

To explore the possibility that the increase in *Hnrnph1* expression may compensate for the loss of *Hnrnph2* in KO mice, we investigated the spatiotemporal expression of these two genes. Assessment of the Allen mouse brain atlas revealed that *Hnrnph1* is expressed at high levels across all 12 major regions of the adult mouse brain, whereas *Hnrnph2* expression is detected at low levels in the olfactory areas and cortical subplate only (42). To examine the spatiotemporal expression of these two genes during mouse brain development, we performed ddRT-PCR and ISH on WT C57BL/6J mice at embryonic day 12.5, 14.5, 16.5, and postnatal day 0, 7, 14, and 56. Quantification of the ISH generated an “H-score” reflecting the level of mRNA expression in the tissue section based on the detection of specific probe signal in cells of interest. In the cortex, *Hnrnph1* was expressed at significantly higher levels than *Hnrnph2* at all time points examined by ddRT-PCR (**Figure 8B**). Furthermore, whereas *Hnrnph1* mRNA levels decreased significantly after E16.5, *Hnrnph2* mRNA levels did not significantly change over the course of the 7 developmental time points (**Figure 8B**). ISH on whole brains showed similar results, with the H-score for *Hnrnph1* being significantly higher than that for *Hnrnph2* at all time points except P56. Furthermore, *Hnrnph1* H-scores significantly decreased after E12.5, whereas *Hnrnph2* H-scores remained stable over all time points (**Figure 8C**). Spatial expression analysis of adult (P56) brains revealed that both *Hnrnph1* and *Hnrnph2* were expressed in similar areas, including regions within the telencephalon, brain stem, and hindbrain, as well as fiber tracts (**Figure 8D**). A similar pattern of spatiotemporal expression has been reported for *HNRNPH1* and *HNRNPH2* in human brain (43, 44) and human brain organoids (45) (**Supplemental Figure 15, D–F**). In sum, *HNRNPH1* and *HNRNPH2* show similar spatial and temporal expression patterns in human and mouse brains. *HNRNPH1* is highly expressed during early developmental stages and decreases over time, whereas *HNRNPH2* expression is consistently modest throughout development, suggesting that hnRNPH1 may govern early brain developmental processes that are gradually shared with its homolog hnRNPH2 at later and/or post-developmental stages.

To define the specific type of mouse brain cells that express *Hnrnph1* and *Hnrnph2*, we turned to publicly available databases. At P7, RNA-seq data indicated that both genes are expressed in all of the major cell classes of the brain, including astrocytes, neurons, oligodendrocyte precursor cells, newly formed oligodendrocytes, myelinating oligodendrocytes, microglia, and endothelial cells (46, 47) (**Figure 8E**). In the adult mouse brain, single-cell RNA-seq (48) has demonstrated that the top 5 expression cell clusters for *Hnrnph1* and *Hnrnph2* show significant overlap, including cells that undergo adult neurogenesis in the striatum, granular neurons in the cerebellum, and Cajal-Retzius neurons in the hippocampus (**Figure 8F**). Together, these data suggest that *Hnrnph1* and *Hnrnph2* have closely matching expression patterns with regard to brain regions and cell types. Importantly, as development proceeds, expression of *Hnrnph1* decreases while expression of *Hnrnph2* persists; thus, normal cellular function becomes progressively more dependent on hnRNPH2. These observations support our hypothesis that upregulation of *Hnrnph1* in the setting of *Hnrnph2* KO mice compensates for the functional loss of hnRNPH2.

### Expression of pathogenic hnRNPH2 variants leads to more severe alterations in gene expression and RNA splicing compared to *HNRNPH2* KO

Finally, we evaluated the effects of pathogenic mutations in hnRNPH2 on the expression and splicing of its target RNAs. To test this, we performed RNA sequencing in 3-week-old neurons derived from human iPSCs genetically engineered to bear hnRNPH2 R206W, R206Q, or P209L mutations or KO of *HNRNPH2*. Analysis of gene expression pattern changes revealed that pathogenic variants and KO of hnRNPH2 significantly altered global gene expression (**Supplemental Figure 16, A and B**). A total of 3,745 RNAs were significantly up- or downregulated in one or more conditions as compared to WT controls (**Supplemental Figure 16A and Supplemental Table 2**).

Interestingly, the patterns of up- or downregulation caused by pathogenic mutations and KO of hnRNPH2 were very similar: genes that were up- or downregulated in KO neurons were similarly up- or downregulated in neurons expressing mutant protein (**Supplemental Figure 16B**). However, the levels of alteration were more severe in mutant-expressing neurons compared with KO neurons (**Supplemental Figure 16B**). This observation is consistent with our hypothesis that the pathomechanism of hnRNPH2-linked disorder is loss of function with an incomplete compensation by hnRNPH1. Indeed, we confirmed that *HNRNPH1* transcript levels were increased in KO neurons (152%) but were less affected in mutant-expressing neurons (99%, 108%, and 128% in R206W, P209L, and R206Q, respectively). We further noted that the severity of gene expression changes in mutant-expressing neurons was inversely correlated with the levels of *HNRNPH1* upregulation, with R206W (99% of WT expression levels) being the most severe and R206Q (128% of WT expression levels) being the least severe (**Supplemental Figure 16B**). The gene ontology (GO) terms linked to commonly up- or downregulated genes in mutant-expressing and KO neurons demonstrated enrichment for terms related to neuronal synapses, neuronal transport, ion channel, and innate immunity (**Supplemental Figure 16, C– F**). Moreover, we identified 10,691 aberrant alternative splicing events (ASEs), the most common of which were skipped exons (SE; 6,829 events), followed by mutually exclusive exons (MXE; 1,429 events), retained introns (RI; 930 events), alternative 3′ splice sites (A3SS; 843 events), and alternative 5′ splice sites (A5SS; 660 events) (**Supplemental Figure 16, G and H**).

We also performed RNA-seq analyses using the cortices of our knock-in and KO mice (**Supplemental Figure 17**). A much smaller number of differentially expressed genes was identified in mouse cortices compared to that observed in human iPSC-derived neurons, possibly due to the stochastic nature of gene expression coming from bulk tissues (**Supplemental Figure 17, A and B and Supplemental Table 3**). Due to the small number of genes altered, we were unable to perform GO analyses in R206W and KO mice; however, GO analysis of differentially expressed genes in P209L mice revealed enrichment for GO terms related to neurogenesis and differentiation, similar to those identified from the human RNA-seq analyses, and also GO terms related to muscle (**Supplemental Figure 16C**). We identified 10 genes that were significantly differentially upregulated in R206W and P209L mice, but not in *Hnrnph2* KO mice (**Supplemental Figure 17, D and E**). Interestingly, these genes all appeared as significantly differentially expressed genes in human RNA-seq analyses (**Supplemental Figure 16 and Supplemental Table 2**) and are all known to have functional roles in neurons (**Supplemental Figure 17, E and F**). For 4 of these 10 genes, pathogenic variants have been established as the cause of neurodevelopmental disorders, including *CTNNA2* (OMIM: 618174), *TNPO2* (OMIM: 619556), *ASH1L* (OMIM: 617796), and *SHANK1* (OMIM: 209850) (**Supplemental Figure 17E**). Similar to RNA-seq results from the human iPSC-derived neurons, pathogenic mutations and KO of *Hnrnph2* caused widespread changes in alternative pre-mRNA splicing (**Supplemental Figure 17, G and H**). Taken together, these results demonstrate that pathogenic mutations induce more severe transcriptome changes than KO both in human iPSC-derived neurons and in mice. Consistent upregulation of *HNRNPH1*/*Hnrnph1* in KO, but not in mutants, suggests that hnRNPH1 might be compensating for the loss of hnRNPH2 function and thereby blunting hnRNPH2-related phenotypes in the setting of *HNRNPH2* KO.

## Discussion

The hnRNP family of proteins has a significant enrichment of de novo variants associated with neurodevelopmental disorders with similar clinical phenotypes and potentially shared molecular pathogenesis (49). Mutations in *HNRNPH2* and its close paralog *HNRNPH1* provide one such example, wherein similar mutations (i.e., missense mutations frequently located in the PY-NLS) cause syndromes with overlapping symptoms (9, 10). Here we investigated the pathological mechanism underlying hnRNPH2-related disorder using in vitro studies, cell lines, and multiple knock-in and KO mouse models. Our results strongly indicate that the mechanism underlying hnRNPH2-related disease is not a simple loss of function but is instead a complex mechanism involving toxic gain of function and/or loss of hnRNPH2 function with inadequate genetic compensation by *HNRNPH1*.

Our in vitro characterization of the consequences of common pathogenic hnRNPH2 mutations revealed, as predicted, that mutations in the PY-NLS lead to a partial redistribution of hnRNPH2 protein from the nucleus to the cytoplasm. Notably, this redistribution was modest, with the majority of hnRNPH2 protein remaining in the nucleus. Furthermore, for several pathogenic variants that lie outside the PY-NLS of hnRNPH2, we observed no redistribution of the protein to the cytoplasm.

These findings are complemented by our characterization of KO and knock-in mouse models. Importantly, we found that two independent *Hnrnph2* KO mouse lines were phenotypically normal across a wide variety of measures, with a consistent absence of pathological phenotypes, consistent with ongoing phenotyping of the KOMP KO line reported by the International Mouse Phenotype Consortium. These observations strongly argue that hnRNPH2-related disease cannot be attributed to a simple loss of function. In contrast, *Hnrnph2^P209L^* and *Hnrnph2^R206W^* knock-in mice recapitulated the modest redistribution of hnRNPH2 from the nucleus to the cytoplasm while driving a highly penetrant phenotype that reproduced multiple clinical features of human disease, including facial abnormalities, seizure propensity, reduced viability in males, and several behavioral abnormalities, including reductions in motor ability. Furthermore, *Hnrnph2*^R206W^ mutant mice showed increased anxiety, impaired spatial learning and memory, and deficits in social interaction, echoing several phenotypes observed in human disease and presenting additional quantifiable measures that may be modifiable with therapeutic intervention. Indeed, the extensive, robust phenotypes observed in *Hnrnph2^P209L^* and *Hnrnph2^R206W^*mice suggest strong potential for their use in preclinical studies and to reveal novel targets for therapy.

This pattern of phenotypes in physiological models – no apparent phenotypic consequence in KO mice, robust recapitulation of disease features in knock-in mice – suggests two possible mechanisms as drivers of disease. The first possibility is a toxic gain of function, a mechanism with precedents in several common neurological diseases (e.g., ALS caused by mutations in *SOD1*, Parkinson’s disease caused by mutations in *SNCA*) in which disease phenotypes are absent in KO animals but are faithfully recapitulated in animals expressing disease mutations. However, our results are also consistent with an alternative disease mechanism in which mutations in *HNRNPH2* directly cause a loss of hnRNPH2 function, but the persistence of significant hnRNPH2 protein in the nucleus results in a failure to induce compensatory *HNRNPH1* expression. Indeed, *HNRNPH1* has an expression pattern that is nearly identical to that of *HNRNPH2* with respect to brain region and cell type, and it encodes a highly similar protein to hnRNPH2. Interestingly, whereas the expression of *HNRNPH1* decreases as development proceeds, the expression of *HNRNPH2* persists, such that cells become progressively more dependent upon hnRNPH2. In this context, our findings from RNA-seq analyses – namely, that KO of *HNRNPH2/Hnrnph2* and knock-in of pathogenic mutations alter gene expression into the same direction, where knock-in mutations induce more severe transcriptome changes than KO, and that KO of *HNRNPH2/Hnrnph2*, but not knock-in, consistently leads to upregulation of *HNRNPH1/Hnrnph1* – suggest that upregulation of *Hnrnph1* is responsible for rescuing KO animals from the consequences of the loss of hnRNPH2 function. In contrast, the introduction of disease mutations in *Hnrnph2* in mice is not accompanied by significant upregulation of *Hnrnph1*. As such, disease-causing mutations in *HNRNPH2* thwart the physiological mechanism that would otherwise compensate for the loss of hnRNPH2 protein function.

Importantly, both of these possible gain-of-function and loss-of-function mechanisms would be predicted to respond positively to therapies designed to deplete expression of mutant proteins (e.g., antisense oligonucleotides). Indeed, genetic compensation between *HNRNPH1* and *HNRNPH2* suggests a therapeutic strategy wherein knockdown of *HNRNPH2* in patients would be predicted to be well tolerated – as KO of *HNRNPH2* is well tolerated in cells and in mice – while also triggering compensatory upregulation of *HNRNPH1*. Further investigation will be needed to determine the mechanisms underlying cross-regulation of *HNRNPH1* and *HNRNPH2*, and how normal functions of hnRNPH2 are disrupted in disease. Of particularly high priority is determining the prospects for therapy aimed at knockdown of mutant *Hnrnph2* to look for upregulation of *Hnrnph1* and potential rescue of the phenotype in mice.

## Methods

### Cell culture and transfection

HEK293T (CRL-3216) and HeLa (CCL-2) cells were originally purchased from ATCC and periodically authenticated by short tandem repeat (STR) profiling. Cells were grown in Dulbecco’s modified Eagle’s medium (DMEM) supplemented with 10% fetal bovine serum (FBS), 1% penicillin/streptomycin, and 1% L-glutamate. Cells were counted using ADAM-CellT (NanoEntek), plated and transfected using Lipofectamine 3000 (Thermo Fisher; L3000008) for transient overexpression or RNAiMAX (Thermo Fisher; 13778075) for siRNA knockdown according to the manufacturer’s instructions.

### Differentiation of IPSC-derived neurons

iPSCs were differentiated into cortical neurons with a two-step protocol (pre-differentiation and maturation) as previously described (Wang et al., 2017). When IPSCs reached to %70-80 confluence, cells washed with DPBS twice and dissociated with accutase (Stemcell technologies) and collected cells were filtered with cell strainer (Stemcell technologies). Filtered Cells were Centrifuged with 200 rcf for 5 min in RT and the pellet was resuspended with medium containing N2 suplement, non-essential amino acids (NEAA), GlutaMAX™ Supplement and Y-27632 (Stem cell technologies, USA) and 1 µg/ml of Doxycycline hyclate (Sigma Aldrich, St Louis, MO, USA). For cell counting 10 ul was used (Thermo Fisher Scientific Countess).

Cells were subplated 1.2x 10^6^ cells/well in six-well dish coated with Matrigel in knockout Dulbecco’s modified Eagle’s medium (KO-DMEM)/F12 and wells were filled with same medium to 2 ml/each 6 well plate. The medium was changed daily for 3 days, and Y-27632 was removed from day 2. For maturation, pre-differentiated precursor cells were washed, dissociated, counted, and subplated at 25x10^4^ cells/ml on dishes coated with 50 ug/ml Poly-L-ornithine in BrainPhys neuronal medium (Stem cell technologies, USA) containing N2 (Thermo Fisher Scientific, Waltham, MA, USA), B-27™, 20 ng/ml BDNF (Peprotech), 20 ng/ml GDNF (Peptrotech), 500 µg/ml Dibutyryl cyclic-AMP (Sigma Aldrich, St Louis, MO, USA), 200 nM L-ascorbic acid (Sigma Aldrich, St Louis, MO, USA), 1 µg/ml Natural Mouse Laminin (Thermo Fisher Scientific, Waltham, MA, USA), 1uM AraC (Sigma Aldrich, St Louis, MO, USA) and 1 µg/ml Doxycycline hyclate. Half-medium was changed every other day.

### Immunofluorescence and microscopy in human cell lines

HeLa cells were seeded on 8-well glass slides (Millipore). Twenty-four hours post transfection for overexpression or 72 hours post transfection for siRNA knockdown, cells were stressed with 500 μM sodium arsenite (Sigma-Aldrich) for times as indicated in text and legends. Cells were then fixed with 4% paraformaldehyde (Electron Microscopy Sciences), permeabilized with 0.5% Triton X-100, and blocked in 5% bovine serum albumin (BSA). Primary antibodies used were mouse monoclonal anti-FLAG (M2, F1804; Sigma), goat polyclonal anti-eIF3η (sc-16377; Santa Cruz Biotechnology), rabbit monoclonal anti-hnRNPH2 (ab179439; Abcam), mouse monoclonal anti-G3BP (BD Biosciences; 611126), and mouse monoclonal anti-hnRNPA1 (Millipore; 05- 1521). For visualization, the appropriate host-specific Alexa Fluor 488, 555, or 647 (Invitrogen) secondary antibody was used. Slides were mounted using Prolong Gold Antifade Reagent with DAPI (Life Technologies). Images were captured using a Leica TCS SP8 STED 3X confocal microscope (Leica Biosystems) with a 63x objective. Fluorescent images were subjected to automated compartmentalization analysis using CellProfiler software (Broad Institute). Cells were segmented using DAPI and eIF3η channels to identify the nucleus and cytoplasm.

Integrated intensity of nucleus, cytoplasm, and cells were measured. Percent cytoplasmic signal was calculated with the integrated cytoplasmic signal over the integrated cell signal.

### Immunoprecipitation and Western blot analysis in cell lines

Cell lysates were prepared by lysing cells in buffer containing 20 mM phosphate buffer pH 7.5, 150 mM NaCl, 0.2% Triton X-100, and 10% glycerol with complete protease inhibitor cocktail (Clontech Laboratories). Cells were incubated on ice for 20 minutes before centrifugation at 14,000 rpm at 4°C. The resulting supernatant was pre-treated with EZview Red Protein A agarose beads (P6486; Sigma) for 45 minutes to reduce the likelihood of nonspecific binding to the agarose, and the beads were removed. EZview Red Anti-FLAG M2 agarose beads (F2426; Sigma) were then added to the pre-treated lysates and incubated at 4°C for 2 hours. The agarose beads were washed three times with buffer above to remove any remaining nonspecific binding. Samples were eluted with FLAG peptide (F3290; Sigma) at a final concentration of 100 μg/ml for 30 minutes at vortex setting 5 (Scientific Industries) at 4°C. Samples were boiled in 1x LDS sample buffer (Thermo Fisher). Samples were resolved by electrophoresis on NuPAGE Novex 4-12% Bis-Tris gels (Invitrogen). Gels were transferred to nitrocellulose using an iBlot 2 gel transfer device (Thermo Fisher) and blocked in 5% BSA. Primary antibodies used were rabbit polyclonal anti-FLAG (F7425; Sigma) and mouse monoclonal anti-Kapβ2 (ab10303; Abcam). Blots were subsequently incubated with IRDye fluorescence antibodies (LI-COR) and protein bands were visualized using the Odyssey Fc system (LI-COR) and Image Studio (LI-COR). Bands were quantified by densitometry in ImageJ (NIH).

### Pulldown assays for Kapβ2 binding to immobilized GST-hnRNPH2 peptides

*E. coli* (BL21) transformed with pGEX-4TT3 plasmids expressing GST-hnRNPH2 proteins were grown in 35 ml LB with 100 μg/ml ampicillin to OD600 0.6. Protein expression was then induced with 0.5 mM isopropyl-β-d-1-thiogalactoside (IPTG) for 5 hours at 37°C. Cells were harvested by centrifugation, resuspended in lysis buffer (50 mM Tris pH 7.4, 150 mM NaCl, 1 mM EDTA, 2 mM DTT, 15% glycerol, and protease inhibitors), lysed by sonication, the lysate centrifuged, and supernatant containing GST-hnRNPH2 proteins added to Glutathione Sepharose 4B (GSH; GE Healthcare) beads. The beads with immobilized GST-hnRNPH2 proteins were washed with lysis buffer. 50 μl beads containing ∼60 μg immobilized GST-hnRNPH2 proteins were incubated with 8 μM Kapβ2 in 100 μl total volume for 30 minutes at 4°C and then washed three times with 1 ml lysis buffer. Proteins bound on the beads were eluted by boiling in SDS sample buffer and visualized by Coomassie staining of SDS-PAGE gels.

### Generation of *Hnrnph2* mutant and knockout mice

gRNA was in vitro transcribed using MEGAshortscript T7 kit (Life Tech Corp; AM1354), and the template PCR amplified using the following primers:

Forward: 5′ - CCTTAATACGACTCACTATAGGGCTCATGACTATGCAGCGCCGTTTTAGAGCTAGAAATAGC-3′

Reverse: 5′- AAAAGCACCGACTCGGTGCCACTTTTTCAAGTTGATAACGGACTAGCCTTATTTTAACTTGCTATTTCT AGCTCTAAAAC-3′

The resulting PCR products contained the T7 promoter, gRNA sequence, and tracrRNA (5′- CCTTAATACGACTCACTATAGGGCTCATGACTATGCAGCGCCGTTTTAGAGCTAGAAATAG CAAGTTAAAATAAGGCTAGTCCGTTATCAACTTGAAAAAGTGGCACCGAGTCGGTGCTTTT-3′). Synthetic single-strand DNA was used as mutation donor. Donor DNA sequences are shown below.

P209L: 5′- ACAAGGAAAGAATAGGGCATAGGTACATCGAAATCTTCAAGAGTAGCCGAGCTGAAGTCCGAACCCACTATGATCCACCTAGAAAGCTCATGACTATGCAGCGCCCGGGTCTTTACGATAGGCCAGGGGCTGGAAGAGGGTATAATAGCATTGGCAGAGGAGCCGGGTTTGAAAGAATGAGGCGGGGTGCCTATGGTGGA-3′

R206W: 5′- AACACAAGGAAAGAATAGGGCATAGGTACATCGAAATCTTCAAGAGTAGCCGAGCTGAAGTCCGAACCCACTATGATCCACCTAGAAAGCTCATGACTATGCAGTGGCCGGGTCCTTACGAT AGGCCAGGGGCTGGAAGAGGGTATAATAGCATTGGCAGAGGAGCCGGGTTTGAAAGAATGAGGCGGGGTGCCTATGGT-3′

The gRNAs, Cas9 mRNA, and ssDNA were co-microinjected into C57BL/6J zygotes at 25, 25, and 10 ng/μl respectively. Seven mice with P209L and 11 mice with R206W mutations were identified by PCR (5′-GACACTGCCAGTGGACTTTC-3′ and 5′-TGCTCTGGAAACTGGACCCA-3′) followed by sequencing (5′-TGCTCTGGAAACTGGACCCA-3′). These potential founders were crossed with WT C57BL/6J mice to confirm transmission of the mutation. Resulting progeny carrying the mutations were tested for possible off-target effects as predicted by the Wellcome Sanger Institute Genome Editing Off-Target by Sequence tool (50). Of the 61 predicted off-target sites (1: 0, 2: 0, 3: 9, 4: 52), all nine 3-nucleotide mismatch sites were tested by high-resolution melt analysis. All but one of the lines tested showed no off-target effects at these sites (**Supplemental Figure 4**). One line gave variant calls on all 9 sites, which was attributed to low DNA concentration of the sample. Nevertheless, this line was discarded. One line of each mutation (P209L, R206W, and KO) was chosen for phenotyping and heterozygous mutant or KO females bred to C57BL/6J males to maintain the genetic background. Subsequent generations were genotyped by Transnetyx automated real-time PCR (Transnetyx).

### Breeding of experimental cohorts

For most experiments, heterozygous mutant females were bred to WT males to generate heterozygous mutant or KO females (*Hnrnph2^R206W/X^*, *Hnrnph2^P209L/X^*, *Hnrnph2^KO/X^*), hemizygous mutant or KO males (*Hnrnph2^R206W/Y^*, *Hnrnph2^P209L/Y^*, *Hnrnph2^KO/Y^*), WT females (*Hnrnph2^X/X^*), and WT males (*Hnrnph2^X/Y^*). In addition, for some experiments heterozygous mutant females were bred to hemizygous mutant or KO males to generate homozygous mutant or KO females (*Hnrnph2^R206W/R206W^*, *Hnrnph2^KO/KO^*), heterozygous mutant or KO females (*Hnrnph2^R206W/X^*, *Hnrnph2^KO/X^*), hemizygous mutant or KO males (*Hnrnph2^R206W/Y^*, *Hnrnph2^KO/Y^*), and WT males *Hnrnph2^X/Y^*. We note that this cross could not be performed in the *Hnrnph2*^P209L^ line, as very few *Hnrnph2^P209L/Y^* males survived until sexual maturity (6-8 weeks). All experiments were performed on generation F3 or later. Animals were group housed under standard conditions.

### Mendelian inheritance and survival up to 8 weeks

All pups born and genotyped (samples collected from live pups at P2-P7 and from pups found dead before P2-P7 sample collection) in the colonies from April 2018 to March 2021 were included in calculation of genotype ratios. All pups born and genotyped during this time were also included in survival analyses, except for mice used in cohort 2 (audiogenic seizure cohort) and cohort 3 (µCT and imaging cohort).

### Behavioral phenotyping and long-term survival

Experimental cohort 1, consisting of male hemizygous mutants or KOs, female heterozygous mutants or KOs, and WT littermate controls, were first subjected to an observational test battery at 8 weeks old. This was followed by more specific and sensitive tests of motor and sensory function at 8-9 weeks and 10-12 weeks, respectively. These mice were also weighed weekly from 3 to 8 weeks, then again at 6 months and every 6 months thereafter and followed for survival.

A slightly modified protocol of the EMPReSS (European Mouse Phenotyping Resource for Standardized Screens) version of SHIRPA (SmithKline Beecham, Harwell, Imperial College, Royal London Hospital, Phenotype Assessment) level 1 observational test battery was used (27). Briefly, mice were observed undisturbed in a clear viewing jar for activity, tremor, palpebral closure, coat appearance, skin color, whisker appearance, lacrimation, defecation, and urination. Mice were then moved to an arena and the following parameters scored: transfer arousal, locomotor activity, gait, pelvic elevation, tail elevation, startle response, touch escape and righting reflex. Thereafter, mice were held by the tail and scored for positional passivity, trunk curl, limb clasping, and visual placing. After placement on a wire mesh grid, mice were assessed for corneal reflex, pinna reflex, whisker orienting reflex, toe pinch response, and negative geotaxis. Lastly, contact righting response when place in a tube and rolled upside down was tested, and any evidence of biting and excessive vocalization noted. The data were quantified using a binary scoring system as previously described (51). A normal behavior received a score of 0 and an abnormal behavior received a score of 1, enabling a global abnormality score to be determined for each mouse, with a higher score corresponding to a greater degree of abnormality. In addition, scores were also generated for specific functions including motor, sensory, neuropsychiatric, and autonomic function (28).

Rotarod analysis was performed on an accelerating rotarod apparatus (IITC Life Science) using a 2-day protocol. Mice were trained on the first day with one session set at 4 rpm for 5 minutes. The following day, rotation speed was set to accelerate from 4 to 40 rpm at 0.1 rpm/s, mice were placed on the apparatus, and the latency to fall was recorded for four separate trials per mouse. Mice were given a 15-minute rest period between each trial. Grip strength was measured using a grip strength meter (Bioseb) as grams of force for all 4 paws for each mouse in six repeated measurements. The beam walking test was performed using a 2-day, multi- beam protocol (52). Briefly, on day 1 mice were trained to walk across an elevated 12-mm square beam to reach an enclosed goal box. On day 2, mice received one trial each on a 12- mm square beam, a 6-mm square beam, and a 12-mm round beam, and latency to cross, number of hind paw slips, and number of falls recorded. A custom neurological scoring system was also used, where a score of 0 was given if the mouse was unable to traverse the beam in 60 s, 1 if a mouse traversed the entire beam by dragging itself with its front paws (hind paws remain in contact with the side of the beam at all times), 2 if a mouse was able to traverse the beam with some hind paw stepping on top of the beam before starting to drag itself with its front paws, 3 if a mouse was able to traverse the entire beam with hind paw stepping, but placed its hind paws on the side of the beam at least once (no dragging with front paws), and 4 if a mouse was able to traverse the entire beam with hind paw stepping and never placing its hind paws on the side of the beam. In the wire hang test, mice were placed onto a wire cage top, which was then inverted and elevated above a clean cage, and latency to fall (up to 120 s) recorded. For gait analysis, the front and hind paws of each animal were dipped in red and blue paint (water-soluble and non-toxic), respectively. The animal was then placed in a 70-cm long tunnel lined on the bottom with Whatman filter paper, the entrance sealed, and animal allowed to walk through one time. Footprints were scanned and analyzed with Image J for stride length, fore- and hind base width, and overlap (53).

Experimental cohort 2, consisting of male hemizygous mutants or KOs, female heterozygous mutants or KOs, female homozygous mutants or KOs, and WT littermate controls, were tested for audiogenic seizure susceptibility in a clear acrylic box (30 x 30 x 30 cm), with a 6” red fire bell mounted to the underside of a removable lid, and connected to a standard GraLab timer. The bell consistently produced 120-125 dB sound as measured from inside the closed box using a digital sound level meter. At P21, mice were removed from their home cage one by one just before testing, put into a clean holding cage, and moved to the testing room.

Mice were then transferred to the audiogenic seizure chamber and allowed to explore the box for 15 s before the bell was turned on for 60 s. The intensity of the response (seizure severity score) was categorized as 0 for no response or slight startle, 1 for wild running, 2 for clonic seizures, 3 for tonic seizures, and 4 for respiratory arrest (54).

Experimental cohort 4, consisting of hemizygous R206W males and WT littermate controls, were subjected to a battery of tests over 4 weeks to assess learning and memory, emotional behaviors, and social behaviors. To reduce any potential carryover effects, tests were run in the order of least invasive to most invasive, based on the sensitivity of tests on previous handling, and stress induced by each test (55). Tests were conducted with a 1-2 day interval, which has been shown to have little impact on performance compared to 1 week inter-test intervals (56). Starting at 8 weeks old, experimentally naïve mice were tested for anxiety in the elevated plus maze, anxiety and locomotion in the open field test, visual recognition memory in the novel object recognition test, spatial working memory in the Y maze spontaneous alternation test, social preference in the three-chamber social interaction test, spatial learning and memory in the Morris water maze, and repetitive, compulsive-like behavior in the marble burying test.

For all tests, mice were moved to the test room, kept behind a room divider, and allowed to acclimatize undisturbed for 30 minutes before starting tests. At the end of a test, mice were placed in a holding cage until all mice from a home cage completed testing, before being returned to the home cage. During all tests, the investigator remained out of sight behind a room divider. Behavioral parameters were recorded and analyzed using ANY-maze automated activity monitoring system and software (v7.2). To facilitate automated tracking and reduce stress on the animals, all tests were conducted under low (30-50 Lux), indirect lighting. Mazes, objects (novel object recognition test) and wire cages (three-chamber social interaction test) were thoroughly cleaned with 70% vol/vol ethanol after each test and allowed to dry before starting the next test.

In the elevated plus maze (Stoelting), mice were placed in the center facing an open arm away from the investigator and allowed to explore the maze undisturbed for 5 minutes (57).

Parameters including total distance traveled, mean speed, open and closed arm entries and time, and center entries and time were recorded and analyzed for the total test time, as well as temporally across the session to assess any potential differences in habituation to novelty and aversive learning (33, 34). The animal’s entire area was used to score arm entries and exits, with at least 80% of the animal needed for an arm entry to occur, and mice were considered to be in the center zone if not in any arm. Mice that fell off the maze were excluded from the analysis.

The open field test was run according to the protocol by Seibenhener and Wooten (58).

Briefly, mice were placed in the center of a 40 cm x 40 cm open field arena (Stoelting) facing away from the investigator and allowed to explore undisturbed for 20 minutes. Parameters including total distance traveled, mean speed, time in the center zone (defined as the center 24 cm x 24 cm of the maze) were recorded and analyzed for the total test time, as well as temporally across the session to assess any potential differences in habituation to novelty (59). The animal’s center was used to score zone entries and exits.

The novel object test protocol used was based on those published by Leger et al. (60) and Lueptow (61) and included a familiarization phase and test phase. As the test was conducted in the same open field arena that was used in the open field test a few days prior, a habituation phase was not included. During the familiarization phase, mice were placed in the center of a 40 cm x 40 cm open field arena (Stoelting) containing 2 identical novel objects (either white wooden balls with flat bottoms, or grey wooden squares; Stoelting) and allowed to explore undisturbed for 10 minutes. For the testing stage (24 hours after familiarization), one of the objects was switched out for a novel object. The pair of objects used and the position of the novel object (top left vs bottom right) was randomized between mice and groups. Data analysis was limited to the time needed to reach 20 seconds of total object exploration time. Mice that failed to reach 20 seconds of total object exploration time were excluded from the analysis.

Parameters including total distance traveled, mean speed, time taken to reach 20 seconds of total exploration time, and time exploring each object were recorded and analyzed. Object exploration was defined as the animal’s head being within 20 mm of an object and oriented toward the object (orientation angle of 60°). Instances where mice climbed on top of the objects (animal’s center is inside the object) were not considered to be object exploration. An object discrimination index was calculated for the familiarization stage (top left vs bottom right object) and the test stage (novel vs known object) by subtracting the time exploring the bottom right/known object from the time exploring the top left/novel object, divided by the total object exploration time (20 seconds).

In the Y maze (Stoelting), mice were placed in a distal part of an arm, facing away from the investigator and allowed to explore the maze freely for 8 minutes. Parameters including total distance traveled, mean speed, number of entries and time spent in each arm and the maze center were recorded and analyzed for the total test time. The animal’s entire area was used to score arm entries and exits, with at least 80% of the animal needed for an arm entry to occur, and mice were considered to be in the center zone if not in any arm. A spontaneous alternation was defined as consecutive entry into 3 different arms on overlapping triplet sets, and the percentage of spontaneous alternations was calculated as the number of spontaneous alternations divided by the total number of arm entries minus 2, multiplied by 100 (62). Mice that climbed out of the maze were excluded from the analysis.

The three-chamber social interaction test consisted of a pre-test, followed the next day by a social preference test and social novelty test, with an inter-test interval of 5 minutes (63). In the pre-test, mice were placed in the center chamber of the sociability cage (Stoelting) facing away from the investigator and allowed 10 minutes to freely explore the cage, which contained crumpled paper balls inside wire enclosures placed in the right and left chambers of the cage.

The next day, the 2 wire cages contained either a wooden block to serve as the non-social stimulus, or an unfamiliar age-, sex-, and background strain-matched WT mouse to serve as the social stimulus. The test mouse was placed in the center chamber of the cage facing away from the investigator and allowed to explore the cage undisturbed for 10 minutes. At the end of the test, the test mouse was removed and placed into a holding cage and the wooden block was replaced by an unfamiliar age-, sex-, and background strain-matched WT mouse to serve as the novel social stimulus. The mouse used as the social stimulus during the social preference test was kept in the wire cage and served as the known social stimulus during the social novelty test. After 5 minutes, the test mouse was again placed in the center chamber of the sociability cage facing away from the investigator and allowed to explore the cage undisturbed for 10 minutes. Parameters including total distance traveled, mean speed, and time investigating each stimulus were recorded and analyzed. Stimulus investigation was defined as the animal’s head being within 25 mm of the wire cage and oriented toward the stimulus (orientation angle of 60°). A social preference index was calculated by subtracting the time spent investigating the non-social stimulus from the time spent investigating the social stimulus and dividing the result by the time spent investigating the social stimulus plus the non-social stimulus. A social novelty index was calculated by subtracting the time spent investigating the known social stimulus from the time spent investigating the novel social stimulus and dividing the result by the time spent investigating the known social stimulus plus the novel social stimulus (63).

The Morris water maze protocol was modified from that of Vorhees and Williams (35). A blue circular plastic pool was used with a diameter of 120 cm and depth of 81 cm along with an adjustable height circular platform (grey for cued trials, clear for training trials) with a diameter of 10 cm (MazeEngineers). The pool was filled with water to approximately 20 cm from the top and allowed to equilibrate to room temperature (approximately 20°C) for at least 2 days. To facilitate automated tracking and reduce visibility of the platform during training trials, 3 bottles of non-toxic white tempera paint were added to the pool. A cued test was first performed during which the pool was surrounded by black room dividers to eliminate any distal room cues and the platform was visible (height was adjusted to just above the surface of the water and a red plastic flag was attached to the grey platform to improve visibility). The cued test consisted of 4 trials performed on a single day, with an inter-trial interval of 10-15 minutes (mice were run in blocks of 10 and all mice completed a trial before the next trial was started). The position of the visible platform was moved for each trial (southeast, northeast, southwest, northwest) and the starting position was alternated between north and west. Mice were gently placed in the water facing the pool wall and given 60 seconds to find and climb on to the visible platform. Mice that failed to find the platform were gently picked up and placed on the platform. Once on the platform, they were allowed to remain there for 15 seconds. At the end of each trial, mice were returned to a heated holding cage while waiting to start the next trial and returned to the home cage at the completion of the 4 trials. Two days later, the mice were subjected to 4 days of training, consisting of 4 trials per day, with an inter-trial interval of 10-15 minutes (mice were run in blocks of 10 and all mice completed a trial before the next trial was started). During training trials, only 1 black room divider was used to hide the investigator from view during the test. A black and white stripe poster on the room divider, as well as objects in the testing room around the pool (e.g., two lamps, wall cabinet) served as distal spatial cues. A clear platform was submerged 1-2 cm below the surface of the water and was not visible under the water with white paint added. A set of semi-randomly selected distal start positions were used, with the platform remaining in the southwest quadrant as previously described (35). Mice were gently placed in the water facing the pool wall and given 60 seconds to find and climb on to the hidden platform. Mice that failed to find the platform were gently picked up and placed on the platform. Once on the platform, they were allowed to remain there for 15 seconds. At the end of each trial, mice were returned to a heated holding cage while waiting to start the next trial and returned to the home cage at the completion of the 4 trials. After 4 days of training, mice were tested for spatial memory in a single probe trial, during which the platform was removed from the pool. Mice were gently placed in the water facing the pool wall at a location not used during training, directly across from the previous location of the platform (northeast), and removed after 30 seconds. Parameters including total distance traveled, mean swim speed, time in each quadrant, latency to reach the platform (cued and training trials), percentage of time in the thigmotaxis zone (within 10 cm of the pool wall), cumulative distance from hidden platform (training trials), platform location crossings (probe trial), and mean distance from platform location (probe trial) were recorded and analyzed. Data for the training trials are averaged across 4 trials per day and plotted as block means. The animal’s center was used to score zone entries and exits.

The marble burying test was performed according to the protocol by Angoa-Perez et al. (64). Standard polycarbonate rat cages were filled with fresh mouse bedding to a depth of 5 cm and the bedding surface leveled. Twenty standard glass toy marbles of assorted styles and colors were placed gently on the surface of the bedding in 5 rows of 4 marbles each. A single mouse was placed in a cage away from the marbles, the cage was covered with a filter-top lid, and the mouse allowed to remain in the cage undisturbed for 1 hour. After the test, the mouse was removed, taking care not to move any marbles, and returned to its home cage. The number of marbles buried (at least two-thirds of surface area covered by bedding) was counted.

### EEG implantation

EEG/EMG headstage (Model 8431-SM, Pinnacle Technology) implantation was completed according to the manufacturer’s instructions unless otherwise noted. In brief, mice were anesthetized using isoflurane vapors at 3% and maintained during implantation at 1.5%. A small midline incision was made, exposing the skull. Six holes were made in the skull using a 23- gauge needle; following this, each hole had a screw with a wire lead attached placed inside.

Dental cement (DuraLay, Reliance Dental Manufacturing) was used to cover the screws, leaving the wires exposed. The headstage was then placed on top of the dried dental cement and EMG leads were inserted into the nuchal muscles via a pocket created using forceps. Headstage wires were then connected to the lead wire from the screw using wire glue (Anders Products) and allowed to dry completely before covering them with dental cement. Animals were then given a post-operative analgesic injection (meloxicam, 2 mg/kg) and mush food for the next three days to ease recovery. Animals were given at least one week to recover prior to data collection.

### EEG data collection and analysis

EEG/EMG collection was synchronized with video recording using Pinnacle Seizure Acquisition Software (v2.1.0). Data was obtained using the Pinnacle Acquisition System with a 8406- SE31M pre-amplifier at a sampling rate of 2000 Hz with a high and low bandpass filter at 0 and 500 Hz, respectively. Representative 1-hour time spans were selected in an unbiased manner (12am-1am and 6:30am-7:30am for the dark and light phases, respectively) and were analyzed blind to genotype for spectral power and epileptiform activity. All analyzed data was recorded after at least 2 hours of acclimation to the recording arena and connection to the wiring tether.

Electrical activity was converted to frequency using a fast Fourier transformation (FFT) algorithm using the Hann method. Pinnacle Sirenia Seizure Pro (v2.2.5) software was used for spectral power analysis at individual frequencies and delta (0.5-4 Hz), theta (5- Hz), alpha (9-13 Hz), and beta (14-30 Hz) bands. Spectral power data were used in combination with video recordings to rule out motion-based artifacts and to manually quantify the percentage of time spent in epileptiform activity. Spikes that were greater than 2x the baseline amplitude and <200 ms in duration and showing polyspikes (defined as spikes crossing baseline more than two times) were considered epileptiform. The “time spent” satisfying criterion was summed across the entire hour and calculated as the percentage of time spent exhibiting the phenotype. The lambdoid lead EEG waveform and video from 12am to 1am was watched and scored to detect any absence-like (behavioral arrest during EEG spiking) or motor (behavioral changes similar to myoclonic jerks, tonic-clonic action, wild running, or postural loss) seizures.

### In vivo MRI and µCT

Experimental cohort 3, consisting of male hemizygous mutants or KOs, female heterozygous mutants or KOs, and WT littermate controls, were imaged at the Center for In Vivo Imaging and Therapeutics at St. Jude Children’s Research Hospital using micro-computed tomography (µCT) and magnetic resonance imaging (MRI) at 6 and 24 weeks of age. The µCT was performed on a Siemens Inveon PET/CT system (Siemens) at 88-µm resolution, and the MRI was performed on a Bruker Clinscan 7T MRI system (Bruker Biospin MRI GmbH). MRI was acquired with a mouse brain surface receive coil positioned over the mouse head and placed inside a 72-mm transmit/receive coil. After the localizer, a T2-weighted turbo spin echo sequence with variable flip-angle echo trains was performed in the coronal orientation (TR/TE = 2500/114 ms, matrix size = 192 × 192 x 104, resolution = 0.12 x 0.12 x 0.12 mm, number of averages = 4). Prior to scanning, mice were anesthetized in a chamber (3% isoflurane in oxygen delivered at 1 L/min) and maintained using nose-cone delivery (1-2% isoflurane in oxygen delivered at 1 L/min).

Animals were provided thermal support using an inbuilt electronic heating pad (µCT) or a heated bed with warm water circulation (MRI) and a physiological monitoring system to monitor breath rate. After imaging, animals were allowed to recover on a heating pad.

Morphometric analysis was performed on the µCT images to identify group differences in skull shape. Linear measurements of 11 key craniofacial parameters (19) were performed manually on µCT slices using Inveon Research Workplace software (IRW 4.2, Siemens). This was followed by automated imaged-based shape analysis using a population-level atlas of the *Mus musculus* craniofacial skeleton (20). Briefly, the head was extracted from the whole-body µCT images using an iterative search and best-match algorithm. The µCT atlas (https://github.com/muratmaga/mouse_CT_atlas) was then aligned to native space images using a first pass affine transform, followed by a non-linear warping. The calculated transform was then applied to a set of 51 previously identified landmarks and the coordinates for the landmarks in native space were extracted. Processing steps were performed using the ANTS software package (https://github.com/ANTsX/ANTsPy). All alignment results were visually inspected by at least 2 raters. The Euclidean distance between each point was calculated and used for subsequent analysis. First, we performed pairwise comparisons of linear distances between all 51 landmarks. Next, we performed Euclidean distance matrix analysis (EDMA), a geometric morphometric approach enabling the quantification and comparison of shape in three dimensions (21). For global EDMA analysis all 51 landmarks were included, whereas the regional EDMA analysis was performed on a subset of landmarks that summarize regions with specific embryonic tissue origins, further divided into anatomically relevant subsets including palate, midface, and nasal regions (23). To account for overall difference in size, both the global and regional EDMA analyses were scaled to centroid size (calculated as the square root of the sum of squared distances of all landmarks from their centroid), which is a common proxy for overall size in geometric morphometric analyses (22).

Brain parcellation and volumetrics were performed to investigate group differences in total and regional brain volumes. We used the DSURQE atlas (25), which contains 336 cortical, white matter, subcortical, and CSF defined regions. The DSURQE anatomical image was first downsampled to 120-μm isotropic resolution to satisfy the Nyquist criteria of our image resolution and reduce computational time for fitting. The acquired T2 images were preprocessed, including skull-stripping and intensity normalization. The images were then aligned to the atlas by a first-pass affine registration, followed by a non-linear warping. The inverse warping was applied to the labeled atlas to bring all labeled areas into native space. All image processing steps were performed using the AFNI software package (https://afni.nimh.nih.gov/). The volume (number of voxels times native resolution) of each labeled area from the atlas was extracted for subsequent analysis. The results of the inverse warping were quality checked by visual inspection by at least 2 raters. Cases with poor alignment (17 out of a total of 140) were removed from the final volumetric analysis.

### Mouse histology and immunofluorescence

For confirmation of hydrocephalus, mice were anesthetized by isoflurane inhalation and transcardially perfused with 10% neutral buffered formalin (NBF) (mice flagged for domed heads) or postfixed in 10% neutral buffered formalin (mice found dead). Heads were decalcified, paraffin-embedded in the coronal plane, 10 4-µm step sections (every 50 µm) cut, stained with hematoxylin and eosin (H&E), and reviewed by a veterinary pathologist.

Brains from experimentally naïve male hemizygous mutants or KOs and male WT littermate controls were harvested at 8 weeks (*Hnrnph2^R206W^* and KO) or 3 weeks (*Hnrnph2^P209L^*) of age for histology and immunofluorescence. Briefly, mice were anesthetized by isoflurane inhalation and transcardially perfused with 10% NBF, the brain dissected from the skull and cut in half on the sagittal plane, processed for paraffin embedding, and cut at 10 µm. Sections were stained with H&E and Luxol fast blue-cresyl violet (LFB-CV) to ascertain overall morphology and myelination. In addition, IHC was performed using antibodies against neurons (NeuN; 2367, Cell Signaling Technology), astrocytes (GFAP; Z0334, DAKO), microglia (IBA1; CP290A, BioCare Medical), and oligodendrocytes (OLIG2; ab109186, Abcam). Sections were deparaffinized, followed by heat-induced epitope retrieval (HIER) with appropriate buffer (AR9640, Leica; 950- 500 or 760-107, Roche), incubation with primary antibodies and Bond Polymer Refine Detection with DAB (DS9800, Leica), or incubation with OmniMap Rabbit HRP antibody (760-4311, Roche) and ChromoMap DAB (760-159, Roche). Lastly, sections were counterstained with hematoxylin and, if needed, post-counterstained with Bluing Reagent (760-2037, Roche), before being coverslipped. Sections were reviewed by a veterinary pathologist and immunoreactivity quantified using HALO image analysis platform (Indica Labs). The number of NeuN-positive cells was quantified using QuPath software (65). Briefly, visual, somatosensory, and somatomotor cortices were manually annotated according to the Allen mouse brain atlas, and QuPath’s positive cell detection function applied. The number of NeuN-positive neurons were expressed as number of cells per mm^2^.

To assess the expression of hnRNPH2, immunofluorescence was performed using an N-terminal hnRNPH2 antibody (ab179439, Abcam) or a C-terminal hnRNPH2 antibody (ab181171, Abcam), as well as antibodies for neuronal nuclei (NeuN; ab104224, Abcam), or neuronal cytoplasm and processes (beta III tubulin; ab78078, Abcam). To assess cortical cytoarchitecture, immunofluorescence was performed using antibodies against SATB2 (sc- 81376, Santa Cruz Biotechnologies), which is broadly expressed in upper layer (II-IV) neurons as well as in subpopulations of deep layer (V-VI) neurons, CTIP2 (ab18465, Abcam), which is expressed exclusively in a subpopulation of layer V neurons, and FOXP2 (HPA000382, Atlas Antibodies), which is expressed in layer VI neurons. Sections were deparaffinized, followed by HIER using Universal Antigen Retrieval Reagent (Roche, CTS015), permeabilization in PBS containing 2% Triton X-100, and treatment with TrueBlack Lipofuscin Autofluorescence Quencher (23007, Biotium). Thereafter, slides were blocked in PBS containing 4% bovine serum albumin (A2153, Sigma-Aldrich) and 2% normal goat serum (S-1000, Vector Laboratories), and incubated with primary antibodies and species-specific Alexa Fluor secondary antibodies (A32732A11029, A21434, and A21244, Thermo Fisher Scientific). Finally, slides were coverslipped with anti-fade mounting media containing DAPI (P36931, Thermo Fisher Scientific). Fluorescence slide scanning was performed using a Zeiss Axio Scan.Z1 with a Hamamatsu ORCA-Flash4.0 V3 camera using Zeiss ZEN 3.1 software. Images were created with a Zeiss Plan-Apochromat 20X/0.8 objective lens with illumination by Zeiss Colibri.2 LEDs (365 nm, 470 nm, 555 nm) and corresponding filters (Zeiss Filter Set 49, 38 HE, and 43 HE, respectively). For subcellular localization of mutant hnRNPH2, fluorescent imaging was performed using a Yokogawa CSU W1 spinning disk attached to a Nikon Ti2 eclipse with a Photometrics Prime 95B camera using Nikon Elements software. A 60× Plan Apo 1.40NA oil objective was used and Perfect Focus 2.0 (Nikon) was engaged for all captures. Imaging was performed using 405-nm, 488-nm, and 561-nm lasers for DAPI, Alexa Fluor 488, and Alexa Fluor 555, respectively. Image J/Fiji software (66) was used for maximum intensity Z-projection and color image processing (LUT Fire) for visualization of cytoplasmic hnRNPH2 signal. For cortical cytoarchitecture, fluorescently labeled cells were quantified using QuPath software (65). Briefly, rectangular regions of interest were positioned over visual, somatosensory, and somatomotor cortical regions, with each region of interest subdivided into eight equal bins from the pia to the inner border of the cortex (67). QuPath’s positive cell detection function was used to detect all cells using the DAPI channel, followed by application of a single measurement classifier for the remaining channels. The distribution of neurons was expressed as the number of SATB2, CTIP2, and FOXP2 neurons as a percentage of the total number of DAPI-positive cells within each bin.

### Mouse in situ hybridization

Whole embryos and brains of WT C57BL/6J mice were harvested at embryonic day 12.5, 14.5, 16.5 and postnatal day 0, 7, 14, and 56, respectively. Samples were fixed in 10% neutral buffered formalin and in situ hybridization performed with a chromogenic (Fast Red), single-plex BaseScope assay (Advanced Cell Diagnostics) according to the manufacturer’s instructions with custom probes against *Hnrnph1* (BA-Mm-Hnrnph1-3zz-st) and *Hnrnph2* (BA-Mm-Hnrnph2-2zz- st). Slides were scanned on the PANNORAMIC 250 Flash digital scanner (3DHISTECH) and analyzed using HALO image analysis platform according to the RNAscope quantification protocol (Indica Labs). Briefly, cells in a tissue section were grouped into 5 bins based on the number of dots per cell ranging from 0+ to 4+. Clusters were divided by the typical probe signal area to calculate a dot number for the cluster in identified cells of interest. Each sample was evaluated for the percentage of cells in each bin. The H-score for the sample was calculated by totaling the percentage of cells in each bin according to the weighted formula shown below, and a single score was assigned to an entire tissue section based on the average target expression in this tissue. H-scores were provided on a weighted scale of 0–400. The H-score was calculated using the algorithm with the following equation: H-Score = (1 × % *Probe* 1 + *Cells*) + (2 × % *Probe* 2 + *Cells*) + (3 × % *Probe* 3 + *Cells*) + (4 × % *Probe* 4 + *Cells*).

### Mouse Western blots and digital droplet RT-PCR

Brains from experimentally naïve male hemizygous mutants or KOs and male WT littermate controls were harvested at 8 weeks (*Hnrnph2^R206W^* and KO) or 3 weeks (*Hnrnph2^P209L^*) of age for Western blots and ddRT-PCR. In addition, brains of WT C57BL/6J mice were harvested at embryonic day 12.5, 14.5, 16.5 and postnatal day 0, 7, 14, and 56. Brains were removed, cortices dissected out, flash frozen in liquid nitrogen, and stored at -80°C.

For Western blots, samples were subjected to sequential solubility fractionation or nucleocytoplasmic fractionation as previously described (68). Protein concentrations were determined by DC protein assay (5000111, Bio-Rad), and 35 µg RIPA-soluble, 80 µg cytoplasmic protein, or maximum volume nuclear lysate (40 µl) was loaded onto the gel. Electrophoresis was performed using the Bolt Bis-Tris Plus mini gel system (Thermo Fisher Scientific). Gels were transferred to PVDF membranes using the iBlot 2 dry blotting system (Thermo Fisher Scientific), blocked in Odyssey TBS blocking buffer (LI-COR), incubated with primary antibodies against hnRNPH2 (ab179439, Abcam), hnRNPH1 (PA5-50678, Thermo Fisher Scientific), GAPDH (97166, Cell Signaling Technology), or lamin A/C (Cell Signaling Technology, 2032), followed by species-specific IRDye secondary antibodies (925-3221, 925- 68070, LI-COR). Blots were imaged on the Odyssey CLx system and analyzed on Image Studio software (LI-COR).

For ddRT-PCR, samples were treated with RNAlater-ICE (Thermo Fisher Scientific), RNA extracted using the RNeasy Plus Universal Mini Kit (73404, Qiagen), and treated for DNA contamination with the TURBO DNA-free kit (AM1907, Thermo Fisher Scientific). 10 ng RNA was used with a one-step RT-ddPCR advanced kit for probes (1864021, Bio-Rad), together with the following assays: Mouse *Gapdh* Primer Limited (Mm99999915_g1, Thermo Fisher Scientific), Mouse *Rpp30* (dMmuCPE5097025, Bio-Rad), Mouse *Hnrnph1* (Mm00517601_m1, Thermo Fisher Scientific), and Mouse *Hnrnph2* (Mm01340844_g1, Thermo Fisher Scientific).

### Primary cortical neuron culture

Primary culture of cortical neurons was prepared from P1 mice as previously described (69). The cells were plated on 8-well dishes (Lab-Tek chambered cover glass, Nunc, Thermo Fisher Scientific) coated with 10 μg/ml poly-L-ornithine (Sigma Aldrich) at a density of 30 x 10^4^ cells/well and cultured in NeuroCult neuronal plating medium (Stem Cell Technologies) supplemented with B-27 (Thermo Fisher Scientific) and GlutaMAX Supplement (Gibco, Thermo Fisher). On day 5, primary neurons were transitioned to BrainPhys Neuronal Medium (Stem Cell Technologies) and supplemented with B-27 by performing half-medium changes every 3-4 days.

### Magnetofection of primary cortical neurons

On day 6, neurons were transfected using paramagnetic nanobeads (NeuroMag, OZ Biosciences). pVectOZ-GFP (0.4 μg) (OZ Biosciences) was incubated with 0.75 μl NeuroMag beads in 50 μl Opti-MEM (Thermo Fisher Scientific) for 15 minutes and then added dropwise to the cultures. Cells were incubated on top of a magnetic plate (OZ Biosciences) for 15 minutes.

### Neuronal cell staining and imaging

On day 14, neurons were fixed with 4% paraformaldehyde (Electron Microscopy Services) for 20 minutes, permeabilized with 0.1% Triton-X for 10 minutes, and blocked with 5% normal goat serum in TBST for 1 hr. Neurons were then further incubated with primary antibody against MAP2 (Abcam, ab5392, 1:1000), washed 3 times with TBST, incubated with a host-specific Alexa Fluor 647 antibody (Thermo Fisher, A21449, 1:1000) for 2 hours at room temperature, and washed 3 times with TBST and 2 times with PBS. Imaging was performed on a Yokogawa CSU W1 spinning disk attached to a Nikon Ti2 eclipse with a Photometrics Prime 95B camera using Nikon Elements software (v5.21.02). All imaging was performed on a Nikon Plan Apo 60× 1.40 NA oil objective, with Immersol 518 F (Zeiss). After finding neurons positive for both MAP2 and GFP, a tilescan of 592.92 µm x 592.92 µm was taken centered around the soma, with a z-step size of 0.3 µm for a total z-stack size between 15 µm and 40 µm per neuron. Acquisition of the GFP channel was performed using a 488-nm laser at 50% power with 100-ms exposure. Stitching was performed automatically in Nikon Elements.

### Image analysis of dendritic arborization and spines

Stitched images were imported into Imaris (Bitplane v9.9.1), and using the in-built Filaments module, dendrites and dendritic spines originating from the centered soma in each image were segmented. Any dendrite originating from the cell body was considered as a primary dendrite. A dendritic branch point was determined when the original dendrite bifurcated into two or more daughter trees. For assigning branch level, the lowest level was given to the primary dendrites and levels were increased when a branch point was reached. Dendritic full branch depth was calculated by the sum of dendritic full branch points of a given neuron and dendritic full branch level was the maximum value reached by any primary tree on a given neuron. For Sholl analyses, Sholl radii originating from the centroid of the soma were increased at 1-μm intervals. The Sholl intersection profile was obtained by counting the number of dendritic branches at each given distance from the soma and/or averaging them across the entire neuron. Morphometric measurements for dendrite full branch depth, dendrite full branch level, dendrite length, number of Sholl intersections, total number of spines, and spine density (per 10 µm) of the segmentation were extracted and plotted in GraphPad Prism 9.

### RNA sequencing

Total RNA was extracted with an RNeasy Universal Plus Mini kit (Qiagen; 73404). A sequencing library was prepared with a TruSeq Stranded Total RNA Kit (Illumina) and sequenced with an Illumina HiSeq system with 100-bp read length. Total stranded RNA sequencing data were processed by the internal AutoMapper pipeline. Raw reads were first trimmed (Trim-Galore v0.60), mapped to mouse (GRCm38) or human (hg38) genome assembly (STAR v2.7) and then the gene level values were quantified (RSEM v1.31) based on GENCODE annotation (vM22).

We obtained the TPM (transcript per million) counts for genes with TPM greater than one in at least one sample and the gene expression analysis was performed with non-parametric ANOVA using Kruskal-Wallis and Dunn’s tests on log-transformed TPM counts between three replicates of each experimental group, implemented in Partek Genomics Suite v7.0 software (Partek). The expression of a gene was considered significantly different if *P* < 0.05 and log2FC > 0.5 in at least one of the group comparisons. The calculated z-scores of significantly differential expressed genes or log2Rs were plotted using hierarchical clustering in a heat map, using correlation distance measure, implemented in Spotfire v7.5.0 software (TIBCO). For alternative splicing analysis, the aligned and sorted BAM files after STAR alignment were used for AS analysis using rMATS (v4.0.2) (70). A3SS, A5SS, SE, RI, and MXE events were evaluated. Significant AS events were identified while average coverage >10 and delta percent spliced in (ΔPSI) > 0.1.

### Statistics

Significant differences from expected Mendelian inheritance ratios were determined by chi-square tests. The log-rank (Mantel-Cox) test was used to determine significant differences between survival curves and hazard ratios computed by a log-rank approach. Differences in body weight over time were determined by fitting a mixed-effects model (REML) for time, genotype, and time x genotype interaction, followed by Sidak’s multiple comparisons test to compare WT mice to mutants or KOs. For differences in linear craniofacial measurements, we used a two-way ANOVA (line, genotype, line x genotype interaction) followed by Sidak’s multiple comparisons test to compare WT mice to mutants or KOs. For MRI analysis and linear interlandmark distance analysis, Wilcoxon rank sum test was used to compare groups. In the EDMA analysis, biological shapes were compared using an EDMA bootstrap test (21). The global test is based on the pairwise distances in the form matrices, taking the max/min ratio of the distances. This is then done for all the B replicates, which provides the null distribution. The analysis was performed using the R package EDMAinR. Correction for multiple testing was performed using the FDR method. Significance for the incidence of hydrocephalus was determined by Fisher’s exact test. SHIRPA and audiogenic seizure scores were analyzed by aligned ranks transformation (ART) non-parametric two-way ANOVA (line, genotype, line x genotype interaction), followed by Mann-Whitney U test to compare WT mice to mutants or KOs. Differences in all motor tests, optomotor response, and hot plate test were determined by two-way ANOVA (line, genotype, line x genotype interaction) followed by Sidak’s multiple comparisons test to compare WT mice to mutants or KOs. Scent habituation data were analyzed by repeated measures two-way ANOVA (trial, genotype, trial x genotype interaction), followed by Sidak’s multiple comparisons test to compare WT mice to mutants or KOs. For the open field test, differences between mutants and WT controls were determined by unpaired t test when analyzing the total duration of the test, and by repeated measures two-way ANOVA (time bin, genotype, time bin x genotype interaction) followed by Sidak’s multiple comparisons test when analyzing different time bins across the test. Significance for the incidence of falls from the elevated plus maze was determined by Fisher’s exact test. Total distance was analyzed by unpaired t test, and percentage time spent in the maze zones was subjected to two-way ANOVA (zone, genotype, zone x genotype interaction), followed by Sidak’s multiple comparisons test to compare mutants and controls. For time bin analysis across the elevated plus maze test, repeated measures two-way ANOVAs (time bin, genotype, time bin x genotype interaction), followed by Dunnett’s multiple comparison test to time bin 1 (% time in open arms, % time in closed arms) or by Sidak’s multiple comparisons test to compare mutants and controls (% time in center zone) was used. Morris water maze cued and training data were analyzed by repeated measures two-way ANOVA (trial/day, genotype, trial/day x genotype interaction), followed by Sidak’s multiple comparisons test to compare mutants and controls. Probe trial data were analyzed by unpaired t test (mean speed, mean distance from platform location, % time in the thigmotaxis zone, platform location crossings) and two-way ANOVA (quadrant, genotype, quadrant x genotype interaction) followed by Sidak’s multiple comparisons test to compare mutants and controls. Y maze, novel object recognition and three chamber social interaction data were analyzed by unpaired t tests. In the Y maze, the correlation between total arm entries and % spontaneous alternations were evaluated by Pearson’s correlation.

Nuclear hnRNPH2 levels in mouse cortex by Western blot were compared using unpaired t tests. *Hnrnph1* and *Hnrnph2* transcript levels measured by ddRT-PCR were analyzed by two-way ANOVA (line, genotype, line x genotype interaction) followed by Sidak’s multiple comparisons test to compare WT to mutants or KOs. *Hnrnph1* and *Hnrnph2* expression by ddRT-PCR and ISH were analyzed by two-way ANOVA (genotype, developmental time point, genotype x developmental time point interaction) followed by Sidak’s multiple comparisons test to compare *Hnrnph1* levels to *Hnrnph2* levels at each time point, or Tukey’s multiple comparisons test to compare transcript levels between developmental time points for each gene separately. *Hnrnph2* expression by ddRT-PCR in *Hnrnph2* KO mice were analyzed by one-way ANOVA followed by Sidak’s multiple comparisons test to compare WT males to hemizygous KO males, as well as WT females to heterozygous and homozygous KO females. NeuN-positive cell counts were analyzed by two-way ANOVA (line, genotype, line x genotype interaction) followed by Sidak’s multiple comparisons test to compare WT to mutants or KOs. For cortical layer analysis, the percentage of SATB2-, CTIP2- and FOXP2-positive neurons were analyzed by two-way ANOVA (genotype, bin, genotype x bin interaction) followed by Tukey’s multiple comparisons test. A *P* value less than 0.05 was considered significant.

### Study approval

All animal experiments were reviewed and approved by the IACUC of St. Jude Children’s Research Hospital.

## Author contributions

AK, XY, KO, AG, BJWT, HN, and FY designed and performed laboratory experiments. AK, KO, AG, BJWT, JM, AC, YDW, TP, HT, SSZ, YMC, JPT, and HJK analyzed data. AK, JPT, and HJK drafted and revised the manuscript. JPT and HJK supervised the overall study.

## Acknowledgments

We thank members of the St. Jude Center for In Vivo Imaging and Therapeutics, the Department of Diagnostic Imaging, and the Developmental Neurobiology behavior core for assistance with MRI, µCT, and sensory function testing, including Jieun Kim, Young Sang Ryo, Kaitlin Lord, Matthew Scoggins, Silu Zhang, Bogdan Mitrea, Donnie Eddins, and Lauren Beloate. We thank Natalia Nedelsky for editorial assistance. This work was supported by the Howard Hughes Medical Institute and R35NS097974 grants to JPT; R35GM141461, R01GM069909, the Welch Foundation Grants I-1532, support from the Alfred and Mabel Gilman Chair in Molecular Pharmacology, and Eugene McDermott Scholar in Biomedical Research to YMC; and grant F31NS120465 to AG.

## Supplemental Figure Legends

**Supplemental Figure 1.**
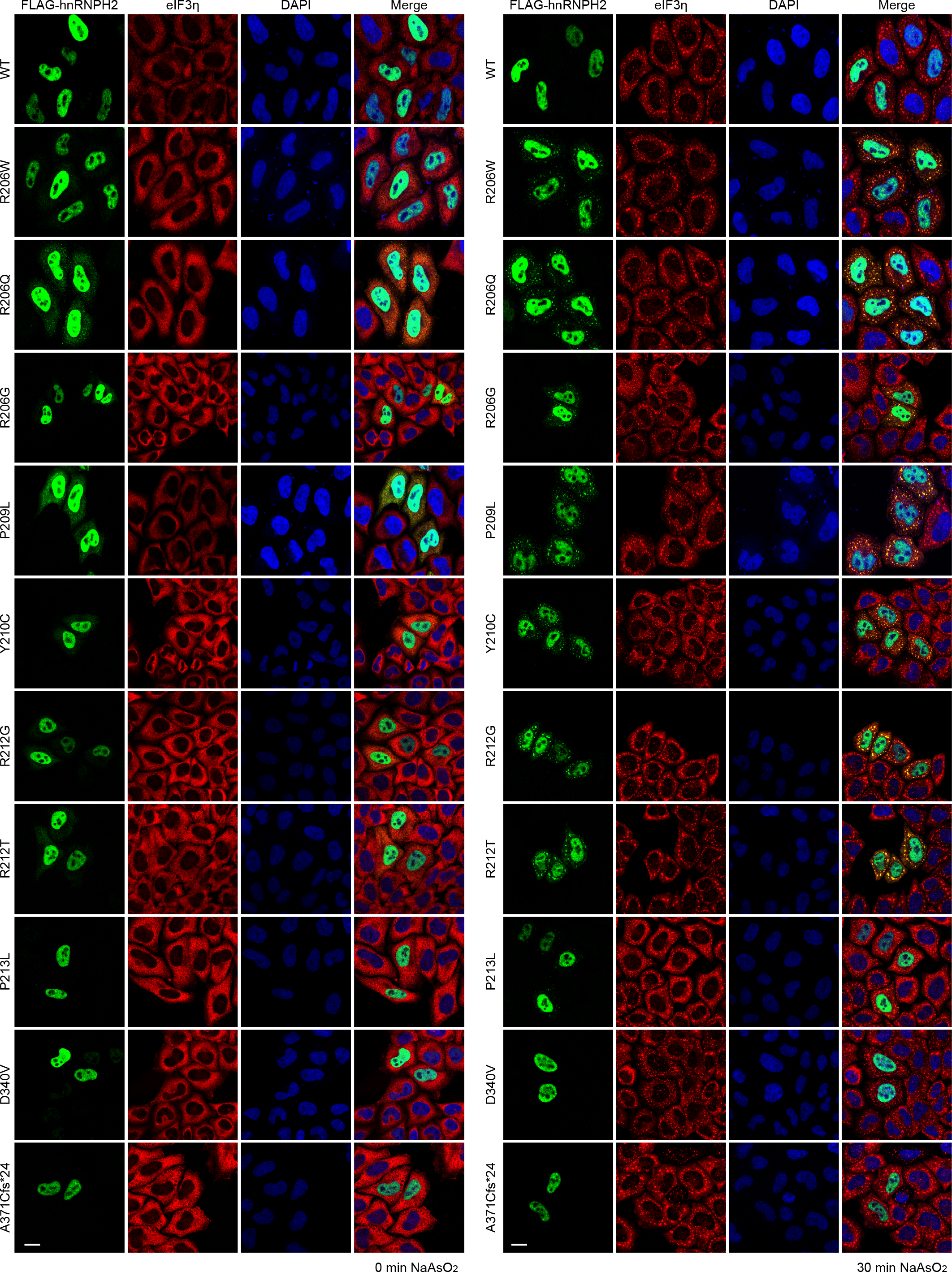
Pathogenic variants alter the nucleocytoplasmic ratio of hnRNPH2 and enhance its recruitment to RNA granules. Intracellular localization of indicated FLAG-tagged hnRNPH2 proteins under basal (left) and stressed (right) conditions in HeLa cells. eIF3η was used as a cytoplasmic and stress granule marker. Scale bar, 10 μm.

**Supplemental Figure 2.**
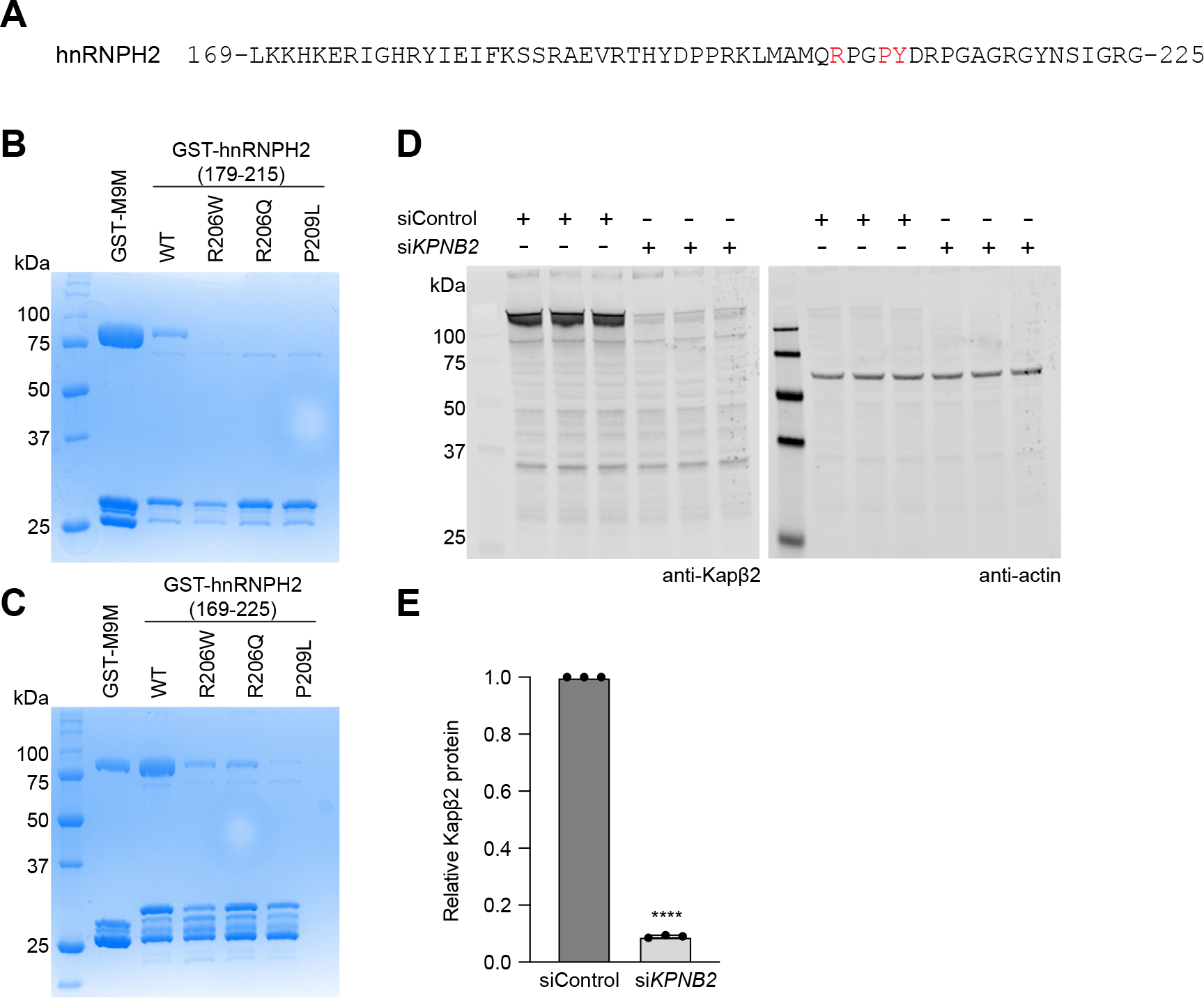
Frameshift variants impair the interaction between hnRNPH2 and its nuclear transport receptor Kapβ2. (**A**) hnRNPH2 amino acid sequence used for GST pulldown experiments. Amino acid residues mutated in patients are in red. (**B** and **C**) Peptides spanning amino acids 179-215 (**B**) or 169-225 (**C**) were fused to the C terminus of GST. Gels show Coomassie Blue staining following GST pulldown of purified GST-hnRNPH2 peptides with Kapβ2. (**D**) Immunoblot showing knockdown of Kapβ2 by si*KPNB2*. Three independently prepared samples were loaded on the gel. (**E**) Quantification of Kapβ2 signal shown in (**D**). Graph shows relative Kapβ2 levels normalized to actin; error bars represent mean ± SD. \*\*\*\**P* < 0.0001 by student’s t-test.

**Supplemental Figure 3.**
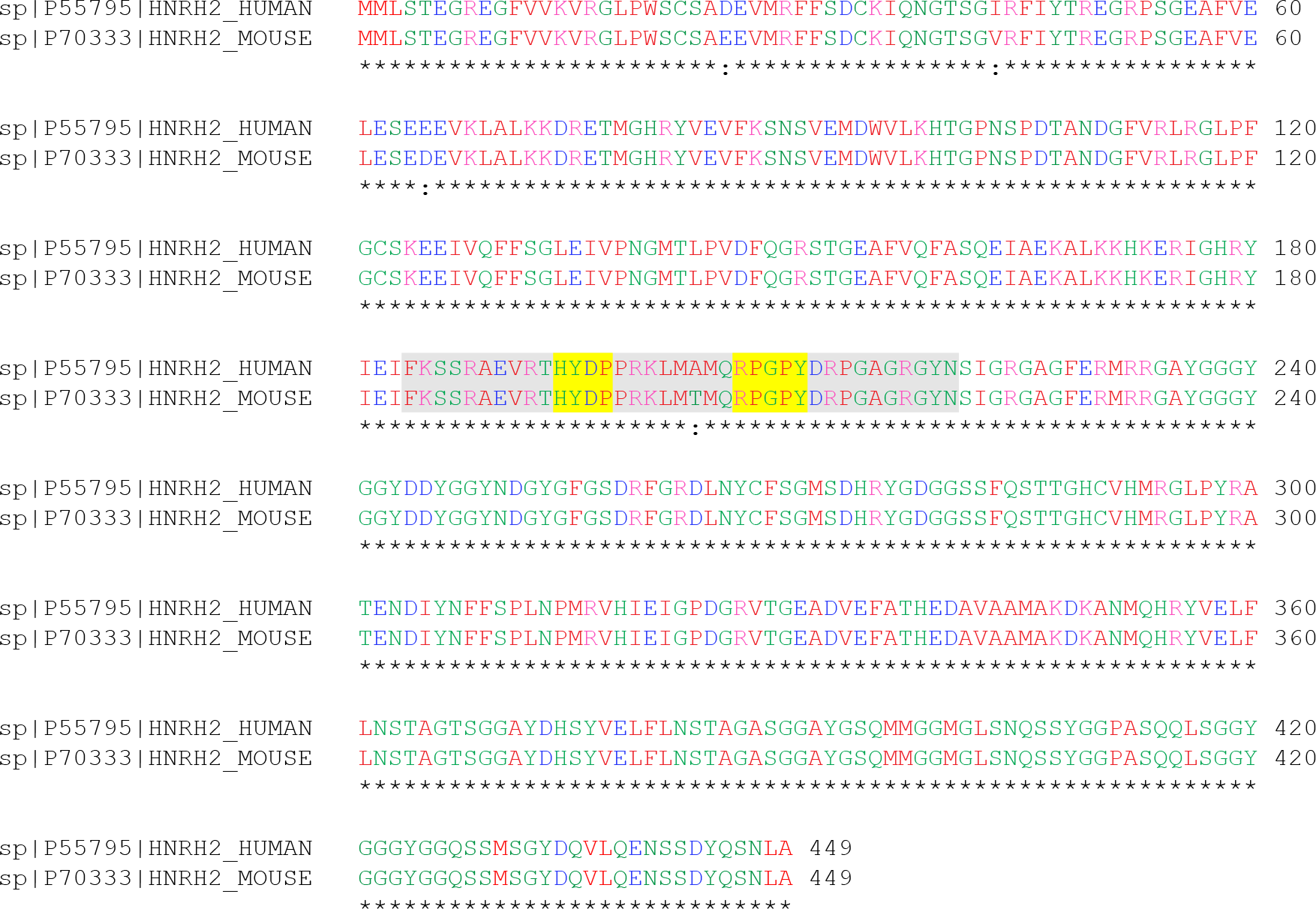
Sequence alignment of human and mouse hnRNPH2 proteins. Human hnRNPH2 and mouse hnRNPH2 are highly conserved. Both proteins are composed of 449 amino acids, of which only 4 amino acids differ. The PY-NLS motif (yellow highlight) within the PY-NLS (gray highlight) is absolutely conserved between the two species.

**Supplemental Figure 4.**
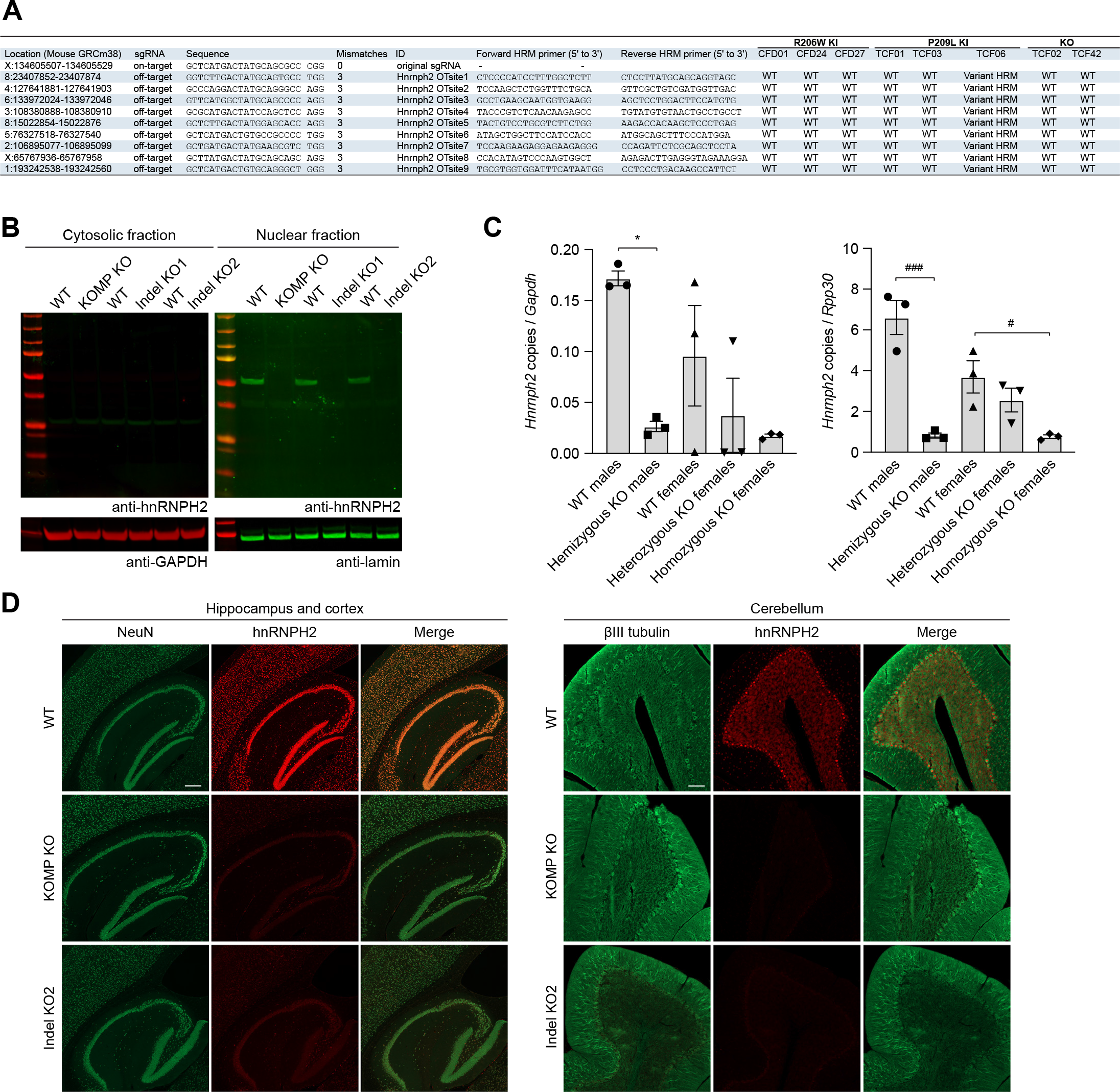
Generation and validation of *Hnrnph2* mutant and KO mouse lines. (**A**) Off-target analysis of knock-in mice generated. Lines CFD01, TCF03, and TCF02 were selected for *Hnrnph2^R206W^*, *Hnrnph2^P209L^*, and *Hnrnph2^KO^* genotypes, respectively, for further experiments. (**B**) Western blot of hnRNPH2 expression in cortex of WT, commercially available *Hnrnph2* KO line (KOMP KO, C57BL/6NJ-*Hnrnph2^em1(IMPC)J^*/Mmjax), and two KO lines we generated (Indel KO1 (TCF42) and Indel KO2 (TCF02)). Indel KO2 (line TCF02) was chosen as the *Hnrnph2* KO line for further experiments. Representative images from n ≥ 3 experiments. (**C**) Expression of *Hnrnph2* transcript by ddRT-PCR in the cortex of *Hnrnph2* KO (Indel KO2 (TCF02)) mice. Normalization to *Gapdh* and *Rpp30* are shown; error bars represent mean ± SEM. \**P* = 0.0121 WT males vs. hemizygous *Hnrnph2* KO males with normalization to *Gapdh*, ^###^*P* = 0.0001 WT males vs. hemizygous *Hnrnph2* KO males with normalization to *Rpp30*, ^#^*P* = 0.0153 WT females vs. homozygous *Hnrnph2* KO females with normalization to *Rpp30,* by one-way ANOVA with Sidak’s multiple comparisons test. n = 3 for all groups. (**D**) Immunofluorescent staining of hnRNPH2 in brain sections of WT, a commercially available *Hnrnph2* KO line (KOMP KO), and a newly generated *Hnrnph2* KO line (Indel KO2). Representative images are shown from n ≥ 3 experiments. NeuN and βIII tubulin were used as neuronal nuclear and cytoplasmic markers, respectively. Scale bar, 200 μm.

**Supplemental Figure 5.**
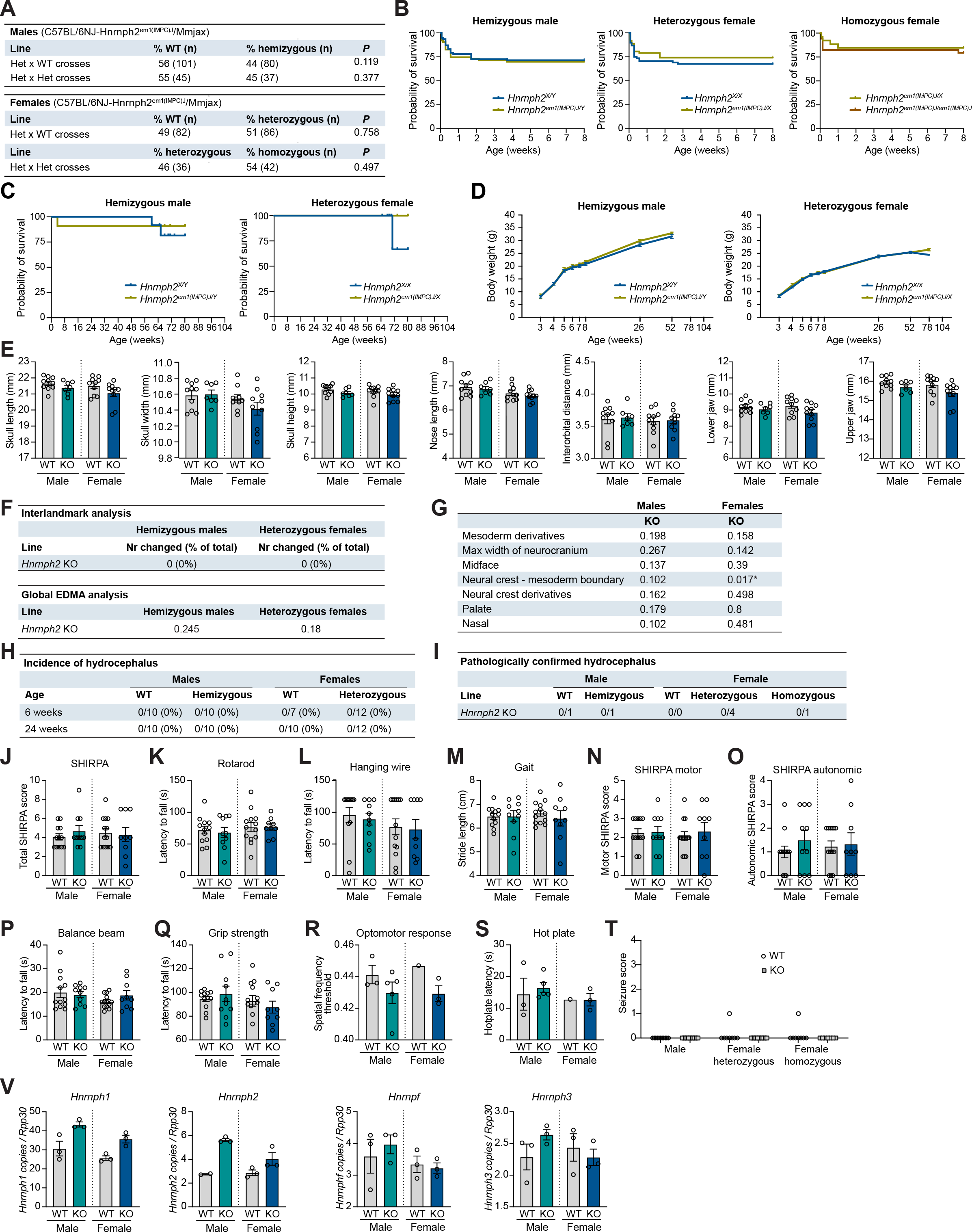
Summary of data from *Hnrnph2^em1(IMPC)J^*/Mmjax (KOMP) KO mice. (**A**) Ratios of genotyped mice organized by sex and breeding strategy. (**B**) Kaplan-Meier survival curves of male and female mice up to 8 weeks of age. *P* = 0.8028 (HR = 1.079), 0.4151 (HR = 0.7729), and 0.5998 (HR = 1.378) for hemizygous male, heterozygous female, and homozygous female, respectively. Group sizes were as follows: *Hnrnph2^em1(IMPC)J/Y^* (n = 63) vs. *Hnrnph2^X/Y^* (n = 77); *Hnrnph2^em1(IMPC)J/X^* (n = 62) vs. *Hnrnph2^X/X^*(n = 68); *Hnrnph2^em1(IMPC)J/em1(IMPC)J^* (n = 34) vs. *Hnrnph2 ^em1(IMPC)J/X^* (n = 26). (**C**) Kaplan-Meier survival curves of male and female mice up to 2 years of age. *P* = 0.6406 (HR = 0.5688) and 0.1573 (HR = undefined) for hemizygous male and heterozygous female, respectively. Group sizes were as follows: *Hnrnph2^em1(IMPC)J/Y^* (n = 11) vs. *Hnrnph2^X/Y^* (n = 12); *Hnrnph2^em1(IMPC)J/X^* (n = 9) vs. *Hnrnph2^X/X^* (n = 13). (**D**) Mean body weight of male and female mice over time. Error bars represent mean ± SEM. Group sizes were as follows: *Hnrnph2^em1(IMPC)J/Y^* (n = 11) vs. *Hnrnph2^X/Y^*(n = 12); *Hnrnph2^em1(IMPC)J/X^* (n = 9) vs. *Hnrnph2^X/X^* (n = 13). (**E**) Linear measurements of key craniofacial parameters in hemizygous male and heterozygous female mice. Error bars represent mean ± SEM. Group sizes for craniofacial analysis by µCT were as follows: *Hnrnph2^em1(IMPC)J/Y^* (n = 7) vs. *Hnrnph2^X/Y^* (n = 10); *Hnrnph2^em1(IMPC)J/X^* (n = 10) vs. *Hnrnph2^X/X^* (n = 10). (**F**) Number of significantly changed linear inter-landmark distances (top) and results of global EDMA analysis (bottom). (**G**) *P* values for regional EDMA analysis for hemizygous males and heterozygous females. (**H**) Incidence of hydrocephalus at 6 and 24 weeks of age. (**I**) Number of mice found dead or flagged for domed heads with pathologically confirmed hydrocephalus. (**J-M**) Characterization of hemizygous male and heterozygous female mice showing (**J**) total SHIRPA abnormality scores, (**K**) latency to fall from rotarod, (**L**) latency to fall from a wire cage top, and (**M**) quantification of stride length. (**N-O**) Subdomain SHIRPA scores for hemizygous male and heterozygous female mice are shown for (**N**) motor function and (**O**) autonomic function. (**P-S**) Characterization of hemizygous male and heterozygous female mice showing (**P**) latency to cross a balance beam, (**Q**) grip strength, (**R**) optomotor response test of visual acuity, and (**S**) hot plate test of pain response. Group sizes for SHIRPA and motor tests were as follows: *Hnrnph2^em1(IMPC)J/Y^* (n = 10) vs. *Hnrnph2^X/Y^* (n = 12); *Hnrnph2^em1(IMPC)J/X^* (n = 9) vs. *Hnrnph2^X/X^* (n = 13). Group sizes for sensory tests were as follows: *Hnrnph2^em1(IMPC)J/Y^* (n = 5) vs. *Hnrnph2^X/Y^* (n = 3); *Hnrnph2^em1(IMPC)J/X^* (n = 3) vs. *Hnrnph2^X/X^* (n = 1). (**T**) Audiogenic seizure severity scores. Group sizes were as follows: *Hnrnph2^em1(IMPC)J/Y^* (n = 11) vs. *Hnrnph2^X/Y^* (n = 11); *Hnrnph2^em1(IMPC)J/X^* (n = 11) vs. *Hnrnph2^X/X^* (n = 8); *Hnrnph2^em1(IMPC)J/X^* (n = 8) vs. *Hnrnph2 ^em1(IMPC)J/em1(IMPC)J^* (n = 7). (**U**) Number of copies of *Hnrnph1*, *Hnrnph2*, *Hnrnpf*, and *Hnrnph3* normalized to *Rpp30* in the cortex of *Hnrnph2^em1(IMPC)J^/*Mmjax hemizygous male and heterozygous female KO mice by ddRT-PCR. n = 3 per group. In all analyses, WT mice are littermate controls. Error bars represent mean ± SEM in graphs J-U.

**Supplemental Figure 6.**
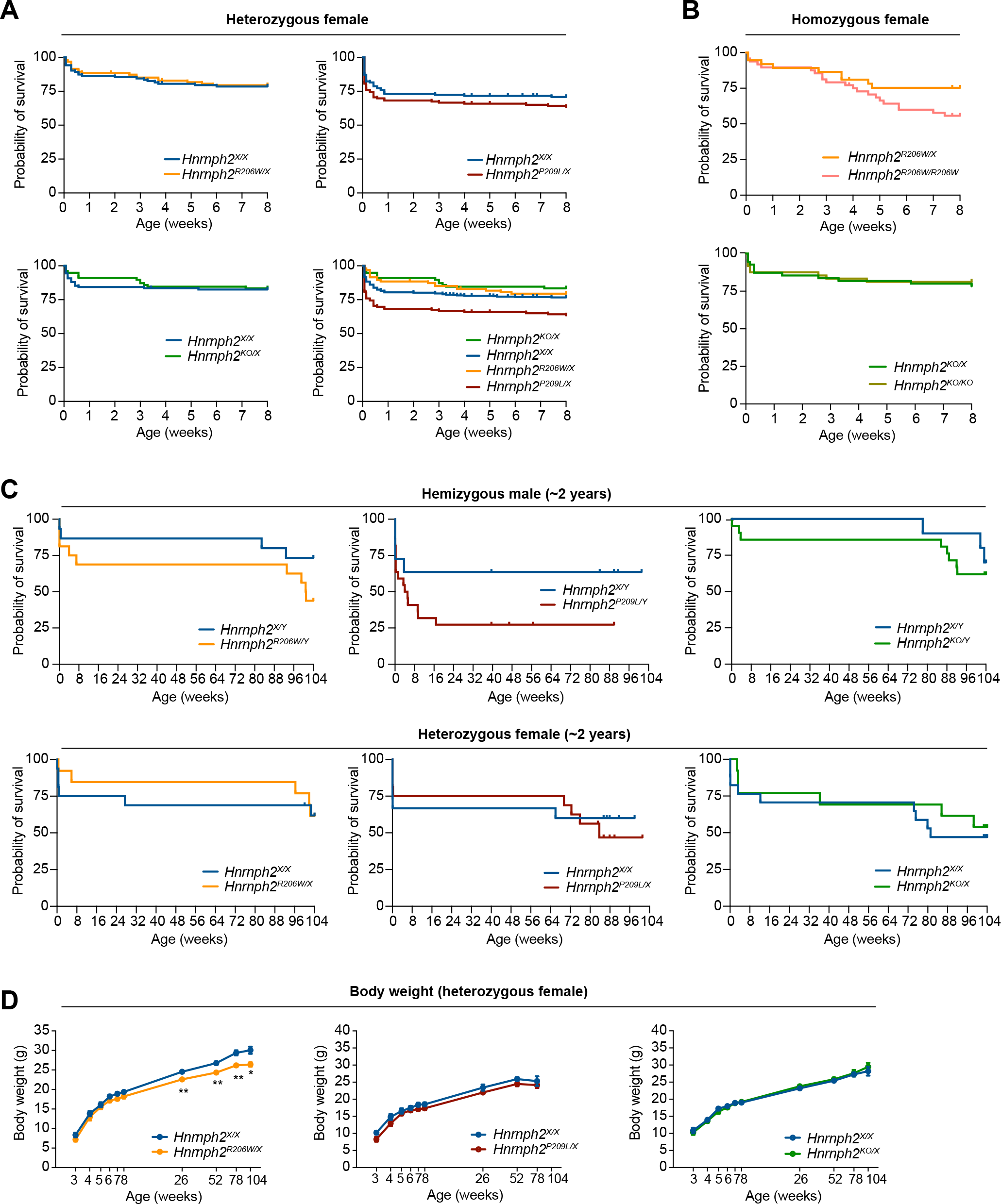
Survival and body weight of *Hnrnph2* mice. (**A-B**) Kaplan-Meier survival curves of female heterozygous (**A**) and homozygous (**B**) mice up to 8 weeks of age. (**A**) *P* = 0.8296, HR = 0.9355 for *Hnrnph2^R206W/X^* (n = 95) vs. *Hnrnph2^X/X^* (n = 103); *P* = 0.1893, HR = 1.306 for *Hnrnph2^P209L/X^* (n = 129) vs. *Hnrnph2^X/X^* (n = 141); *P* = 0.8142, HR = 0.9197 for *Hnrnph2^KO/X^* (n = 78) vs. *Hnrnph2^X/X^* (n = 109) by Mantel-Cox test. Bottom right graph shows overlay of all individual graphs. (**B**) *P* = 0.1026, HR = 1.889 for *Hnrnph2^R206W/R206W^* (n = 48) vs. *Hnrnph2^R206W/X^* (n = 37); *P* = 0.7415, HR = 0.8664 for *Hnrnph2^KO/KO^* (n = 48) vs. *Hnrnph2^KO/X^* (n = 55) by Mantel-Cox test. (**C**) Kaplan-Meier survival curves of male and female mice up to 2 years of age. For males, *P* = 0.1288, HR = 2.386 for *Hnrnph2^R206W/Y^* (n = 16) vs. *Hnrnph2^X/Y^* (n = 15); *P* = 0.1154, HR = 2.272 for *Hnrnph2^P209L/Y^* (n = 22) vs. *Hnrnph2^X/Y^* (n = 11); *P* = 0.5484, HR = 1.496 for *Hnrnph2^KO/Y^*(n = 21) vs. *Hnrnph2^X/Y^* (n = 10) by Mantel-Cox test. For females, *P* = 0.8944, HR = 0.9236 for *Hnrnph2^R206W/X^* (n = 13) vs. *Hnrnph2^X/X^* (n = 16); *P* = 0.6974, HR = 1.219 for *Hnrnph2^P209L/X^* (n = 16) vs. *Hnrnph2^X/X^* (n = 15); *P* = 0.6151, HR = 0.7691 for *Hnrnph2^KO/X^* (n = 13) vs. *Hnrnph2^X/X^* (n = 17) by Mantel-Cox test. (**D**) Mean body weight of female mice over time; error bars represent mean ± SEM. *Hnrnph2^R206W/X^* vs. *Hnrnph2^X/X^* \*\**P* < 0.01 at 26, 52, and 78 weeks, \**P* = 0.045 at 104 weeks by mixed-effects model (REML) with Sidak’s multiple comparisons test. Group sizes were as follows: *Hnrnph2^R206W/X^* (n = 11) vs. *Hnrnph2^X/X^* (n = 12); *Hnrnph2^P209L/X^* (n = 12) vs. *Hnrnph2^X/X^* (n = 10); *Hnrnph2^KO/X^* (n = 10) vs. *Hnrnph2^X/X^* (n = 13).

**Supplemental Figure 7.**
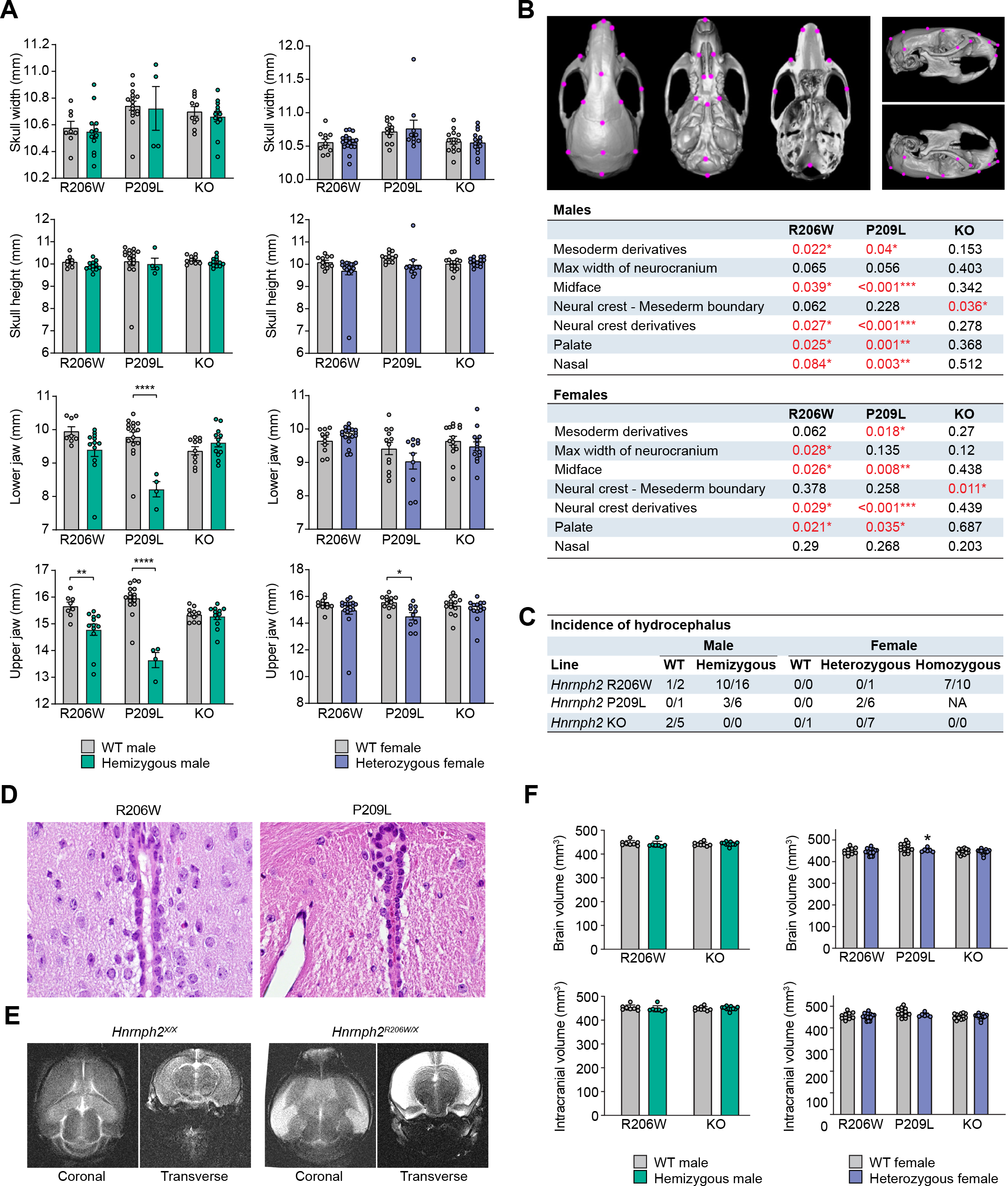
Craniofacial dysmorphology and hydrocephalus in *Hnrnph2* mice. (**A**) Key craniofacial parameters measured manually on individual µCT scans. Error bars represent mean ± SEM. Lower jaw length: \*\*\*\**P* < 0.0001 *Hnrnph2^P209L/Y^* vs. *Hnrnph2^X/Y^*; upper jaw: \*\**P* = 0.0014 *Hnrnph2^R206W/Y^* vs. *Hnrnph2^X/Y^*; \*\*\*\**P* < 0.0001 *Hnrnph2^P209L/Y^* vs. *Hnrnph2^X/Y^*; \**P* = 0.0139 *Hnrnph2^P209L/X^* vs. *Hnrnph2^X/X^* by two-way ANOVA with Sidak’s multiple comparisons test. Group sizes for µCT were as follows: *Hnrnph2^R206W/Y^* (n = 12) vs. *Hnrnph2^X/Y^* (n = 8); *Hnrnph2^P209L/Y^* (n = 4) vs. *Hnrnph2^X/Y^*(n = 16); *Hnrnph2^KO/Y^* (n = 12) vs. *Hnrnph2^X/Y^* (n = 10); *Hnrnph2^R206W/X^* (n = 17) vs. *Hnrnph2^X/X^* (n = 11); *Hnrnph2^P209L/X^* (n = 10) vs. *Hnrnph2^X/X^* (n = 12); *Hnrnph2^KO/X^* (n = 14) vs. *Hnrnph2^X/X^*(n = 14). (**B**) Figure depicting the subset of anatomically relevant landmarks used in the regional EDMA analysis and *P* values of the regional EDMA analysis. (**C**) Number of mice found dead or flagged for domed heads with pathologically confirmed hydrocephalus. (**D**) H&E staining showing patent aqueducts in hnRNPH2 P209L (hemizygous male) and hnRNPH2 R206W (heterozygous female) mice with hydrocephalus. (**E**) Representative MRI images showing hydrocephalus in a *Hnrnph2^R206W/X^* heterozygous female compared with a WT littermate. (**F**) Brain volume (grey matter + white matter) and intracranial volume (brain volume + CSF volume) measured on individual MRI scans. Error bars represent mean ± SEM. \**P* = 0.0496 *Hnrnph2^P209L/X^* vs. *Hnrnph2^X/X^* by two-way ANOVA with Sidak’s multiple comparisons test. Group sizes for brain volume analysis were as follows: *Hnrnph2^R206W/Y^* (n = 9) vs. *Hnrnph2^X/Y^* (n = 7); *Hnrnph2^KO/Y^* (n = 11) vs. *Hnrnph2^X/Y^* (n = 9); *Hnrnph2^R206W/X^* (n = 16) vs. *Hnrnph2^X/X^* (n = 11); *Hnrnph2^P209L/X^* (n = 7) vs. *Hnrnph2^X/X^* (n = 12); *Hnrnph2^KO/X^* (n = 12) vs. *Hnrnph2^X/X^* (n = 14).

**Supplemental Figure 8.**
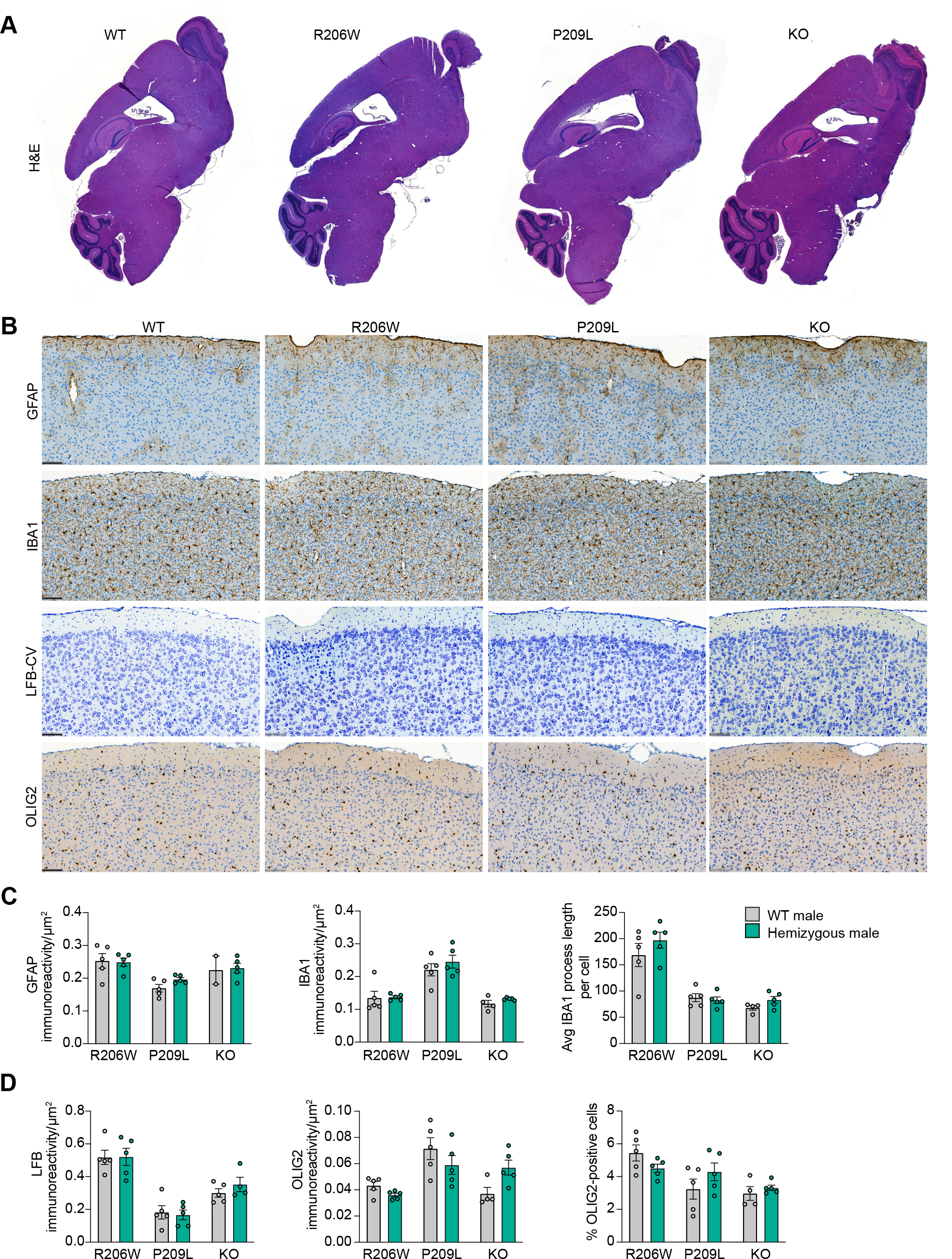
H&E staining, Luxol fast blue-cresyl violet (LFB-CV) staining, and immunohistochemistry in hnRNPH2 P209L hemizygous male mouse brains. (A) H&E staining showing no gross abnormalities in the brains of mutant males compared to WT littermates. (B) Immunohistochemistry with markers against astrocytes (GFAP), microglia (IBA1), and oligodendrocytes (OLIG2) was normal. LFB-CV staining also showed regular morphology of neurons in gray and white matters. Representative images of the primary motor and somatosensory cortex for each stain and marker are shown. Scale bar, 100 μm. (C and D) Quantification of GFAP and IBA1 (C) and LFB-CV staining and OLIG2 immunoreactivity (D) in the whole brain are shown. Error bars represent mean ± SEM. n = 5 per group.

**Supplemental Figure 9.**
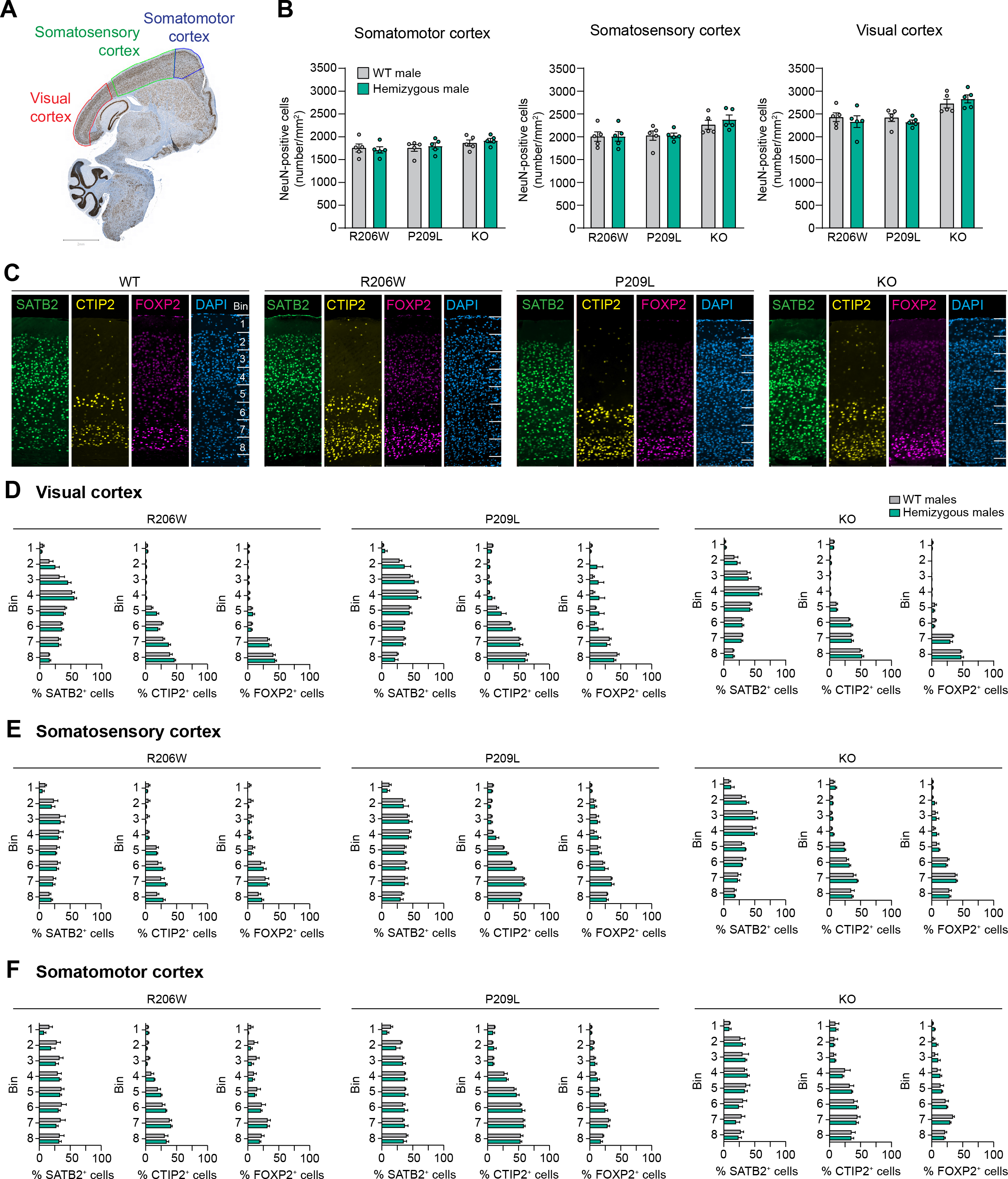
Cortical neuronal count and distribution in *Hnrnph2* mice. (**A**) Immunohistochemistry with NeuN in a WT mouse brain with manual annotation of visual, somatosensory, and somatomotor cortex. (**B**) Quantification of neurons in somatomotor, somatosensory, and visual cortex. The number of NeuN-positive cells per mm^2^ is shown. Error bars represent mean ± SEM. (**C**) Immunofluorescence with cortical layer-specific markers SATB2 (layer II-IV), CTIP2 (layer V), and FOXP2 (layer VI) performed in WT, hnRNPH2 R206W, hnRNPH2 P209L, and *Hnrnph2* KO male mice. Regions of interest were positioned over the visual, somatomotor, and somatosensory cortex and subdivided into 8 equal bins. (**D– F**) The number of SATB2-, CTIP2-, and FOXP2-positive cells were counted and expressed as a percentage of the total number of DAPI-positive cells within each bin in the visual (**D**), somatosensory (**E**), and somatomotor cortex (**F**) as defined in panel (**C**). Error bars represent mean ± SEM. n = 5 per group.

**Supplemental Figure 10.**
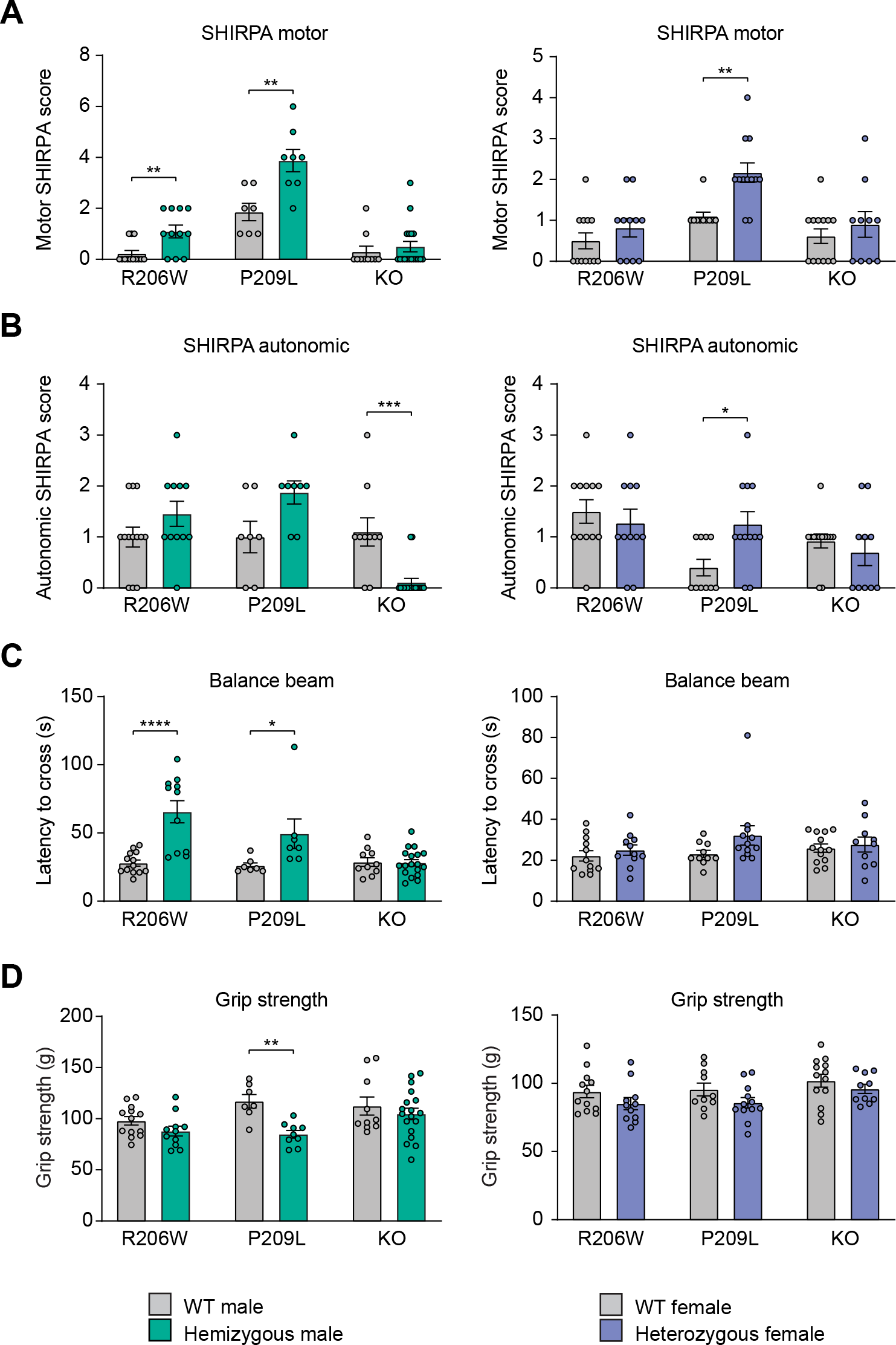
H*n*rnph2 mutant mice have impaired motor function. (**A** and **B**) Subdomain SHIRPA scores are shown. Motor function (**A**): \*\**P* = 0.008 *Hnrnph2^R206W/Y^* vs. *Hnrnph2^X/Y^*; \*\**P* = 0.008 *Hnrnph2^P209L/Y^* vs. *Hnrnph2^X/Y^*; \*\**P* = 0.001 *Hnrnph2^P209L/X^* vs. *Hnrnph2^X/X^* by two-way non-parametric ANOVA with Mann-Whitney U test for group wise comparisons. Autonomic function (**B**): \*\*\**P* = 0.0003 *Hnrnph2^KO/Y^* vs. *Hnrnph2^X/Y^*; \**P* = 0.017 *Hnrnph2^P209L/X^* vs. *Hnrnph2^X/X^* by two-way non-parametric ANOVA with Mann-Whitney U test for group wise comparisons. Error bars represent mean ± SEM in all graphs. (**C**) Latency to cross balance beam, \*\*\*\**P* < 0.0001 *Hnrnph2^R206W/Y^* vs. *Hnrnph2^X/Y^*; \**P* = 0.0264 *Hnrnph2^P209L/Y^* vs. *Hnrnph2^X/Y^* by two-way ANOVA with Sidak’s multiple comparisons test. Error bars represent mean ± SEM in all graphs. (**D**) Grip strength, \*\**P* = 0.0065 *Hnrnph2^P209L/Y^* vs. *Hnrnph2^X/Y^* by two-way ANOVA with Sidak’s multiple comparisons test. Error bars represent mean ± SEM in all graphs. Group sizes for SHIRPA and motor tests were as follows: *Hnrnph2^R206W/Y^* (n = 11) vs. *Hnrnph2^X/Y^*(n = 13); *Hnrnph2^P209L/Y^* (n = 8) vs. *Hnrnph2^X/Y^* (n = 7); *Hnrnph2^KO/Y^* (n = 18) vs. *Hnrnph2^X/Y^* (n = 10); *Hnrnph2^R206W/X^* (n = 11) vs. *Hnrnph2^X/X^* (n = 12); *Hnrnph2^P209L/X^* (n = 12) vs. *Hnrnph2^X/X^* (n = 10); *Hnrnph2^KO/X^* (n = 10) vs. *Hnrnph2^X/X^* (n = 13).

**Supplemental Figure 11.**
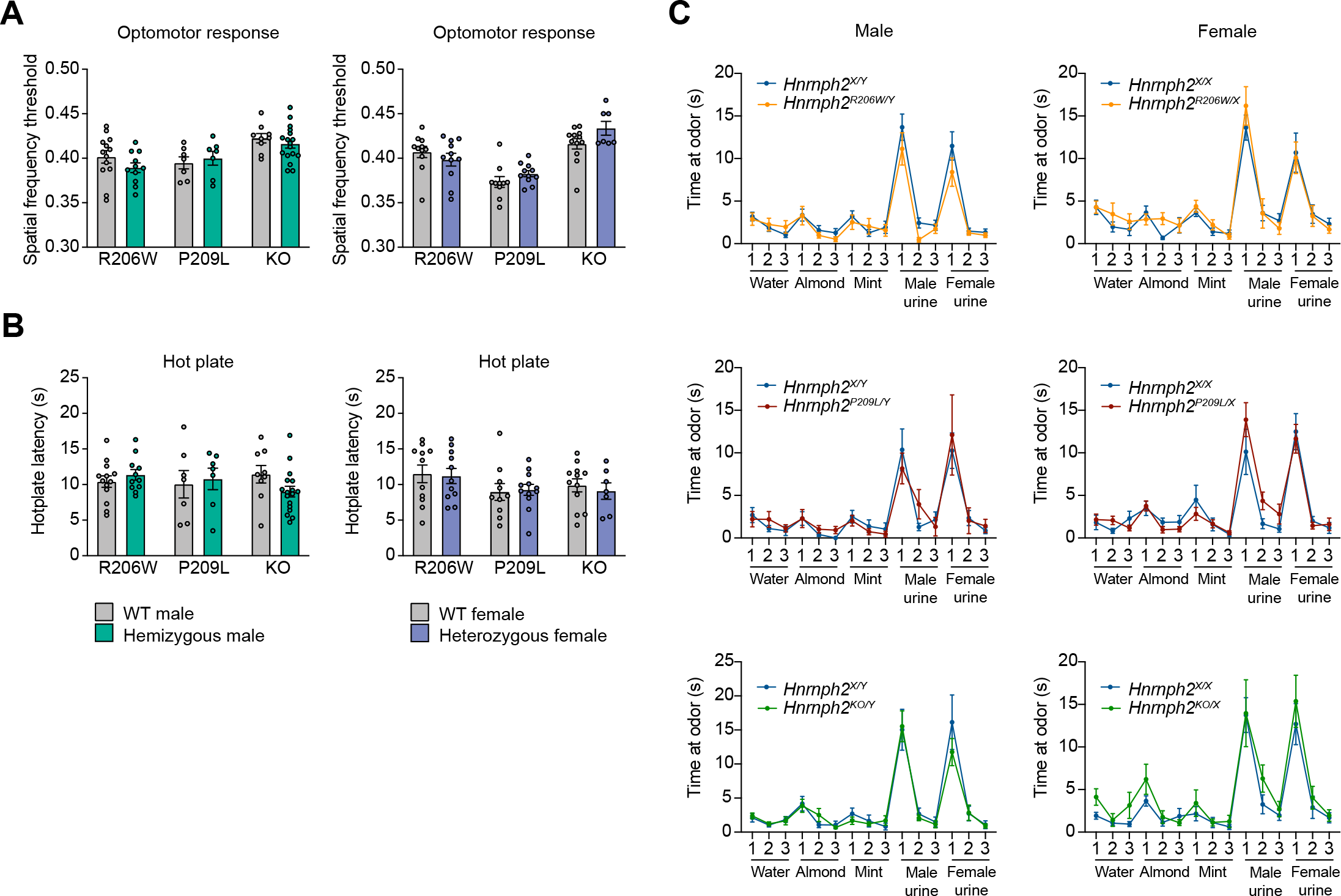
H*n*rnph2 mutant and KO mice have normal sensory function. (**A**) Optomotor response test of visual acuity. (**B**) Hot plate test of pain response. (**C**) Scent habituation test of olfaction. Error bars represent mean ± SEM in all graphs. Group sizes were as follows: *Hnrnph2^R206W/Y^* (n = 11) vs. *Hnrnph2^X/Y^* (n = 13); *Hnrnph2^P209L/Y^* (n = 7) vs. *Hnrnph2^X/Y^* (n = 7); *Hnrnph2^KO/Y^* (n = 17) vs. *Hnrnph2^X/Y^* (n = 10); *Hnrnph2^R206W/X^* (n = 11) vs. *Hnrnph2^X/X^* (n = 12); *Hnrnph2^P209L/X^* (n = 11) vs. *Hnrnph2^X/X^*(n = 10); *Hnrnph2^KO/X^* (n = 7) vs. *Hnrnph2^X/X^* (n = 12).

**Supplemental Figure 12.**
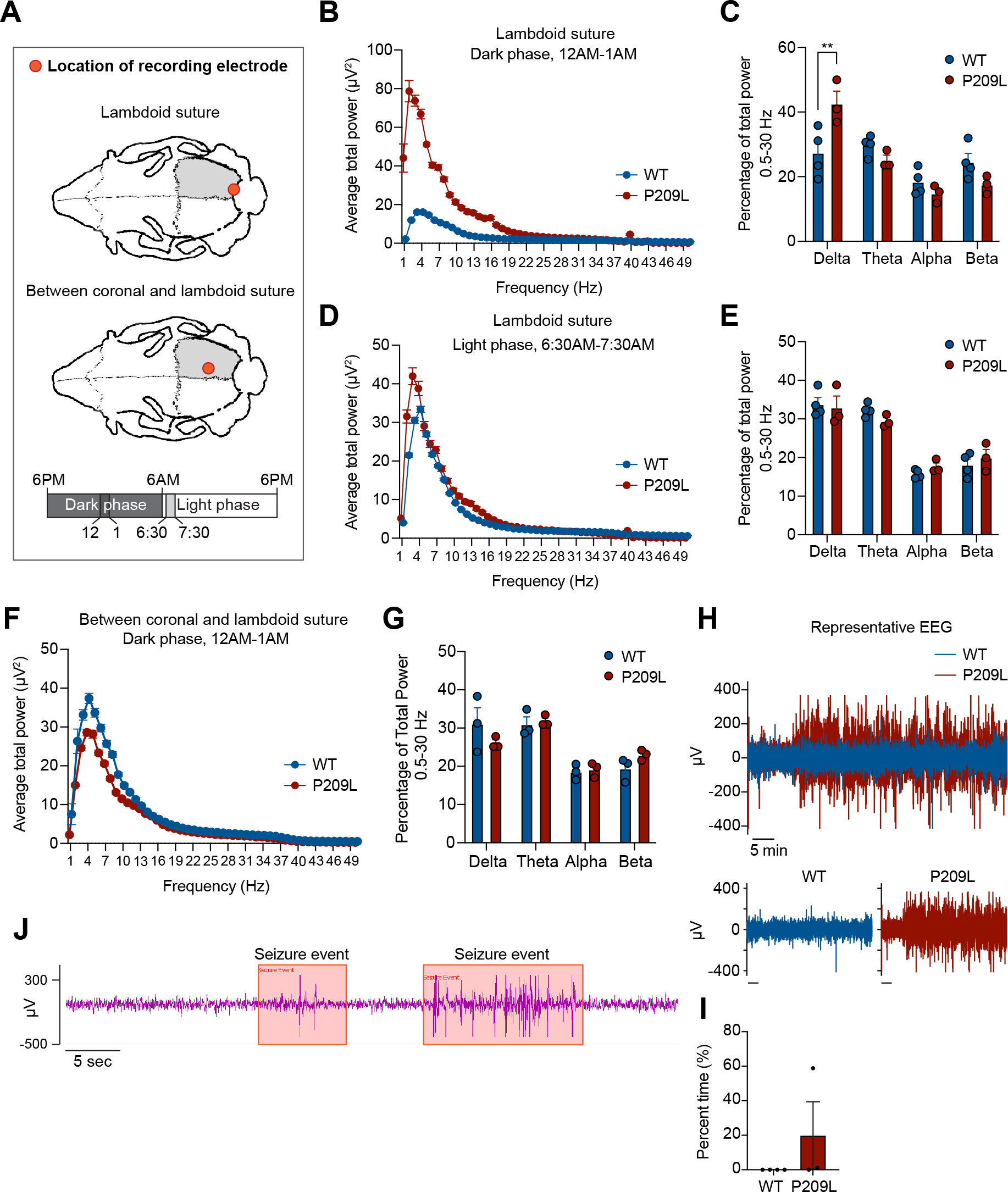
Cortical EEG dynamics are altered during the dark phase in *Hnrnph2* mutant mice. (**A**) Schematic depicting location of EEG leads. Top: lambdoid suture; bottom: between coronal and lambdoid suture. Mice were monitored and analyzed for 24 hours. Representative 1-hour windows for the dark phase (12am-1am) and light phase (6:30am- 7:30am) are presented. (**B, C**) Group analysis for a lead at the right lambdoid suture showing spectral power (**B**) and the relative power distribution across wavelength categories (**C**) from 12am to 1am. \*\**P* = 0.0021 by two-way ANOVA with Sidak’s multiple comparisons test. (**D, E**) Group analysis for a lead at the right lambdoid suture showing average spectral power (**D**) and the relative power distribution across wavelength categories (**E**) from 6:30am to 7:30am. (**F, G**) Group analysis for a lead between the right coronal and lambdoid sutures showing average spectral power (**F**) and the relative power distribution across wavelength categories (**G**) from 12am to 1am. (**H**) Representative EEG activity in the lead at the right lambdoid suture from 12am to 1 am. (**I**) Percentage of time with epileptiform activity in the lead at the right lambdoid suture from 12am to 1 am. (**J**) EEG activity of P209L hemizygous male showing seizure-like behavior. Group sizes for EEG analysis were as follows: *Hnrnph2^P209L/Y^* (n = 3) vs. *Hnrnph2^X/Y^* (n = 4).

**Supplemental Figure 13.**
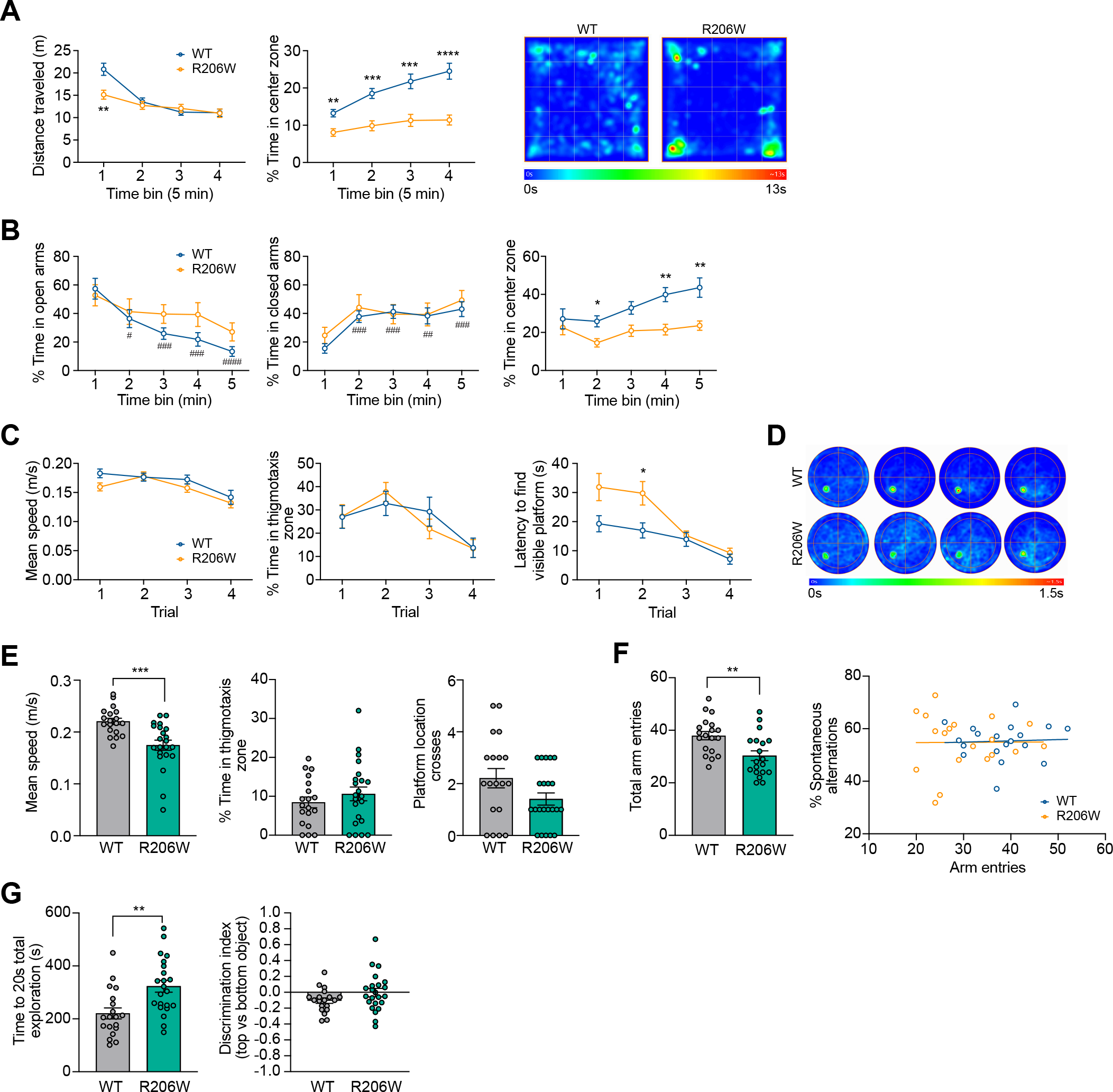
hnRNPH2 R206W males have increased anxiety, impaired spatial learning and memory, deficits in social interaction, and reduced marble burying. (**A**) Distance traveled (time bin 1 \*\**P* = 0.0069 *Hnrnph2^R206W/Y^* (n = 22) vs. *Hnrnph2^X/Y^* (n = 19)) and percentage time spent in the center zone (time bin 1 \*\**P* = 0.0031, time bin 2 \*\*\**P* = 0.0002, time bin 3 \*\*\**P* = 0.0009, time bin 4 \*\*\*\**P* < 0.0001 *Hnrnph2^R206W/Y^* vs. *Hnrnph2^X/Y^*) of the open field test across 5-minute bins by two-way ANOVA with Sidak’s multiple comparisons test. At right is an averaged heat map of the animals’ center point for the groups for the entire duration of the 20-minute test. (**B**) Percentage time spent in the open arms (time bin 2 ^#^*P* = 0.0359, time bin 3 ^###^*P* = 0.0004, time bin 4 ^###^*P* = 0.0004, time bin 5 ^####^*P* < 0.0001 vs. time bin 1 for *Hnrnph2^X/Y^* by two-way ANOVA with Dunnett’s multiple comparisons test), closed arms (time bin 2 ^###^*P* = 0.0002, time bin 3 ^###^*P* = 0.0006, time bin 4 ^##^*P* = 0.0042, time bin 5 ^###^*P* = 0.0004 vs. time bin 1 for *Hnrnph2^X/Y^*by two-way ANOVA with Dunnett’s multiple comparisons test), and center zone (time bin 2 \**P* = 0.02, time bin 4 \*\**P* = 0.0024, time bin 5 \*\**P* = 0.0087 *Hnrnph2^R206W/Y^* (n = 13) vs. *Hnrnph2^X/Y^* (n = 18) by two-way ANOVA with Sidak’s multiple comparisons test) of the elevated plus maze across 1-minute bins. (**C**) Mean speed, percentage time spent in the thigmotaxis zone, and latency to find the visible platform (trial 2 \**P* = 0.0467 *Hnrnph2^R206W/Y^* (n = 22) vs. *Hnrnph2^X/Y^*(n = 19) by two-way ANOVA with Sidak’s multiple comparisons test) in the cued trials of the Morris water maze. (**D**) Averaged heat map of the animals’ center point for the groups for training day 1-4 of the Morris water maze. (**E**) Mean speed (\*\*\**P* = 0.0004 *Hnrnph2^R206W/Y^* (n = 22) vs. *Hnrnph2^X/Y^*(n = 19) by unpaired t test), percentage time spent in the thigmotaxis zone, and number of platform location crossings during the probe trial of the Morris water maze. (**F**) Number of total arm entries (\*\**P* = 0.0034 *Hnrnph2^R206W/Y^* (n = 20) vs. *Hnrnph2^X/Y^* (n = 19) by unpaired t test) and lack of correlation between percentage spontaneous alternations and number of total arm entries by Pearson’s correlation analysis. (**G**) Time to reach 20 seconds of total object exploration (\*\**P* = 0.0019 *Hnrnph2^R206W/Y^* (n = 22) vs. *Hnrnph2^X/Y^* (n = 19) by unpaired t test) and object position discrimination index in the familiarization stage of the novel object recognition test.

**Supplemental Figure 14.**
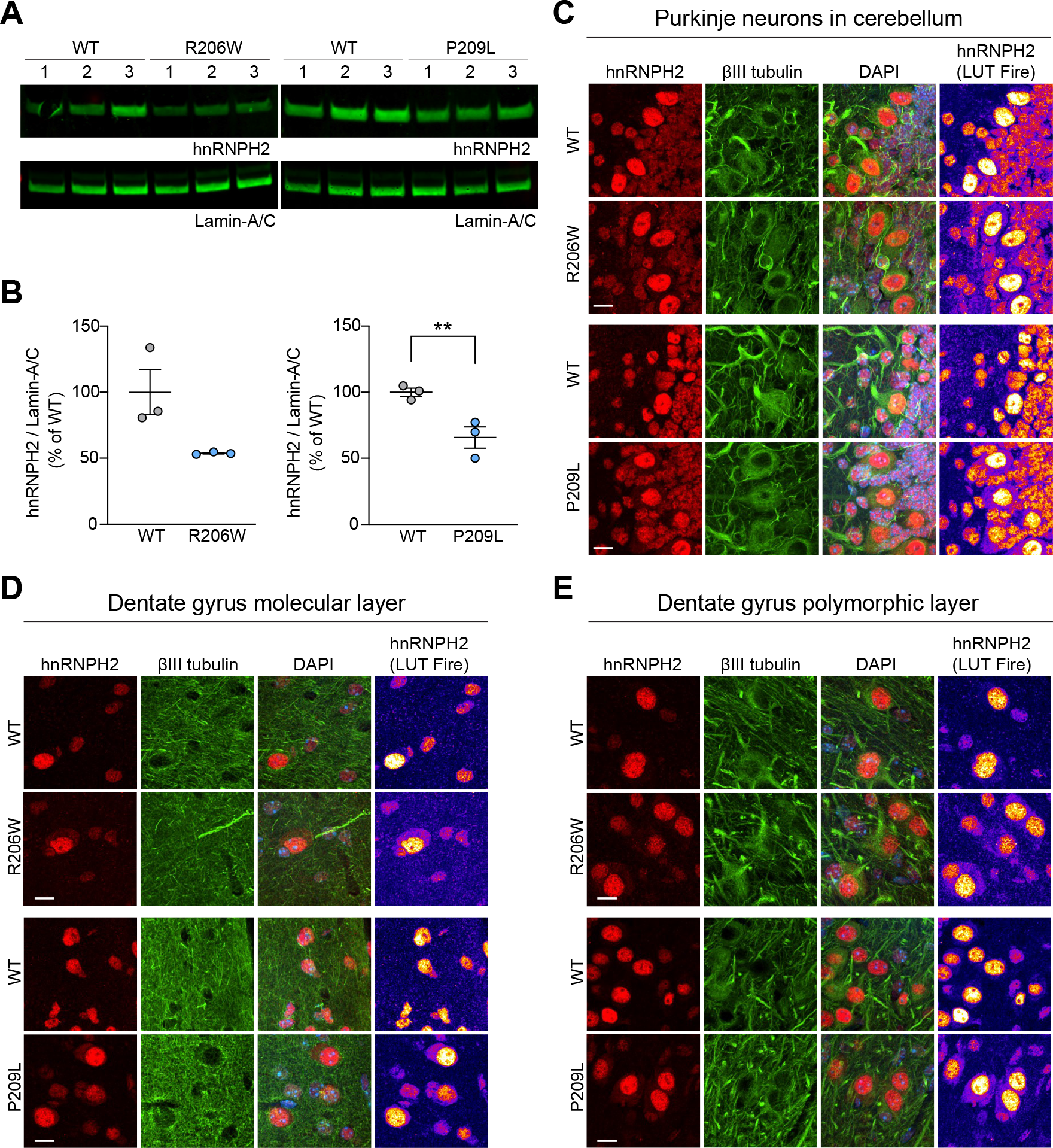
Pathogenic variants alter the nucleocytoplasmic ratio of hnRNPH2 in hnRNPH2 P209L and R206W mice. (**A**) Immunoblot of hnRNPH2 in cortical nuclear fractions. Lamin A/C was used as a loading control; labels 1-3 indicate three biological replicates. (**B**) Quantification of hnRNPH2 normalized to lamin A/C; error bars represent mean ± SEM. \*\**P* = 0.0063 *Hnrnph2^P209L/Y^* vs. *Hnrnph2^X/Y^* by two-way ANOVA with Sidak’s multiple comparisons test. n = 3 per group. (**C-E**) Immunofluorescent staining of hnRNPH2 in mouse brain sections. Purkinje neurons in cerebellum (**C**), molecular layer (**D**) and polymorphic layer (**E**) of dentate gyrus are shown. βIII tubulin and DAPI were used as neuronal cytoplasmic and nuclear markers, respectively. Look-up table (LUT) fire was used to increase the visibility of the hnRNPH2 cytoplasmic signal. Scale bars, 10 μm.

**Supplemental Figure 15.**
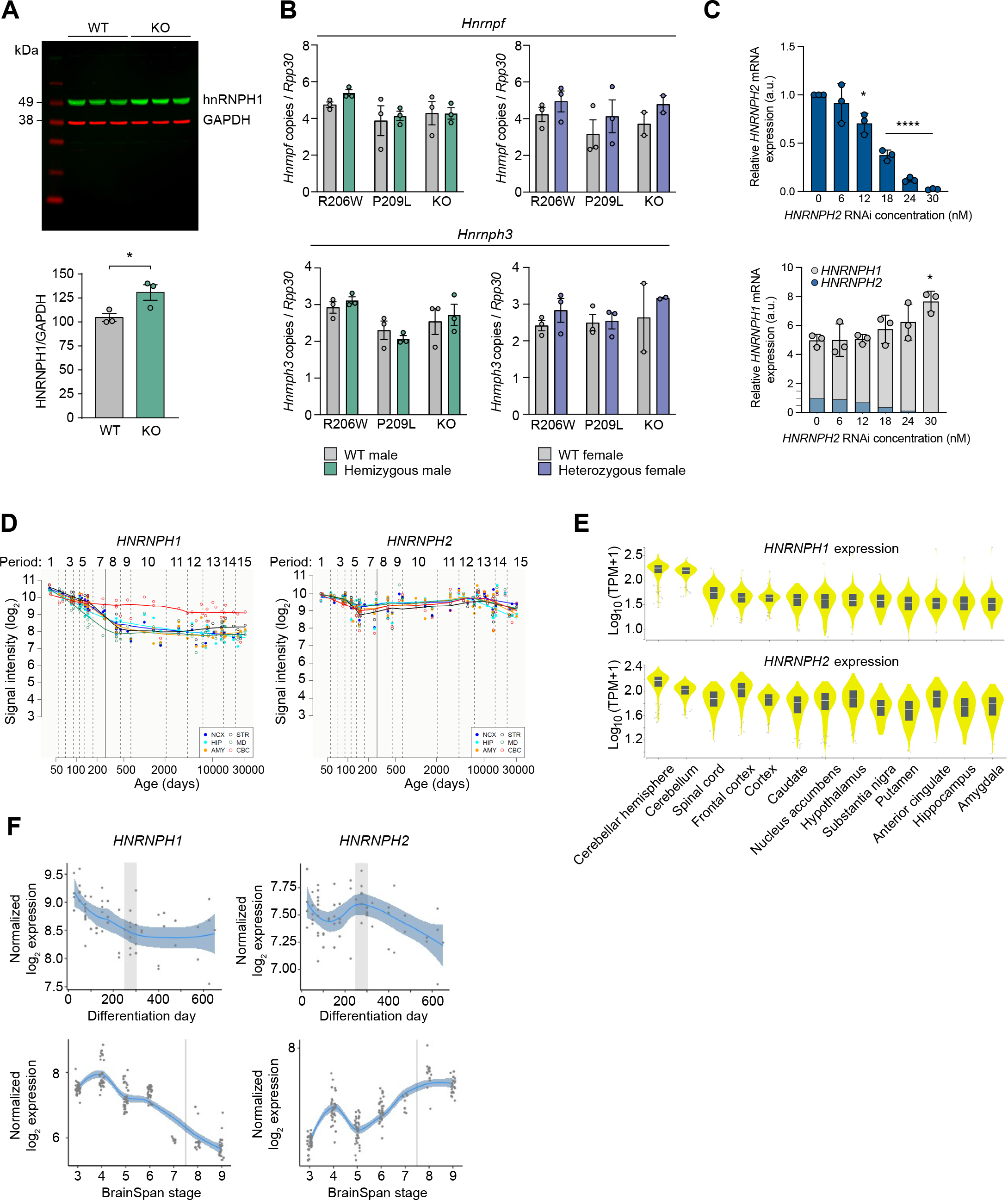
Spatiotemporal expression of *HNRNPH1* and *HNRNPH2* in human brain and cortical organoids. (**A**) Immunoblot and quantification of hnRNPH1 expression in whole brain RIPA-soluble fractions. GAPDH was used as a loading control; \**P* = 0.0454 WT vs. *Hnrnph2* KO by t-test. Error bars represent mean ± SEM. n = 3 per group. (**B**) Number of *Hnrnpf* and *Hnrnph3* copies normalized to *Rpp30* in the cortex of *Hnrnph2^R206W^*, *Hnrnph2^P209L^*, and *Hnrnph2^KO^*mice by ddRT-PCR. Error bars represent mean ± SEM. n = 3 per group, except for *Hnrnph2^KO/X^* and *Hnrnph2^X/X^* which are n=2 per group. (**C**) Relative expression levels of *HNRNPH1* and *HNRNPH2* in HEK293T cells transfected with siRNA targeting *HNRNPH2* by qRT-PCR. To show the relative expression of *HNRNPH1* and *HNRNPH2*, the *HNRNPH2* graph is overlaid on the *HNRNPH1* graph with an adjusted scale. \**P* = 0.0107 (*HNRNPH2*), 0.0106 (*HNRNPH1*), and \*\*\*\**P* < 0.0001 by one-way ANOVA with Dunnett’s multiple comparisons test. Error bars represent mean ± SD. (**D**) Trajectory plots showing the expression of *HNRNPH1* and *HNRNPH2* in 6 major brain regions across 15 developmental time points by Affymetrix GeneChip Human Exon 1.0 ST Arrays. Period 1-7, fetal development; solid line, birth; period 8-9, infancy; period 10-11, childhood; period 12, adolescence; period 13-15, adulthood. Neocortex (NCX), hippocampus (HIP), amygdala region (AMY), striatum (STR), mediodorsal nucleus of the thalamus (MD), and cerebellar cortex (CBC) are shown. Reprinted from the Human Brain Transcriptome dataset (43, 71, 72). (**E**) Violin plots showing the expression of *HNRNPH1* and *HNRNPH2* in 13 brain regions by RNA-seq. Data used for the analyses described here were obtained from the Genotype-Tissue Expression (GTEx) Portal, dbGaP accession number: phs000424.v8.p2. TPM, transcripts per million. (**F**) Trajectory plots showing the expression of *HNRNPH1* and *HNRNPH2* in human cortical organoids across differentiation day and mapped BrainSpan stages by RNA-seq. Stage 3-7, fetal development; stage 8, birth to 6 months; stage 9, 6 months to 19 months. Transition from prenatal to postnatal stages is indicated with a vertical grey area/line. Reprinted from the Gene Expression in Cortical Organoids dataset (45).

**Supplemental Figure 16.**
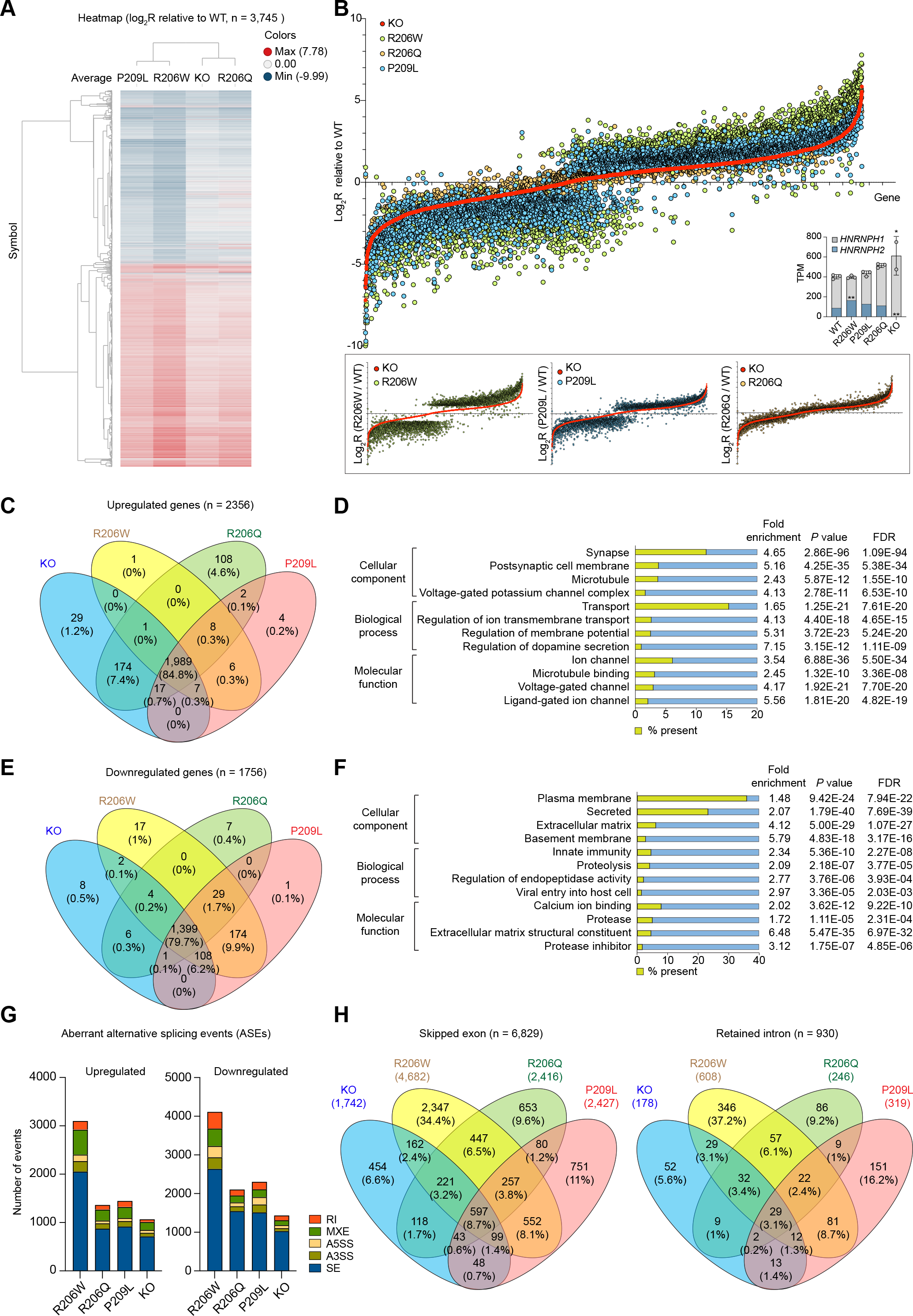
RNA sequencing of human iPSC-derived neurons harboring a pathogenic mutation or deletion of *HNRNPH2*. (**A**) Heatmap of differentially expressed genes from 3-week-old neurons comparing WT (n = 3), hnRNPH2 R206W (n = 3), hnRNPH2 R206Q (n = 3), and *HNRNPH2* KO (n = 2). Colors of bars represent log2 ratio normalized to WT and scaled expression levels. Red, white, and blue correspond to high (max = 7.78), 0, and low (min = -9.99) expression, respectively. (**B**) Log2 ratio (relative to WT) for each genotype is plotted for individual genes (X-axis). Inset bar graphs show *HNRNPH1* and *HNRNPH2* transcript levels in each genotype. \*\**P* = 0.0024 (R206W) and \*\**P* = 0.0026 (KO) by one-way ANOVA with Dunnett’s multiple comparisons test. Error bars represent mean ± SD. (**C**) Venn diagram showing the overlap of genes upregulated in each genotype. (**D**) Gene ontology (GO) analysis of commonly upregulated genes in mutant and KO neurons (1,989 genes). (**E**) Venn diagram showing the overlap of genes downregulated in each genotype. (**F**) GO analysis of commonly downregulated genes in mutant and KO neurons (1,399 genes). (**G**) Graphs summarizing aberrant alternative splicing events. Numbers of upregulated and downregulated events are shown separately. RI: retained intron, MXE: mutually exclusive exons, A5SS: alternative 5′ splice sites, A3SS: alternative 3′ splice sites, SE: skipped exons. (**H**) Venn diagrams showing the overlap of skipped exon and retained intron events in each genotype.

**Supplemental Figure 17.**
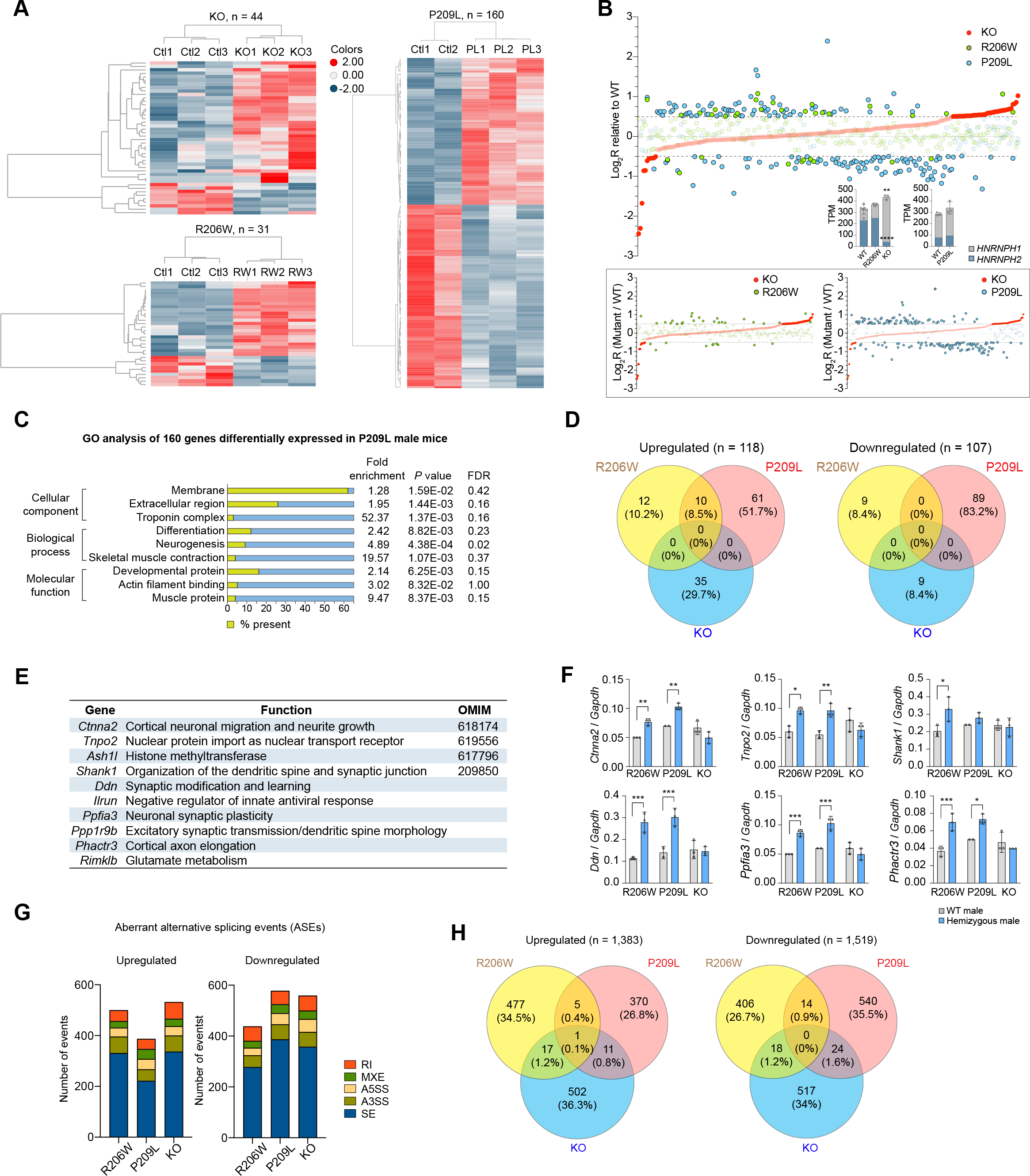
RNA sequencing of cortices from mice harboring a pathogenic mutation or deletion of *Hnrnph2*. (**A**) Heatmap of differentially expressed genes in cortices from 8-week-old KO (n = 3) and R206W male mice (n = 3) and 3-week-old P209L male mice (n = 3). Littermate WT male mice (n = 3, except for WT males from the P209L line which was n = 2)) were used as controls. Colors of bars represent log2 ratio normalized to WT and scaled expression levels. Red, white, and blue correspond to high (max = 2.00), 0, and low (min = - 2.00) expressions, respectively. (**B**) Log2 ratio (mutant / WT) for each mutant is plotted for individual genes (X-axis). Inset bar graphs show *HNRNPH1* and *HNRNPH2* transcript levels in each genotype. \*\*\*\**P* < 0.0001 (*Hnrnph2* in KO) and \*\**P* = 0.0029 (*Hnrnph1* in KO) by one-way ANOVA with Dunnett’s multiple comparisons test. Error bars represent mean ± SD. (**C**) Gene ontology (GO) analysis of differentially regulated genes in P209L male mice compared to littermate controls (536 genes). (**D**) Venn diagrams showing the overlap of genes upregulated and downregulated in each genotype. (**E**) Table listing 10 commonly upregulated genes in R206W and P209L mice and their known neuron-related functions. (**F**) ddRT-PCR analyses validating upregulation of subset of genes listed in (**E**). For *Ctnna2*, \*\**P* = 0.0031 (R206W), \*\**P* = 0.0013 (P209L); for *Tnpo2*, \**P* = 0.0111 (R206W), \*\**P* = 0.0100 (P209L); for *Shank1*, \**P* = 0.0128 (R206W); for *Ddn, \**\*\**P* = 0.0003 (R206W), \*\*\**P* = 0.0008 (P209L); for *Ppfia3*, \*\*\**P* = 0.0006 (R206W), \*\*\**P* = 0.0004 (P209L); for *Phactr3*, \*\*\**P* = 0.0005 (R206W), \**P* = 0.0159 (P209L) by two-way ANOVA with Sidak’s multiple comparisons test. Error bars represent mean ± SD. n = 3 per group, except for WT males from the P209L line which was n = 2. (**G**) Graphs summarizing aberrant alternative splicing events. Numbers of upregulated and downregulated events are shown separately. RI: retained intron, MXE: mutually exclusive exons, A5SS: alternative 5′ splice sites, A3SS: alternative 3′ splice sites, SE: skipped exons. (**H**) Venn diagrams showing the overlap of upregulated and downregulated ASEs in each genotype.

**Supplemental Table 1.** Regional brain volumes normalized to total brain tissue volume in mice harboring a pathogenic mutation or deletion of *Hnrnph2*. Group sizes for brain volume analysis were as follows: *Hnrnph2^R206W/Y^* (n = 9) vs. *Hnrnph2^X/Y^* (n = 7); *Hnrnph2^KO/Y^* (n = 11) vs. *Hnrnph2^X/Y^* (n = 9); *Hnrnph2^R206W/X^* (n = 16) vs. *Hnrnph2^X/X^* (n = 11); *Hnrnph2^P209L/X^* (n = 7) vs. *Hnrnph2^X/X^* (n = 12); *Hnrnph2^KO/X^* (n = 12) vs. *Hnrnph2^X/X^* (n = 14).

**Supplemental Table 2.** RNA composition of human iPSC-derived neurons harboring a pathogenic mutation or deletion of HNRNPH2.

**Supplemental Table 3.** RNA composition of cortices from male mice harboring a pathogenic mutation or deletion of *Hnrnph2*.

## References

1. Bain JM, Cho MT, Telegrafi A, Wilson A, Brooks S, Botti C, et al. Variants in HNRNPH2 on the X Chromosome Are Associated with a Neurodevelopmental Disorder in Females. Am J Hum Genet. 2016;99(3):728–34.

2. Bain JM, Thornburg O, Pan C, Rome-Martin D, Boyle L, Fan X, et al. Detailed Clinical and Psychological Phenotype of the X-linked HNRNPH2-Related Neurodevelopmental Disorder. Neurol Genet. 2021;7(1):e551.

3. White-Brown AM, Lemire G, Ito YA, Thornburg O, Bain JM, and Dyment DA. A disease-causing variant in HNRNPH2 inherited from an unaffected mother with skewed X-inactivation. Am J Med Genet A. 2022;188(2):668–71.

4. Kreienkamp HJ, Wagner M, Weigand H, McConkie-Rossell A, McDonald M, Keren B, et al. Variant-specific effects define the phenotypic spectrum of HNRNPH2-associated neurodevelopmental disorders in males. Hum Genet. 2022;141(2):257–72.

5. Somashekar PH, Narayanan DL, Jagadeesh S, Suresh B, Vaishnavi RD, Bielas S, et al. Bain type of X-linked syndromic mental retardation in a male with a pathogenic variant in HNRNPH2. Am J Med Genet A. 2020;182(1):183–8.

6. Harmsen S, Buchert R, Mayatepek E, Haack TB, and Distelmaier F. Bain type of X-linked syndromic mental retardation in boys. Clin Genet. 2019;95(6):734–5.

7. Jepsen WM, Ramsey K, Szelinger S, Llaci L, Balak C, Belnap N, et al. Two additional males with X-linked, syndromic mental retardation carry de novo mutations in HNRNPH2. Clin Genet. 2019;96(2):183–5.

8. Salazar R, Beenders S, LaMarca NM, Thornburg O, Rubin-Thompson L, Snow A, et al. Cross-sectional, quantitative analysis of motor function in females with HNRNPH2- related disorder. Res Dev Disabil. 2021;119:104110.

9. Pilch J, Koppolu AA, Walczak A, Murcia Pienkowski VA, Biernacka A, Skiba P, et al. Evidence for HNRNPH1 being another gene for Bain type syndromic mental retardation. Clin Genet. 2018;94(3-4):381–5.

10. Reichert SC, Li R, S AT, van Jaarsveld RH, Massink MPG, van den Boogaard MH, et al. HNRNPH1-related syndromic intellectual disability: Seven additional cases suggestive of a distinct syndromic neurodevelopmental syndrome. Clin Genet. 2020;98(1):91–8.

11. Geuens T, Bouhy D, and Timmerman V. The hnRNP family: insights into their role in health and disease. Hum Genet. 2016;135(8):851–67.

12. Van Dusen CM, Yee L, McNally LM, and McNally MT. A glycine-rich domain of hnRNP H/F promotes nucleocytoplasmic shuttling and nuclear import through an interaction with transportin 1. Mol Cell Biol. 2010;30(10):2552–62.

13. Lee BJ, Cansizoglu AE, Suel KE, Louis TH, Zhang Z, and Chook YM. Rules for nuclear localization sequence recognition by karyopherin beta 2. Cell. 2006;126(3):543–58.

14. Liu Q, Shu S, Wang RR, Liu F, Cui B, Guo XN, et al. Whole-exome sequencing identifies a missense mutation in hnRNPA1 in a family with flail arm ALS. Neurology. 2016;87(17):1763–9.

15. Naruse H, Ishiura H, Mitsui J, Date H, Takahashi Y, Matsukawa T, et al. Molecular epidemiological study of familial amyotrophic lateral sclerosis in Japanese population by whole-exome sequencing and identification of novel HNRNPA1 mutation. Neurobiol Aging. 2018;61:255 e9–e16.

16. Dormann D, Rodde R, Edbauer D, Bentmann E, Fischer I, Hruscha A, et al. ALS-associated fused in sarcoma (FUS) mutations disrupt Transportin-mediated nuclear import. EMBO J. 2010;29(16):2841–57.

17. Siomi MC, Eder PS, Kataoka N, Wan L, Liu Q, and Dreyfuss G. Transportin-mediated nuclear import of heterogeneous nuclear RNP proteins. J Cell Biol. 1997;138(6):1181–92.

18. Cansizoglu AE, Lee BJ, Zhang ZC, Fontoura BM, and Chook YM. Structure-based design of a pathway-specific nuclear import inhibitor. Nat Struct Mol Biol. 2007;14(5):452–4.

19. Richtsmeier JT, Baxter LL, and Reeves RH. Parallels of craniofacial maldevelopment in Down syndrome and Ts65Dn mice. Dev Dyn. 2000;217(2):137–45.

20. Maga AM, Tustison NJ, and Avants BB. A population level atlas of Mus musculus craniofacial skeleton and automated image-based shape analysis. J Anat. 2017;231(3):433–43.

21. Lele S, and Richtsmeier JT. Euclidean distance matrix analysis: a coordinate-free approach for comparing biological shapes using landmark data. Am J Phys Anthropol. 1991;86(3):415–27.

22. Menegaz RA, Ladd SH, and Organ JM. Craniofacial allometry in the OIM(-/-) mouse model of osteogenesis imperfecta. FASEB J. 2020;34(8):10850–9.

23. Hill CA, Sussan TE, Reeves RH, and Richtsmeier JT. Complex contributions of Ets2 to craniofacial and thymus phenotypes of trisomic “Down syndrome” mice. Am J Med Genet A. 2009;149A(10):2158–65.

24. Vogel P, Read RW, Hansen GM, Payne BJ, Small D, Sands AT, et al. Congenital hydrocephalus in genetically engineered mice. Vet Pathol. 2012;49(1):166–81.

25. Dorr AE, Lerch JP, Spring S, Kabani N, and Henkelman RM. High resolution three-dimensional brain atlas using an average magnetic resonance image of 40 adult C57Bl/6J mice. Neuroimage. 2008;42(1):60–9.

26. Madhok S, and Bain J. In: Adam MP, Everman DB, Mirzaa GM, Pagon RA, Wallace SE, Bean LJH, et al. eds. GeneReviews((R)). Seattle (WA); 1993.

27. Gates H, Mallon AM, Brown SD, and Consortium E. High-throughput mouse phenotyping. Methods. 2011;53(4):394–404.

28. Rogers DC, Fisher EM, Brown SD, Peters J, Hunter AJ, and Martin JE. Behavioral and functional analysis of mouse phenotype: SHIRPA, a proposed protocol for comprehensive phenotype assessment. Mamm Genome. 1997;8(10):711–3.

29. Gonzalez D, Tomasek M, Hays S, Sridhar V, Ammanuel S, Chang CW, et al. Audiogenic Seizures in the Fmr1 Knock-Out Mouse Are Induced by Fmr1 Deletion in Subcortical, VGlut2-Expressing Excitatory Neurons and Require Deletion in the Inferior Colliculus. J Neurosci. 2019;39(49):9852–63.

30. Mandel-Brehm C, Salogiannis J, Dhamne SC, Rotenberg A, and Greenberg ME. Seizure-like activity in a juvenile Angelman syndrome mouse model is attenuated by reducing Arc expression. Proc Natl Acad Sci U S A. 2015;112(16):5129–34.

31. Judson MC, Wallace ML, Sidorov MS, Burette AC, Gu B, van Woerden GM, et al. GABAergic Neuron-Specific Loss of Ube3a Causes Angelman Syndrome-Like EEG Abnormalities and Enhances Seizure Susceptibility. Neuron. 2016;90(1):56–69.

32. Rodgers RJ, and Johnson NJ. Factor analysis of spatiotemporal and ethological measures in the murine elevated plus-maze test of anxiety. Pharmacol Biochem Behav. 1995;52(2):297–303.

33. Rodgers RJ, Johnson NJ, Cole JC, Dewar CV, Kidd GR, and Kimpson PH. Plus-maze retest profile in mice: importance of initial stages of trail 1 and response to post-trail cholinergic receptor blockade. Pharmacol Biochem Behav. 1996;54(1):41–50.

34. Carobrez AP, and Bertoglio LJ. Ethological and temporal analyses of anxiety-like behavior: the elevated plus-maze model 20 years on. Neurosci Biobehav Rev. 2005;29(8):1193–205.

35. Vorhees CV, and Williams MT. Morris water maze: procedures for assessing spatial and related forms of learning and memory. Nat Protoc. 2006;1(2):848–58.

36. Higaki A, Mogi M, Iwanami J, Min LJ, Bai HY, Shan BS, et al. Recognition of early stage thigmotaxis in Morris water maze test with convolutional neural network. PLoS One. 2018;13(5):e0197003.

37. Huang Y, Zhou W, and Zhang Y. Bright lighting conditions during testing increase thigmotaxis and impair water maze performance in BALB/c mice. Behav Brain Res. 2012;226(1):26–31.

38. Dalm S, Grootendorst J, de Kloet ER, and Oitzl MS. Quantification of swim patterns in the Morris water maze. Behav Res Methods Instrum Comput. 2000;32(1):134–9.

39. Hisey E, and Soderling SH. A prefrontal to lateral entorhinal pathway disrupts memory. bioRxiv. 2022:2022.02.21.480888.

40. Thomas A, Burant A, Bui N, Graham D, Yuva-Paylor LA, and Paylor R. Marble burying reflects a repetitive and perseverative behavior more than novelty-induced anxiety. Psychopharmacology (Berl). 2009;204(2):361–73.

41. Grammatikakis I, Zhang P, Panda AC, Kim J, Maudsley S, Abdelmohsen K, et al. Alternative Splicing of Neuronal Differentiation Factor TRF2 Regulated by HNRNPH1/H2. Cell Rep. 2016;15(5):926–34.

42. Lein ES, Hawrylycz MJ, Ao N, Ayres M, Bensinger A, Bernard A, et al. Genome-wide atlas of gene expression in the adult mouse brain. Nature. 2007;445(7124):168–76.

43. Kang HJ, Kawasawa YI, Cheng F, Zhu Y, Xu X, Li M, et al. Spatio-temporal transcriptome of the human brain. Nature. 2011;478(7370):483–9.

44. Consortium GT, Laboratory DA, Coordinating Center -Analysis Working G, Statistical Methods groups-Analysis Working G, Enhancing Gg, Fund NIHC, et al. Genetic effects on gene expression across human tissues. Nature. 2017;550(7675):204–13.

45. Gordon A, Yoon SJ, Tran SS, Makinson CD, Park JY, Andersen J, et al. Long-term maturation of human cortical organoids matches key early postnatal transitions. Nat Neurosci. 2021;24(3):331–42.

46. Zhang Y, Chen K, Sloan SA, Bennett ML, Scholze AR, O’Keeffe S, et al. An RNA-sequencing transcriptome and splicing database of glia, neurons, and vascular cells of the cerebral cortex. J Neurosci. 2014;34(36):11929–47.

47. Zhang Y, Sloan SA, Clarke LE, Caneda C, Plaza CA, Blumenthal PD, et al. Purification and Characterization of Progenitor and Mature Human Astrocytes Reveals Transcriptional and Functional Differences with Mouse. Neuron. 2016;89(1):37–53.

48. Saunders A, Macosko EZ, Wysoker A, Goldman M, Krienen FM, de Rivera H, et al. Molecular Diversity and Specializations among the Cells of the Adult Mouse Brain. Cell. 2018;174(4):1015–30 e16.

49. Gillentine MA, Wang T, Hoekzema K, Rosenfeld J, Liu P, Guo H, et al. Rare deleterious mutations of HNRNP genes result in shared neurodevelopmental disorders. Genome Med. 2021;13(1):63.

50. Hodgkins A, Farne A, Perera S, Grego T, Parry-Smith DJ, Skarnes WC, et al. WGE: a CRISPR database for genome engineering. Bioinformatics. 2015;31(18):3078–80.

51. Glynn D, Bortnick RA, and Morton AJ. Complexin II is essential for normal neurological function in mice. Hum Mol Genet. 2003;12(19):2431–48.

52. Luong TN, Carlisle HJ, Southwell A, and Patterson PH. Assessment of motor balance and coordination in mice using the balance beam. J Vis Exp. 2011(49).

53. Brooks SP, Trueman RC, and Dunnett SB. Assessment of Motor Coordination and Balance in Mice Using the Rotarod, Elevated Bridge, and Footprint Tests. Curr Protoc Mouse Biol. 2012;2(1):37–53.

54. Musumeci SA, Calabrese G, Bonaccorso CM, D’Antoni S, Brouwer JR, Bakker CE, et al. Audiogenic seizure susceptibility is reduced in fragile X knockout mice after introduction of FMR1 transgenes. Exp Neurol. 2007;203(1):233–40.

55. McIlwain KL, Merriweather MY, Yuva-Paylor LA, and Paylor R. The use of behavioral test batteries: effects of training history. Physiol Behav. 2001;73(5):705–17.

56. Paylor R, Spencer CM, Yuva-Paylor LA, and Pieke-Dahl S. The use of behavioral test batteries, II: effect of test interval. Physiol Behav. 2006;87(1):95–102.

57. Walf AA, and Frye CA. The use of the elevated plus maze as an assay of anxiety-related behavior in rodents. Nat Protoc. 2007;2(2):322–8.

58. Seibenhener ML, and Wooten MC. Use of the Open Field Maze to measure locomotor and anxiety-like behavior in mice. J Vis Exp. 2015(96):e52434.

59. Takahashi A, Kato K, Makino J, Shiroishi T, and Koide T. Multivariate analysis of temporal descriptions of open-field behavior in wild-derived mouse strains. Behav Genet. 2006;36(5):763–74.

60. Leger M, Quiedeville A, Bouet V, Haelewyn B, Boulouard M, Schumann-Bard P, et al. Object recognition test in mice. Nat Protoc. 2013;8(12):2531–7.

61. Lueptow LM. Novel Object Recognition Test for the Investigation of Learning and Memory in Mice. J Vis Exp. 2017(126).

62. Miedel CJ, Patton JM, Miedel AN, Miedel ES, and Levenson JM. Assessment of Spontaneous Alternation, Novel Object Recognition and Limb Clasping in Transgenic Mouse Models of Amyloid-beta and Tau Neuropathology. J Vis Exp. 2017(123).

63. Rein B, Ma K, and Yan Z. A standardized social preference protocol for measuring social deficits in mouse models of autism. Nat Protoc. 2020;15(10):3464–77.

64. Angoa-Perez M, Kane MJ, Briggs DI, Francescutti DM, and Kuhn DM. Marble burying and nestlet shredding as tests of repetitive, compulsive-like behaviors in mice. J Vis Exp. 2013(82):50978.

65. Bankhead P, Loughrey MB, Fernandez JA, Dombrowski Y, McArt DG, Dunne PD, et al. QuPath: Open source software for digital pathology image analysis. Sci Rep. 2017;7(1):16878.

66. Schindelin J, Arganda-Carreras I, Frise E, Kaynig V, Longair M, Pietzsch T, et al. Fiji: an open-source platform for biological-image analysis. Nat Methods. 2012;9(7):676–82.

67. Lee FHF, Lai TKY, Su P, and Liu F. Altered cortical Cytoarchitecture in the Fmr1 knockout mouse. Mol Brain. 2019;12(1):56.

68. Becker LA, Huang B, Bieri G, Ma R, Knowles DA, Jafar-Nejad P, et al. Therapeutic reduction of ataxin-2 extends lifespan and reduces pathology in TDP-43 mice. Nature. 2017;544(7650):367–71.

69. Yamashita N, Morita A, Uchida Y, Nakamura F, Usui H, Ohshima T, et al. Regulation of spine development by semaphorin3A through cyclin-dependent kinase 5 phosphorylation of collapsin response mediator protein 1. J Neurosci. 2007;27(46):12546–54.

70. Shen S, Park JW, Lu ZX, Lin L, Henry MD, Wu YN, et al. rMATS: robust and flexible detection of differential alternative splicing from replicate RNA-Seq data. Proc Natl Acad Sci U S A. 2014;111(51):E5593–601.

71. Pletikos M, Sousa AM, Sedmak G, Meyer KA, Zhu Y, Cheng F, et al. Temporal specification and bilaterality of human neocortical topographic gene expression. Neuron. 2014;81(2):321–32.

72. Johnson MB, Kawasawa YI, Mason CE, Krsnik Z, Coppola G, Bogdanovic D, et al. Functional and evolutionary insights into human brain development through global transcriptome analysis. Neuron. 2009;62(4):494–509.

